# Dosage amplification dictates oncogenic regulation by the *NKX2-1* lineage factor in lung adenocarcinoma

**DOI:** 10.1101/2023.10.26.563996

**Authors:** John L. Pulice, Matthew Meyerson

## Abstract

Amplified oncogene expression is a critical and widespread driver event in cancer, yet our understanding of how amplification-mediated elevated dosage mediates oncogenic regulation is limited. Here, we find that the most significant focal amplification event in lung adenocarcinoma (LUAD) targets a lineage super-enhancer near the *NKX2-1* lineage transcription factor. The *NKX2-1* super-enhancer is targeted by focal and co-amplification with *NKX2-1*, and activation or repression controls *NKX2-1* expression. We find that *NKX2-1* is a widespread dependency in LUAD cell lines, where NKX2-1 pioneers enhancer accessibility to drive a lineage addicted state in LUAD, and NKX2-1 confers persistence to EGFR inhibitors. Notably, we find that oncogenic NKX2-1 regulation requires expression above a minimum dosage threshold—NKX2-1 dosage below this threshold is insufficient for cell viability, enhancer remodeling, and TKI persistence. Our data suggest that copy-number amplification can be a gain-of-function alteration, wherein amplification elevates oncogene expression above a critical dosage required for oncogenic regulation and cancer cell survival.

Highlights
- The most significant amplification event in LUAD targets a lineage super-enhancer that controls expression of the *NKX2-1* lineage oncogene.
- *NKX2-1* is a dosage dependency in most NKX2-1(+) LUAD cell lines
- NKX2-1 remodels lineage enhancer accessibility to drive a lineage addicted state and confer persistence to EGFR targeted therapy
- NKX2-1 oncogenic regulation requires a minimum oncogenic dosage, which dictates NKX2-1 regulation of enhancer remodeling, TKI persistence, and cancer cell viability

## INTRODUCTION

Somatic copy number alterations (SCNAs), which include amplification or deletion of focal^1^ or chromosome arm-level^2^ genomic regions, are critical and defining driver events in cancer^3^. Amplification events target a gene and/or enhancer region to amplify regulation in *cis* to drive oncogenic overexpression of the endogenous target gene^4^, and oncogene amplifications commonly harbor co-amplification of regulatory enhancers^5^. We previously reported recurrent focal enhancer amplifications as non-coding drivers of oncogene activation pan-cancer^6^—validated focal enhancer amplification events recurrently target surprisingly few critical oncogenic transcription factors (TFs) including *MYC*^6-9^*, KLF5*^10^, and *AR*^11-13^. Notably, *MYC*^6-9^ and *KLF5*^10^ are regulated by tissue-specific focal enhancer amplifications in multiple cancer types, suggesting focal enhancer amplification is a critical mechanism of activation for oncogenes such as c-MYC. The total incidence and importance of focal enhancer amplifications across cancer genomes is yet to be understood.

Amplification can target a wild-type (eg. *MYC*)^14^ or mutated (*EGFRvIII*)^15^ gene, thereby elevating gene expression to drive an oncogenic role in cancer. However, most studies of oncogenic TFs have used knockouts, overexpression to a single dosage, and/or genome-wide characterization of endogenous binding, without functional characterization of the role of dosage for an oncogenic TF. For the c-MYC oncogene in Burkitt’s lymphoma, a linear model of transcriptional amplification was identified^14^, however this result was later found to be due to a specific role for c-MYC as a selective transcriptional activator of metabolic processes like ribosome biogenesis^16^. For the SOX9 lineage factor in a craniofacial developmental context, a buffered model of dosage regulation was found, where most changes in chromatin accessibility occurred from 0-25% of wild-type SOX9 dosage^17^. To date, the role of dosage modulation in oncogene regulation for amplified oncogenic TFs has not been characterized, to the best of our knowledge.

Lung cancer remains the largest driver of cancer deaths in the United States^18^. Lung adenocarcinoma (LUAD) is the most common subtype of non-small cell lung cancer (NSCLC), with LUAD comprising 40% of total lung cancer cases, and with a five year survival rate of only 28%^19^. We and others identified the lineage transcription factor gene *NKX2-1* (also known as thyroid transcription factor 1 or TTF-1/TITF-1) as the most significantly amplified gene in LUAD^20-24^. NKX2-1 is a homeobox transcription factor that defines specification and development of the lung, thyroid, and ventral forebrain^25,26^, and is required for alveolar cell identity and survival in mature AT1 and AT2 cells^27,28^. *NKX2-1* is amplified at the earliest stages of LUAD^29^ and across all sites of multi-focal LUAD tumors^30^, suggesting a critical role in LUAD initiation and maintenance. *NKX2-1* has the single strongest pan-cancer correlation between expression and motif accessibility across all transcription factors^31^, suggesting a direct and critical role for NKX2-1 in controlling the accessible genome in primary human tumors.

Our understanding of NKX2-1 is complicated by paradoxical roles in lung adenocarcinoma— *Nkx2-1* loss is seen during metastatic progression in the *Kras^LSL-G12D/+^;Trp53^flox/flox^* (KP) LUAD mouse model^32^. Conversely, *Nkx2-1* is required for *EGFR*-mutant LUAD mouse models, with heterozygous deletion significantly decreasing tumor burden and size^33^. *NKX2-1* is a tumor suppressor subject to recurrent loss-of-function mutations in invasive mucinous adenocarcinoma (IMA)^33,34^, a rare LUAD subtype that exhibits hallmark loss of *NKX2-1* and alveolar identity^32-34^, and the KP mouse model reflects this specific LUAD subtype^33^, suggesting that a tumor suppressor role for *Nkx2-1* / *NKX2-1* is restricted to specific genetic and developmental contexts. In mouse organoid models of LUAD initiation, *Nkx2-1* is downregulated by *Kras^G12D^* induction, however down-regulation of *NKX2-1* is not seen in human organoids upon expression of *KRAS^G12D^*^35^. Despite clear genetic evidence of an oncogenic role for amplified *NKX2-1* in most (ie. non-mucinous) human LUADs, functional analyses of this oncogenic role have been limited to a few target genes or sites^36-38^, with little known about how *NKX2-1* amplification drives oncogenic regulation by this lineage-defining transcription factor.

Here, we find that the most significant focal amplification event in lung adenocarcinoma targets a lineage super-enhancer near *NKX2-1*. We demonstrate that the *NKX2-1* super-enhancer bidirectionally controls *NKX2-1* levels in LUAD, and map the transcriptional activity of the *NKX2-1* super-enhancer to individual enhancers and transcription factor binding sites. We find that *NKX2-1* is a broad dependency in LUAD, where modulation of *NKX2-1* dosage to sub-amplified levels is sufficient to confer a therapeutic vulnerability. Mechanistically, we find that NKX2-1 controls lineage enhancer accessibility to regulate an alveolar differentiation state that is hallmark to most human LUAD tumors, and that NKX2-1 drives a persistent state to targeted therapies to *EGFR*-mutant LUADs. Importantly, we find that a minimum dosage threshold controls the oncogenic regulation of *NKX2-1*—sub-amplified *NKX2-1* dosage is insufficient to mediate enhancer accessibility, TKI persistence, or cancer cell viability. Our data shows that dosage modulation through copy-number amplification drives gain-of-function regulation by NKX2-1, and suggests that copy-number changes to wild-type proteins can be a neomorphic driver in cancer.

## RESULTS

### The most significant focal amplification in LUAD targets a non-coding region near *NKX2-1*

We assembled and analyzed a cohort of genome-wide copy number profiles for human LUAD tumors, totaling 1,422 tumor/normal DNA pairs profiled by either single nucleotide polymorphism (SNP) array hybridization (Campbell+Weir, n=1021)^20,39^, or by whole-genome sequencing (WGS), (CPTAC+APOLLO, n=421)^40,41^. Amplification at chr14q13.3 near *NKX2-1* is the most significant focal somatic amplification event identified in these analyses (**Fig. 1a**), consistent with published studies^20-24^. The chr14q13.3 focal amplification peak called by GISTIC 2.0 does not overlap the *NKX2-1* coding region, comprising a 116 kb noncoding region 206 kb centromeric to the *NKX2-1* gene body (hg38: chr14:36195623-36311566; **Fig. 1b, S1a**). The chr14q13.3 amplification peak exhibits copy-number gain in 242/1021 (23.7%) of LUAD tumors by SNP array (**Fig. 1c**), and is amplified at a higher rate and average copy number than the *NKX2-1* gene (238/1021, 23.3%) (**Fig. 1c, S1a**).

**Figure 1.**
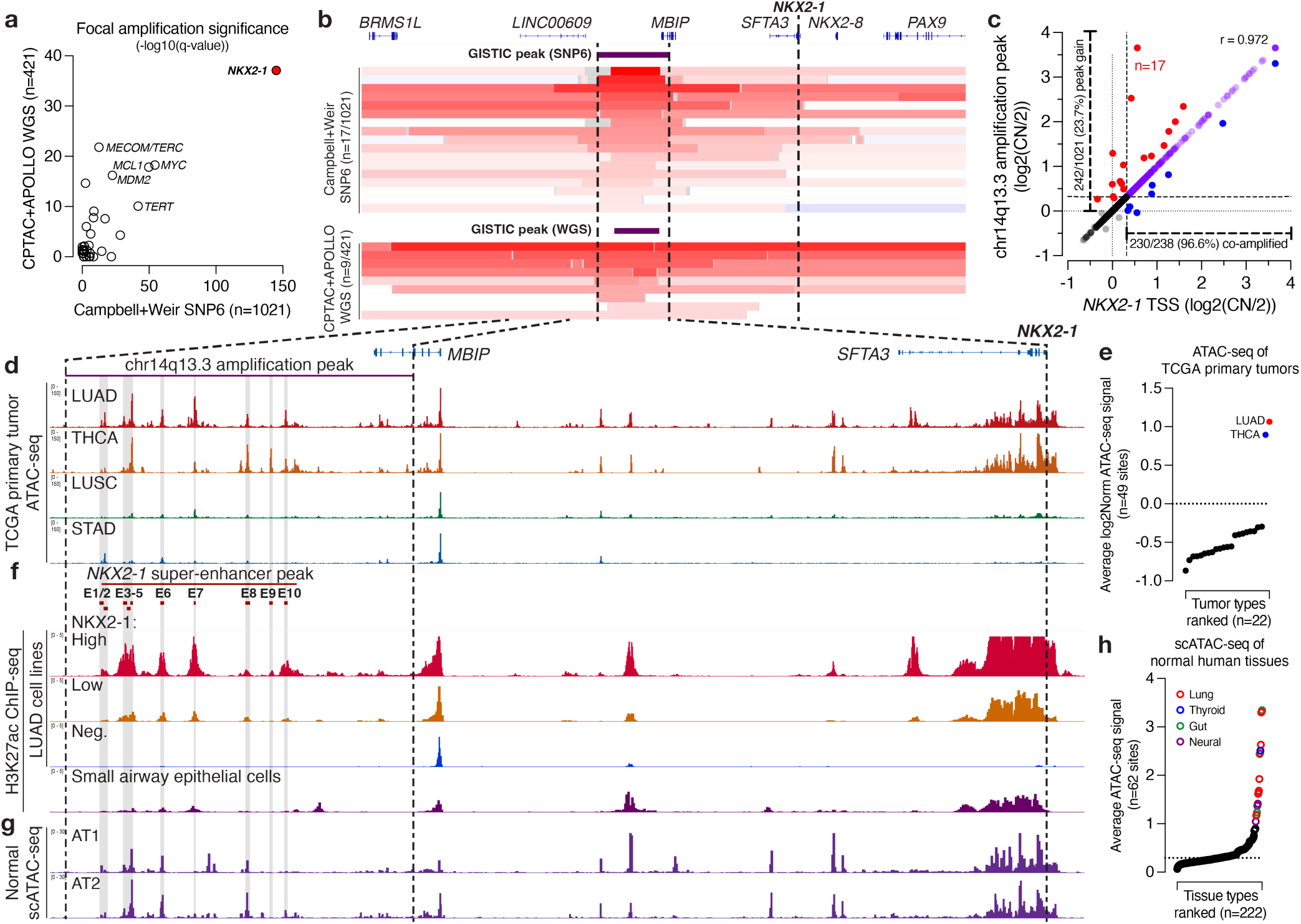
The most significant focal amplification peak in LUAD targets a lineage super-enhancer near *NKX2-1*. **(A)** Significance of focal amplification events in SNP6 (n=1021) and WGS (n=421) copy number profiles of lung adeno-carcinoma (LUAD) tumors. **(B)** Copy number profiles and GISTIC peak calls for chr14q13.3 focally amplified LUAD tumors by (top) SNP6 (n=17/1021) and (bottom) WGS (n=9/421). **(C)** Plot of copy number at the *NKX2-1* transcription start site (TSS) vs. the chr14q13.3 amplification peak in LUAD tumors (n=1021). **(D)** Average ATAC-seq chromatin accessibility across primary tumor types. THCA, thyroid carcinoma; LUAD, lung squamous cell carcinoma; STAD, stomach adenocarcinoma. **(E)** Average ATAC-seq accessibility^43^ within the chr14q13.3 amplification peak for n=22 TCGA primary tumor types, ranked. LUAD (red) and THCA (blue) tumors labeled. **(F)** Average H3K27ac ChIP-seq occupancy in LUAD and lung cell lines by NKX2-1 expression (high, n=10; low, n=3; negative, n=17). SAEC, small airway epithelial cells. Union super-enhancer region (*NKX2-1* SE), with 10 candidate enhancers, is shown. **(G)** Merged scATAC-seq accessibility profiles for distal lung cell types. AT1, alveolar type 1; AT2, alveolar type 2. **(H)** Average scATAC-seq signal^44^ across the chr14q13.3 amplification peak across n=222 annotated cell types, ranked. Lung (red), gut (green), thyroid (blue), and neural (purple) cell types are labeled. See also Figure S1-2.

The chr14q13.3 amplification peak is specifically targeted by focal copy-number gains exceeding the *NKX2-1* gene body in 17/1021 (1.7%) of LUAD tumors by SNP6 and 9/421 (2.1%) by WGS (**Fig. 1b-c, S1b**), as well as in 2/71 (2.8%) of LUAD cell lines profiled by SNP array in the Cancer Cell Line Encyclopedia (CCLE)^42^ (**Fig. S1c-d**). Of *NKX2-1* gene-amplified tumors (238/1021; 23%), the chr14q13.3 amplification peak is co-amplified in 230/238 (96%) of tumors (**Fig. 1c**), with specific co-amplification of the two regions (**Fig. S1e**), suggesting this non-coding region is a critical and specific part of the *NKX2-1* amplicon in gene-amplified tumors. Structural variant (SV) analysis of WGS of LUAD tumors shows that the chr14q13.3 amplification peak is amplified by tandem duplication (**Fig. S1f**).

The chr14q13.3 amplification peak falls within the same topologically associated domain (TAD) as both *NKX2-1* and the *MBIP* gene (**Fig. S1g**). *NKX2-1* expression is lineage restricted while *MBIP* is constitutively expressed (**Fig. S1h**), which suggests that *NKX2-1* is the regulatory target of chr14q13.3 non-coding amplification. *NKX2-1* expression is significantly higher in chr14q13.3 focally amplified LUAD tumors than in samples of normal lung tissue, similar to *NKX2-1* expression levels in tumors harboring high-level co-amplification of the *NKX2-1* gene and chr14q13.3 peak (**Fig. S1i**). These data identify recurrent amplification of a non-coding region near *NKX2-1* as the most significant amplification event in LUAD.

### Focal amplification targets a *NKX2-1* lineage super-enhancer

We posited that the chr14q13.3 amplification peak near *NKX2-1* targets a region containing enhancer elements that could regulate *NKX2-1* expression. Through analysis of epigenomic data from LUAD primary tumors and cell lines, we find that the chr14q13.3 focal amplification peak harbors a lineage super-enhancer. We identify specific and robust accessibility at the chr14q13.3 amplification peak in LUAD and thyroid carcinoma (THCA) tumors with hallmark *NKX2-1* expression using assay of transposase accessible chromatin sequencing (ATAC-seq) from The Cancer Genome Atlas (TCGA)^31^ (**Fig. 1d-e, S2a-b**). Chromatin immunoprecipitation followed by sequencing (ChIP-seq) for histone H3 lysine 27 acetylation (H3K27ac) in 30 LUAD cell lines^44^ identifies multiple enhancers within the chr14q13.3 amplification peak that are specifically marked by H3K27ac in *NKX2-1*-expressing cell lines (**Fig. 1f**). The chr14q13.3 amplification peak harbors at least one ‘super-enhancer’ in 8/13 NKX2-1(+) LUAD cell lines, including 8/10 *NKX2-1* high-expressing LUAD cell lines, as identified from H3K27ac ChIP-seq data using standard methods^45,46^ (**Fig. 1f, S2c**). These analyses demonstrate that the most significant amplification event in LUAD targets a super-enhancer near *NKX2-1*. We therefore denote this region as the *NKX2-1* super-enhancer (*NKX2-1* SE).

Lastly, we sought to determine the tissue specificity of the *NKX2-1* SE using ATAC-seq from primary human tissues^47^. The *NKX2-1* SE contains accessible lineage enhancer elements identified, including multiple lung cell types including alveolar type 1 and 2 (AT1/2), ciliated, and club cells (**Fig. 1g-h, S2d**). In addition, the *NKX2-1* super-enhancer is accessible in multiple thyroid, gut, and neural cell types (**Fig. 1g-h, S2d**). Our data find that the *NKX2-1* SE is a lineage super-enhancer that is co-opted through amplification to drive oncogenic *NKX2-1* expression in LUAD.

In addition to lung adenocarcinoma, analysis of germline studies provides evidence for roles of the *NKX2-1* SE in other diseases. The *NKX2-1* SE harbors multiple SNP associations identified by genome-wide association studies (GWAS), including a thyroid cancer SNP in E10 (rs116909374^48^) and a lung function FEV1/FVC SNP in E3-5 (rs10132289^49^) (**Fig. S2e**). Benign Hereditary Chorea (BHC), also known as Brain Lung Thyroid Syndrome, is a developmental disorder that is defined by hallmark heterozygous loss of *NKX2-1*^50^. Focal deletions upstream of *NKX2-1* have been identified in BHC patients^51,52^—these deletions directly target the *NKX2-1* SE (**Fig. S2e**). Collectively, these data suggest that the *NKX2-1* SE may play a broader role in disease and cancer, as a risk allele for thyroid cancer as well as a critical regulatory region in lung and thyroid development.

### The *NKX2-1* SE controls *NKX2-1* expression through 3 active enhancer elements

We selected four cell lines for interrogation of *NKX2-1* and the *NKX2-1* SE: one that harbors focal amplification of the *NKX2-1* SE (HCC364), and three that harbor co-amplification of the *NKX2-1* SE and *NKX2-1* and have been previously used for study of NKX2-1: NCI-H2087^36,37,53^, NCI- H358^21,36,54^, and NCI-H441^38,53-55^ (**Fig. S3a-c**). We identified a 65 kb union super-enhancer region (hg38: chr14:36207754-36272803) within the chr14q13.3 amplification peak, and within this identified ten candidate enhancer regions (**Fig. 1f, 2a**). We identify three regions, E3-5, E6 and E7, with the hallmarks of active enhancer elements as marked by H3K27ac/H3K4me1 deposition, in all 4 cell lines we assayed by ChIP-seq (**Fig. 2a**). E1-2 and E8-10 are occupied by the hallmarks of poised enhancers (H3K4me1+/ H3K27ac–), in at least 2 of 4 cell lines (**Fig. 2a**), suggesting additional enhancers that could regulate *NKX2-1*.

**Figure 2.**
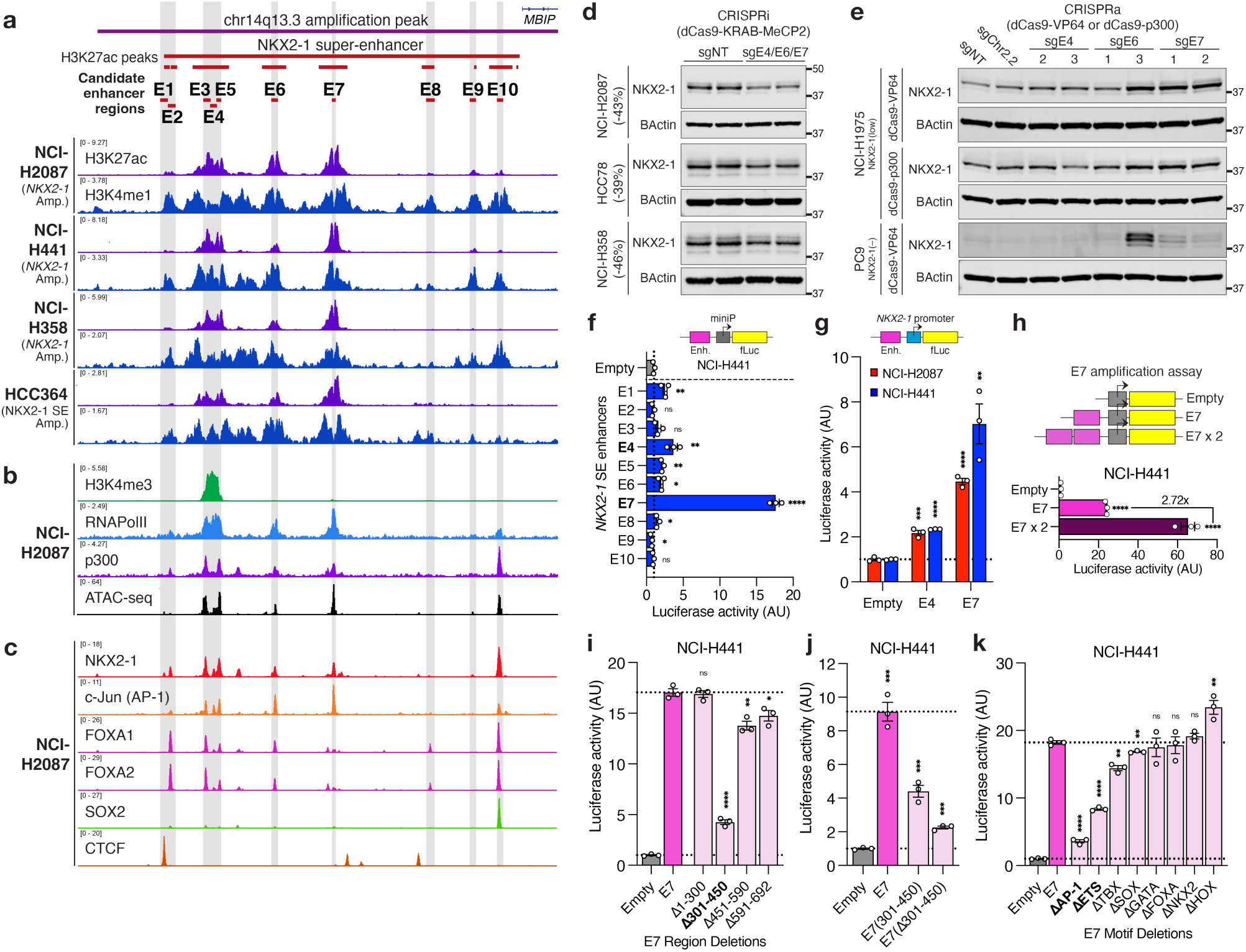
A focally-amplified super-enhancer controls *NKX2-1* expression. **(A)** ChIP-seq occupancy tracks for H3K27ac and H3K4me1 in 4 NKX2-1(+) LUAD cell lines. Union super-enhancer region (*NKX2-1* SE), with 10 candidate enhancers, is shown. **(B)** ChIP-seq occupancy tracks for H3K4me3, RNAPolII and p300, as well as ATAC-seq accessibility, in NCI- H2087 cells. **(C)** ChIP-seq occupancy tracks for NKX2-1, c-Jun, FOXA1/2, SOX2, and CTCF transcription factors in NCI- H2087 cells. **(D)** Immunoblot of NKX2-1 upon repression of the NKX2-1 SE, n=2 biological replicates. **(E)** Immunoblot of NKX2-1 upon activation of the NKX2-1 SE in NKX2-1(low) or NKX2-1(–) cells. **(f-g).** Luciferase enhancer activity for enhancers in the *NKX2-1* super-enhancer using a **(F)** miniP or **(G)** *NKX2-1* promoter reporter. **(H)**. Luciferase enhancer activity of a duplicated E7 enhancer shows >2x activity. **(I-K)**. Luciferase enhancer activity of E7 **(I)** region deletions, **(J)** minimal fragments, or **(K)** motif deletions. **(F-K)** Luciferase activity calculated relative to empty vector, error bars indicate mean±SEM, individual points labeled, n=3 biological replicates, two-tailed t-test, *p<0.05, **p<0.01, ***p<0.001, ****p<0.0001. Significance calculated against **(G-H,J)** empty vector control or **(I,K)** E7 full length. See also Figure S3-4.

We sought to further characterize the binding of regulators and transcription factors to the *NKX2-1* SE. We identify binding of transcriptional regulators including RNA polymerase II (RNAPolII), and p300 to E3-5, E6 and E7 (**Fig. 2b**), confirming these as likely transcribed enhancers. We profiled 6 transcription factors implicated in lung and LUAD by ChIP-seq, and observe varied patterns of transcription factor binding to the *NKX2-1* SE, including enhancer-specific binding such as SOX2 to E10 (**Fig. 2c**). Only NKX2-1 and AP-1 bind to all accessible enhancers (**Fig. 2c**), suggesting that a combination of factors control the *NKX2-1* SE, with NKX2-1 and AP-1 as broad regulators in cooperation with specific regulators at individual enhancers. These data suggest that three active enhancer regions, E3-5, E6, and E7, are likely to define the regulatory activity of the *NKX2-1* SE in LUAD.

To examine the impact of the *NKX2-1* SE on *NKX2-1* expression, we used CRISPR interference (CRISPRi) and CRISPR activation (CRISPRa) to modulate the *NKX2-1* SE. Using a validated dCas9-KRAB-MeCP2^56^ repressor for CRISPRi (**Fig. S3d**), we simultaneously expressed guides targeting the three active enhancers within the *NKX2-1* SE (E3-5, E6, and E7) using a co-expression cassette (U6- sgE7;H1-sgE6;7SK-sgE4) (**Fig. S3e-g**). Repression of the *NKX2-1* SE led to a 39%-46% decrease in NKX2-1 (**Fig. 2d**). These data show that the *NKX2-1* SE is required for endogenous *NKX2-1* expression in NKX2-1(+) LUAD cells.

We noted that a subset of NKX2-1(+) LUAD cell lines express low levels of *NKX2-1* (**Fig. S3h**), distinct from most NKX2-1(+) LUAD cell lines which express high levels of *NKX2-1*. We used *NKX2-1* low-expressing cell lines to model activation of *NKX2-1* dosage from low levels in a putative cell-of-origin to high levels upon amplification in LUAD. We wanted to determine if activation of the *NKX2-1* SE is sufficient to upregulate *NKX2-1*. NKX2-1(low) LUAD cell lines harbor low H3K27ac at the *NKX2-1* SE (**Fig. S3i** and **1f**, above). Using a validated dCas9-VP64^57^ or dCas9-p300^58^ activator for CRISPRa (**Fig. S3j**), we find that CRISPRa targeting to E6 or E7 markedly upregulated NKX2-1, whereas active guides targeting E4 have little effect on NKX2-1 levels (**Fig. 2e, S3k-l**). In *NKX2-1* non-expressing PC-9 cells, we find that CRISPRa targeting E6 or E7 also activates NKX2-1 expression (**Fig. 2e, bottom**). In summary, the activity of the *NKX2-1* SE controls *NKX2-1* expression in LUAD, explaining the hallmark nature of the *NKX2-1* SE to *NKX2-1* amplification and activation in LUAD (**Fig. S3m**).

### Mapping the transcriptional activity of the *NKX2-1* SE to base pair resolution

We wanted to map the specific regions and motifs required for the transcriptional activity of the *NKX2-1* SE. However, the dynamic range of endogenous manipulation by CRISPRi, combined with the technical limitations of manipulating an amplified enhancer region by CRISPR-Cas9, led us to use a luciferase reporter to assay the enhancers of the *NKX2-1* SE. We performed luciferase reporter assays to determine the transcriptional activity of the 10 enhancers of the *NKX2-1* SE.

We find that E4 and E7 are significant and robust (>2.5 fold) enhancers of transcription in all 4 NKX2-1(+) LUAD cell lines assayed (**Fig. 2f, S4a**). E7 is a notably robust transcriptional enhancer, similar to the strong *MYC* E3 enhancer we previously characterized^6^ (**Fig. S4a**). Both E4 and E7 activate transcription at a distance using a reporter where the enhancer is downstream from the miniP- fLuc cassette (**Fig. S4b**), and E7 activates transcription independent of sequence orientation (**Fig. S4c**). We find that E4 and E7 also activate transcription from the *NKX2-1* promoter sequence in a luciferase reporter assay (**Fig. 2g, S4d-e**), demonstrating direct compatibility of the *NKX2-1* promoter sequence with the enhancers in the *NKX2-1* SE. We find a duplicated (“amplified”) E7 drives super-additive transcriptional activity, at 2.7-4.2x the activity of a single E7 allele (**Fig. 2h, S4f**). These results demonstrate that two key enhancers of the *NKX2-1* SE, E4 and E7, are robust transcriptional enhancer elements.

We undertook fine mapping of the E7 enhancer by stepwise or region deletions (**Fig. S4g**). We identified a 150 bp region, 301-450bp, responsible for E7 activity—E7(Δ301-450) resulted in a 75% decrease in enhancer activity (**Fig. 2i, S4h-i**). Notably, a minimal E7(501-692) fragment still retained 39% of full length E7 activity, suggesting multiple hubs critical for E7 activity (**Fig. S4h**). The complementary E7(301-450) fragment recapitulated between 40-66% of full length E7 activity, consistently higher than E7(Δ301-450), which retained only 14-25% of full length E7 activity (**Fig. 2j, S4j**). Deletion of the putative binding motifs for 8 ubiquitous or lineage-restricted transcription factors from the E7 sequence identifies the AP-1 (74-80% decrease) and ETS (37-54% decrease) as critical for E7 activity (**Fig. 2k, S4k**). Endogenously, E7 specifically shows strong binding of AP-1 as compared to other factors (**Fig. 2c** above) which suggests collaboration of NKX2-1, AP-1, and other factors to control a critical enhancer within the *NKX2-1* SE.

### NKX2-1 is a lineage dependency required in LUAD

Given that *NKX2-1* is a target of both gene and enhancer amplifications, we sought to determine if *NKX2-1* is critical for driving cancer cell proliferation as an oncogene. We evaluated dependency for *NKX2-1* in genome-scale shRNA and CRISPR-Cas9 screens^59^, and find that *NKX2-1* is a dependency across most NKX2-1(+) but not NKX2-1(-) LUAD cell lines^59^ (**Fig. 3a-b, S5a-b**). Notably, *NKX2-1* is an essential gene (Chronos ≤ -0.7)^59^ in the majority of NKX2-1(+) LUAD cell lines (median Chronos score = -0.73) (**Fig. 3b**). *NKX2-1* dependency occurs in direct relationship to *NKX2-1* expression, with cell lines expressing highest levels of *NKX2-1* exhibiting strongest *NKX2-1* dependency (**Fig. 3a, S5a**). These results demonstrate that *NKX2-1* is a critical dependency across most NKX2-1(+) LUAD cell lines.

**Figure 3.**
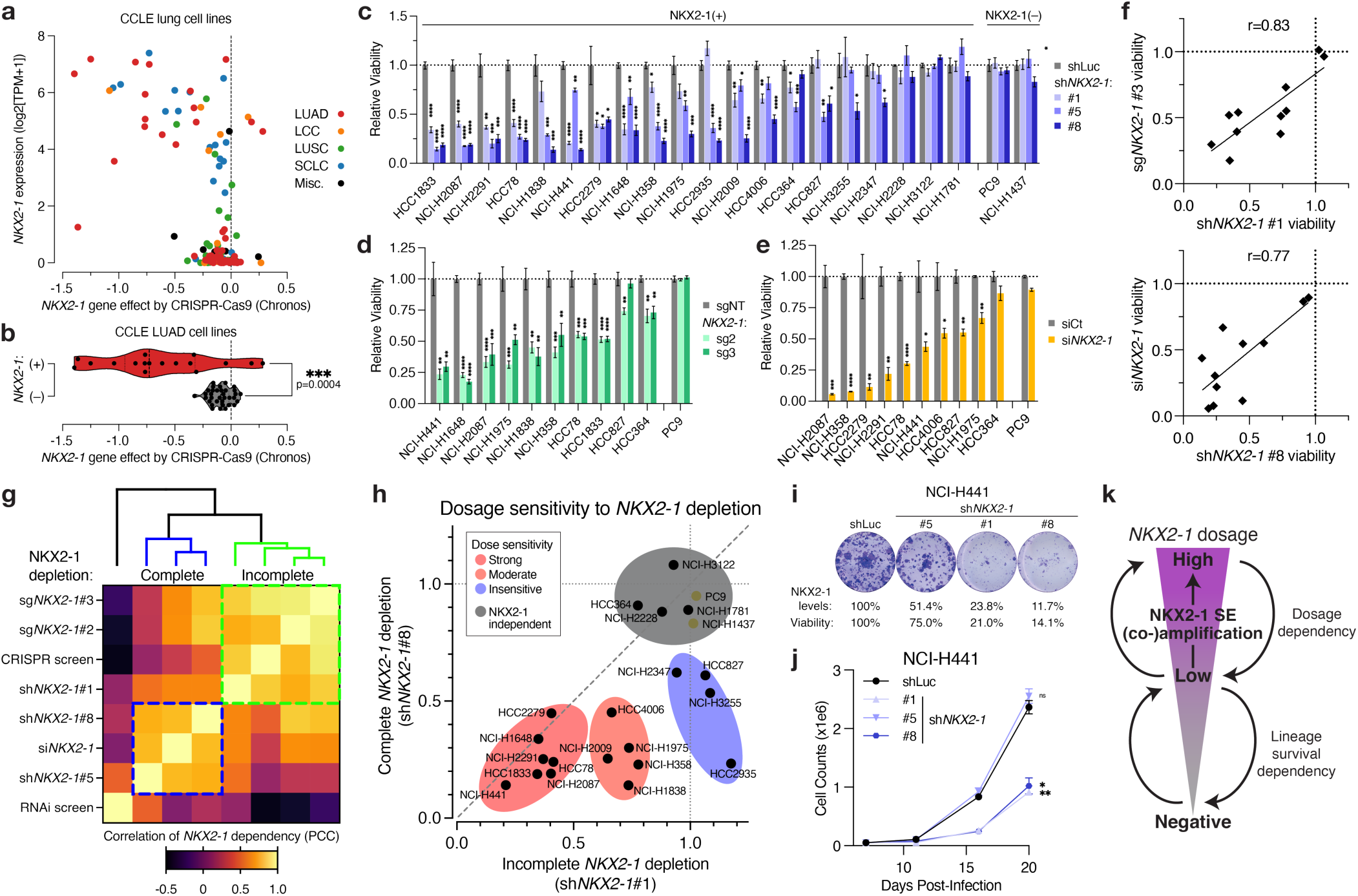
*NKX2-1* is a dosage dependency across LUAD cell lines. **(A)** Plot of *NKX2-1* gene effect by CRISPR-Cas9 vs. expression for CCLE lung cell lines. **(B)** Violin plot of *NKX2-1* gene effect by CRISPR-Cas9 in NKX2-1(+) (red) or NKX2-1(–) (grey) CCLE LUAD cell lines. Two-tailed t-test, ***p=0.0004. **(C)** Relative viability by clonogenic assay of NKX2-1(+) (n=20) and NKX2-1(–) (n=2) cell lines upon shRNA knockdown of *NKX2-1*. n=4 technical replicates. **(D)** As in (C), for CRISPR-Cas9-mediated bulk knockout of *NKX2-1*. n=4 technical replicates. **(E)** As in (C), for siRNA-mediated knockdown of *NKX2-1*. n=3 biological replicates. **(F)** Plot of NKX2-1 dependency in assays of (top) incomplete or (bottom) complete loss. **(G)** Heatmap of correlation between *NKX2-1* dependency across assays. Assays of incomplete (green) or complete (blue) NKX2-1 loss cluster together. **(H)** Plot of NKX2-1 dependency upon incomplete (sh*NKX2-1*#1) vs. complete (sh*NKX2-1*#8) knockdown. Cells with strong (red), moderate (orange), or no (blue) dosage sensitivity to incomplete *NKX2-1* loss are indicated. **(I)** Crystal violet staining of clonogenic assays upon sh*NKX2-1* dosage modulation. Percent NKX2-1 suppression and cell viability loss are indicated. **(J)** Proliferation curves upon sh*NKX2-1* dosage modulation, n=2 technical replicates. **(K)** Schematic for dosage and survival dependency upon amplified NKX2-1 expression in LUAD. **(B-E,J)** Two-tailed t-test against control, *p<0.05, **p<0.01, ***p<0.001, ****p<0.0001. Error bars = Mean±SEM. See also Figure S5.

We selected 20 NKX2-1(+) and 2 NKX2-1(–) LUAD cell lines to characterize *NKX2-1* dependency, spanning dosage levels, copy number status, primary/metastatic origin, and oncogenic driver alterations (**Fig. S5c**). We used three primary shRNAs (#1, #5, #8) to suppress *NKX2-1*—shRNA #1 exhibited consistent but incomplete suppression of *NKX2-1* levels, whereas shRNA #8 consistently displayed near complete *NKX2-1* depletion by immunoblot analysis, and shRNA #5 varied by context (**Fig. S5d**).

We identified a widespread requirement for *NKX2-1* in NKX2-1(+) LUAD cell lines upon shRNA transduction—17/20 cell lines exhibited a statistically significant viability defect upon ≥1 shRNA, and 15/20 cell lines exhibited a statistically significant viability defect for ≥2 shRNAs (**Fig. 3c, S5e**). Half (10/20) of NKX2-1(+) cell lines are strongly dependent on *NKX2-1*, as shown by a >50% viability decrease for ≥2 shRNAs targeting *NKX2-1* (**Fig. 3c, S5e**). Our results confirm and extend prior assays of individual cell lines^20-22^. We validated our results by CRISPR-Cas9 bulk knockout transduction (**Fig. 3d, S5f-g**) and by siRNA transfection (**Fig. 3e, S5h-j**), each of which led to decreased cell viability that aligned with shRNA results (**Fig. 3c**). We confirm our observed viability defects extend to cell proliferation (**Fig. S5k**) and reflect cell cycle arrest in NCI-H441 cells (**Fig. S5l**). Our results demonstrate a widespread and robust requirement for NKX2-1 for LUAD cell survival.

### *NKX2-1* is a dosage dependency for cancer cell viability

We found significant variability in the viability defects we have identified in vitro for *NKX2-1*. For example, we identified much stronger viability defects using siRNA (**Fig. 3e**), which caused near total ablation of NKX2-1 protein levels (**Fig. S5h**), in comparison to the more modest effects of CRISPR- Cas9 bulk knockout (**Fig. 3d**), for which the multiple copies of *NKX2-1* confer a higher likelihood for ≥1 alleles repairing in frame to restore wild-type protein. By comparison, distinct *NKX2-1* shRNAs strongly recapitulated each of these findings—we see strong concordance between sh#1 and sg#3 (r=0.82), as well as between sh#8 and siRNA depletion (**Fig. 3f**).

By Pearson correlation coefficient (PCC) and dendrogram clustering, we identify two clusters of *NKX2-1* viability defects, associated with: (A) incomplete loss of NKX2-1 by sg*NKX2-1*#2 or #3, sh*NKX2-1*#1, and CRISPR-Cas9 screen dependency, and (B) near-complete NKX2-1 ablation by sh*NKX2-1*#5 or #8 and si*NKX2-1*, with RNAi screen dependency as an out group (**Fig. 3g**). We used two shRNAs (#1 and #8) as proxies for incomplete (#1) and complete (#8) loss of *NKX2-1*. Most (16/20) NKX2-1(+) LUAD cell lines exhibit cell lethality upon complete loss of *NKX2-1* by sh#8 (**Fig. 3h**). Among these, cell lines may either be insensitive (4/16, yellow), moderately sensitive (5/16, orange), or highly sensitive (7/16, red) to incomplete NKX2-1 loss (**Fig. 3h**). The majority (12/20) of NKX2-1(+) LUAD cell lines assayed are sensitive to incomplete loss of *NKX2-1* (**Fig. 3h**).

Our data suggest that dosage controls endogenous *NKX2-1* dependency, wherein LUAD cells can be sensitive to incomplete or complete NKX2-1 loss for loss of viability in vitro. For example, in NCI-H441 cells, moderate depletion of NKX2-1 by sh#5 did not affect cell clonogenicity or proliferation, but stronger (sh#1) or near-complete (sh#8) depletion conferred cell lethality in vitro (**Fig. 3i-j, S5k**). Our data find that NKX2-1(+) LUAD cells specifically require elevated *NKX2-1* dosage for their survival, and demonstrates that modulation of *NKX2-1* to sub-oncogenic levels affects the survival of most NKX2-1(+) LUAD cells.

### NKX2-1 remodels lineage enhancer accessibility in LUAD

Given the broad requirement for *NKX2-1* for cancer cell viability, we sought to characterize mechanistically how amplified *NKX2-1* expression regulates the LUAD epigenome and transcriptome. We identify robust NKX2-1 binding across the regulatory epigenome of 4 NKX2-1(+) LUAD cell lines, where we identify widespread (75,781-90,772 sites) and concordant (32,633 sites shared) binding of NKX2-1 by ChIP-seq (**Fig. S6a-c**). NKX2-1 binding is biased toward TSS-distal sites (>10 kb to nearest TSS) (**Fig. S6d**) where we find central enrichment of the canonical CACT[C/T] NKX2/3 family motif (**Fig. S6e**).

We selected NCI-H441 and NCI-H2087 for mechanistic interrogation of NKX2-1 function—these cell lines are the closest cell line models of primarily LUAD tumors and are closer to human tumors than most LUAD patient-derived xenograft models (**Fig. S6f-g**). We performed ATAC-seq of NCI-H2087 and NCI-H441 cells upon NKX2-1 depletion by validated shRNAs (see **Fig. 2**), and found widespread and concordant changes in genome accessibility (**Fig. 4a-c, S6h-i**), with specific loss of accessibility at NKX2-1-bound enhancer sites (**Fig. 4d, S6j**). Loss of NKX2-1-mediated enhancer accessibility can be seen at loci including *HOPX* and *SFTPB* loci (**Fig. S6k**). Our data suggest a direct role for NKX2-1 driving enhancer accessibility, distinct from other transcription factors studied in cancer epigenome regulation^60,61^. NKX2-1 has not been previously suggested to be a pioneer factor^62^—our data suggest NKX2-1 as a factor that can pioneer de novo chromatin accessibility at target sites to regulate gene expression.

**Figure 4.**
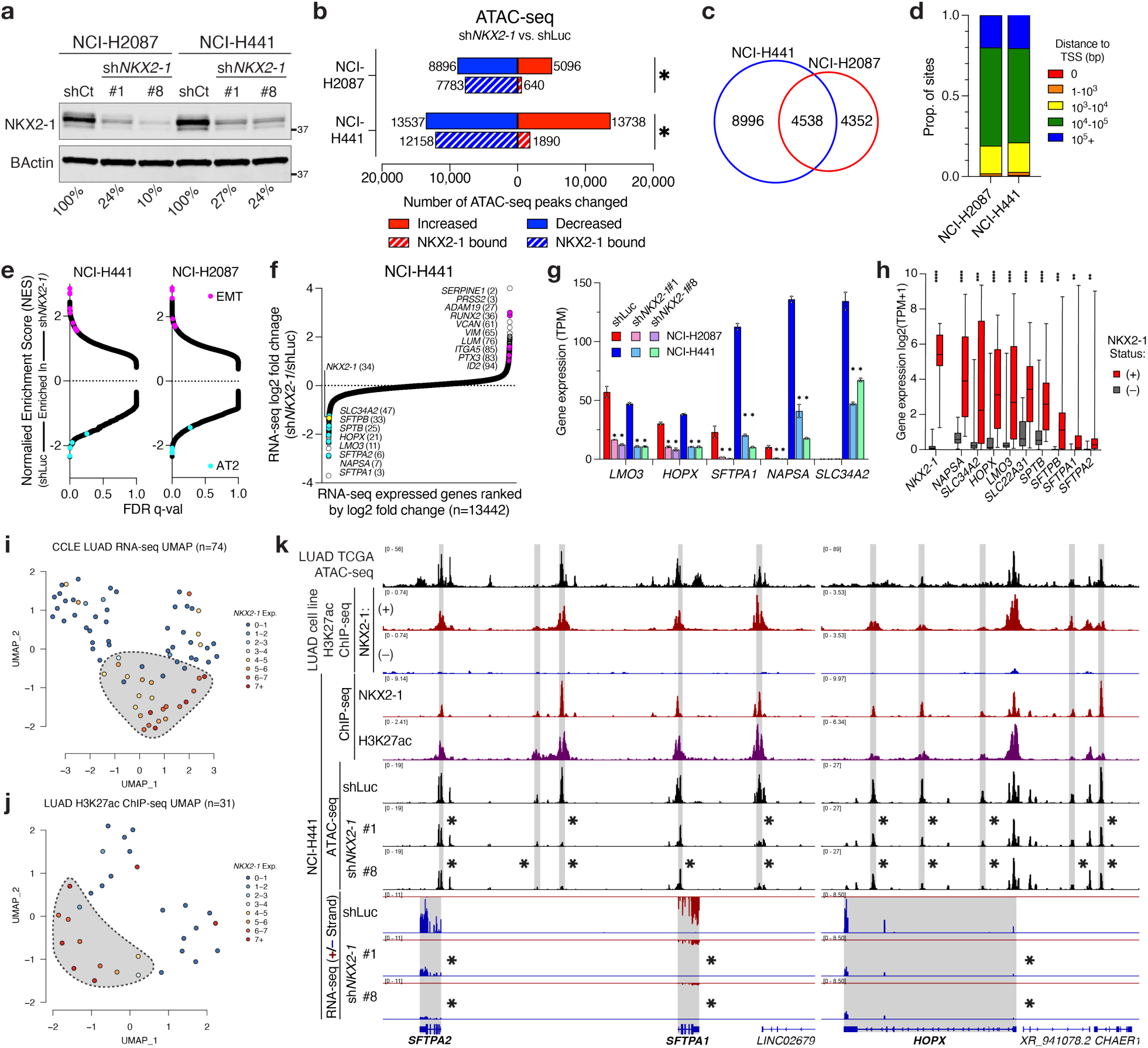
NKX2-1 drives alveolar differentiation through remodeling enhancer accessibility. **(A)** Immunoblot of NKX2-1 upon shRNA-mediated suppression of *NKX2-1*, or a luciferase-targeting control. Percent change in densitometry indicated. **(B)** ATAC-seq sites with increased (red) or decreased (blue) accessibility upon *NKX2-1* knockdown, sites overlapping NKX2-1 ChIP-seq peaks are indicated (hashed). Fisher exact test *p<2.2e-16. **(C)** Venn diagram of ATAC-seq peaks with decreased accessibility upon *NKX2-1* knockdown in NCI-H2087 or NCI-H441 cells. **(D)** Distance to transcription start site (TSS) for peaks with decreased ATAC-seq accessibility upon *NKX2-1* knockdown in NCI-H2087 or NCI-H441 cells. **(E)** GSEA analysis of ranked RNA-seq changes across gene sets. EMT, epithelial-mesenchymal transition; AT2, alveolar type 2. **(F)** Ranked RNA-seq fold-change for all expressed genes, AT2 (cyan) and EMT (magenta) genes labeled. **g.** Bar graph of normalized RNA-seq expression for AT2 genes regulated by NKX2-1. Error bars = mean±SEM, n=2 biological replicates, individual values labeled. * significant by DESeq2 (FC ≥ 1.5, Bonferroni adj. p-value ≤ 1e-3). TPM, transcripts per million. **(H)** Boxplot of NKX2-1 regulated AT2 genes across all CCLE LUAD cell lines (n=74), separated by *NKX2-1* expression as (+) (n=30, red) or (–) (n=44, grey). Two-tailed t-test, *p<0.05, **p<0.01, ***p<0.001, ****p<0.0001). Whiskers = min-max. **(I-J)** UMAP clustering of **(I)** RNA-seq or **(J)** H3K27ac ChIP-seq from LUAD cell lines. Cell lines are colored by *NKX2-1* expression, NKX2-1(+) subpopulation indicated. **(K)** NKX2-1 drives alveolar gene expression through enhancer remodeling. (top) TCGA LUAD ATAC-seq and H3K27ac ChIP-seq (middle) NKX2-1 and H3K27ac ChIP-seq (bottom) ATAC-seq and stranded RNA-seq in the indicated conditions. * indicates significantly downregulated ATAC-seq peak or RNA-seq gene by DESeq2. See also Figure S6-7.

### NKX2-1 drives alveolar differentiation through enhancer regulation

We next sought to determine how NKX2-1-mediated enhancer accessibility regulates gene expression in LUAD. NKX2-1 depletion leads to outlier downregulation of alveolar differentiation gene sets and upregulation of epithelial-mesenchymal transition (EMT) gene sets by RNA sequencing (RNA- seq) (**Fig. 4e, S7a-e**). The downregulation of alveolar differentiation genes includes genes encoding key surfactant proteins (*SFTPA1, SFTPA2, SFTPB)*, an AT1/2 transcription factor (*HOPX*) and key markers of differentiation (*NAPSA, AQP4, SLC34A2, LMO3*) among the top 40 strongest downregulated genes (**Fig. 4f, S7f-g**). Our data shows that regulation of alveolar differentiation is the strongest direct target of NKX2-1 regulation in LUAD cells.

Notably, while some NKX2-1 target genes are consistently regulated by NKX2-1 across cell lines assayed (eg. *LMO3, HOPX*), modulation of other targets by NKX2-1 depletion depends on the baseline expression and differentiation state of each cell line. That is, NKX2-1 depletion modulates some genes from high to low expression in NCI-H441 and low to absent in NCI-H2087 (eg. *SFTPA1*, *NAPSA*) and other targets regulated in NCI-H441 are not expressed at baseline in NCI-H2087 (eg. *SLC34A2*) (**Fig. 4g, S7g**). This reflects the relative alveolar differentiation state of each cell line (**Fig. S7h**)—NKX2-1 depletion leads to de-differentiation in a manner dictated by the initial differentiation state of each LUAD cell line. These results identify a key role for NKX2-1 in controlling differentiation state and EMT in LUAD.

We asked whether NKX2-1 controls a distinct transcriptional and epigenome state in LUAD. Across CCLE LUAD cell lines, alveolar differentiation genes are specifically expressed in NKX2-1(+) LUAD cell lines (**Fig. 4h, S7h**). UMAP dimensionality reduction shows that *NKX2-1* expression identifies a subgroup of NKX2-1(+) cell lines at both the transcriptomic (**Fig. 4i**) and epigenomic (**Fig. 4j**) level, distinct from 2 predominantly NKX2-1(–) populations. Our data identify a specific alveolar differentiation state at the RNA and chromatin level that is specific to *NKX2-1* expression.

We extended our analyses of a *NKX2-1* gene expression signature to primary LUAD tumors. *NKX2-1* is expressed in most primary and metastatic human LUADs, reflecting a central role for *NKX2- 1* across human tumors (**Fig. S7i-k**). NKX2-1(+) and NKX2-1(–) primary tumors also separate at the transcriptome level in TCGA LUAD RNA-seq (**Fig. S7l**), though NKX2-1(–) primary or metastatic LUAD tumors are quite rare (**Fig. S7i-k**). Note that *NKX2-1* expression is 1.9x higher in the highest amplified tumors compared to normal lung or unamplified tumors (**Fig. S1i**), and *NKX2-1* expression in LUAD tumors is similar to that in normal alveolar type 2 cells in scRNA-seq data (**Fig. S7j**). Our data suggest that *NKX2-1* amplification may play two roles: (1) to increase *NKX2-1* expression in *NKX2-1* low expressing lung cells (Clara cells, club cells), or (2) to maintain oncogenic dosage and counteract oncogene-mediated downregulation in *NKX2-1* high expressing lung cells (AT1/2).

Combining our analyses of LUAD tumors and cell lines, we identify an *NKX2-1*-correlated gene signature in CCLE (n=259) and TCGA (n=262) LUADs by RNA-seq, and find significant overlap with genes significantly downregulated by *NKX2-1* depletion (n=383) in NCI-H441 cells (**Fig. S7m**). 17 genes were identified in all 3 gene sets, as correlated to *NKX2-1* expression in cell lines and tumors and anti-correlated to *NKX2-1* depletion, including key alveolar differentiation target genes such as *SFTPA1/2, SFTPB, SLC34A2, NAPSA* and *LMO3* (**Fig. S7m**). These data identify an NKX2-1(+) population of LUAD cell lines and tumors, with distinct transcriptome and epigenome features, where NKX2-1 determines the expression of cell fate genes.

Overall, we can see the role for NKX2-1 in defining the LUAD transcriptome and epigenome at multiple loci, including *SFTPA1/2, LMO3,* and *HOPX* (**Fig. 4k, S7n**). At these sites, we observe ATAC-seq accessibility in primary LUADs, histone H3K27 acetylation specifically in NKX2-1(+) cell lines, and NKX2-1 binding by ChIP-seq (**Fig. 4k, S7n, top**). NKX2-1 depletion leads to a significant decrease in ATAC-seq accessibility at nearby enhancers and promoters, and a marked loss of gene expression by RNA-seq (**Fig. 4k, S7n, bottom**). Loss of linage identity is seen in cell morphology changes upon NKX2-1 suppression—cells lose cell-cell adhesion and change growth morphology in vitro (**Fig. S7o**). Our data suggest that NKX2-1 controls lineage commitment in LUAD cell lines and tumors through enhancer remodeling, driving a lineage addicted state that we suggest is a hallmark feature of non-mucinous NKX2-1(+) human LUAD tumors.

### NKX2-1 enhancer remodeling is governed by a minimum dosage threshold

*NKX2-1* is expressed in the putative cells of origin for lung adenocarcinoma—therefore, we sought to understand how dosage modulation of *NKX2-1* to near-amplified levels alters its regulation in LUAD. We used NCI-H1975 (NKX2-1 low) and PC9 (NKX2-1(–)) cells to modulate NKX2-1 dosage with overexpression of *NKX2-1* on constitutive promoters to low (hPGK) and high (EF1α) dosage (**Fig. 5a- b**). Exogenous NKX2-1(high) expression resembles levels seen in NCI-H441 cells (**Fig. 5b**). We observe widespread changes in the accessible genome by ATAC-seq upon exogenous NKX2-1(high) expression in PC9 (15,207 sites changed) and NCI-H1975 (35,564 sites) cells (**Fig. 5c, S8a**), with sites of gained accessibility promoter-distal and enriching the NKX2-1 motif (**Fig. S8b-c**).

**Figure 5.**
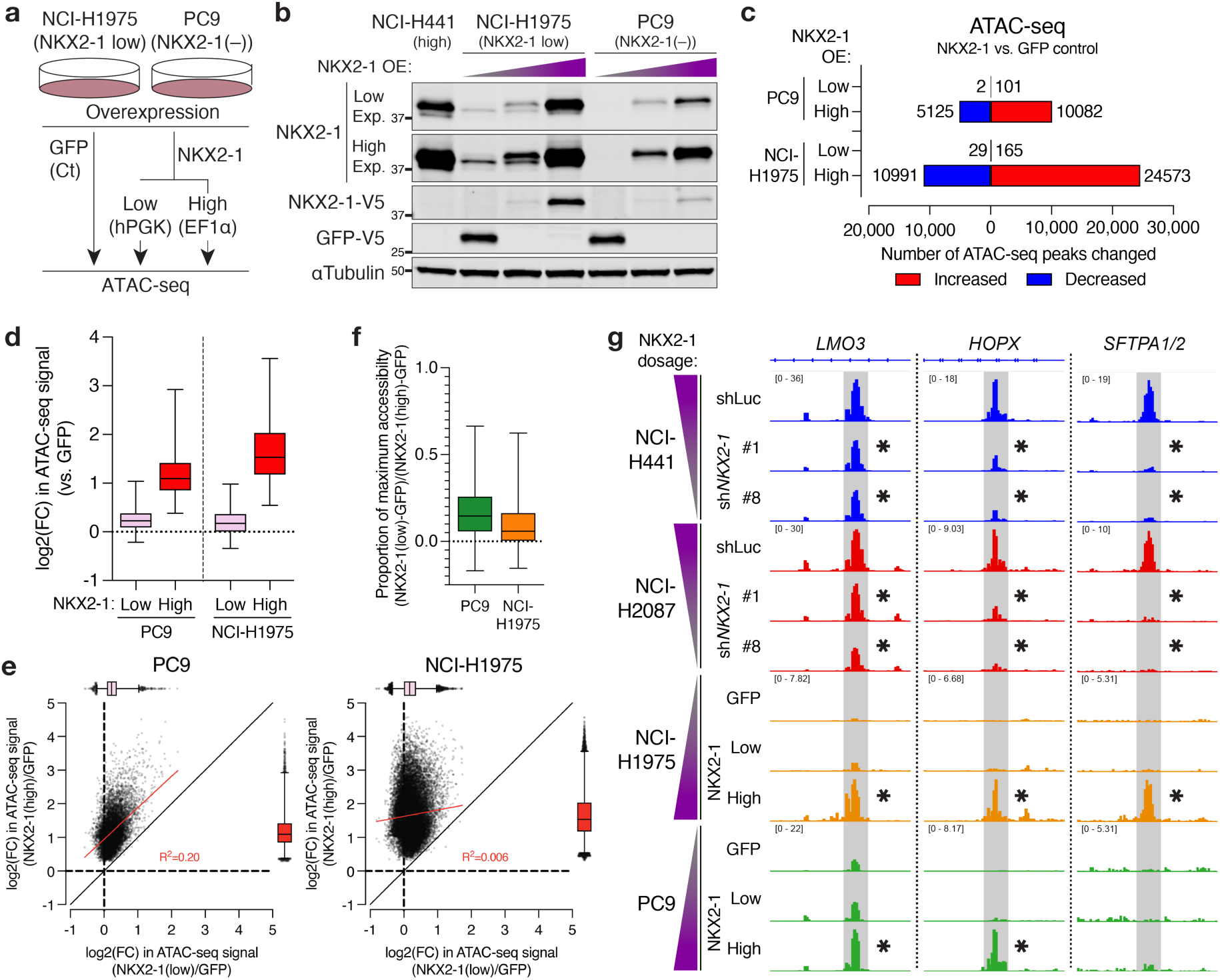
NKX2-1 remodeling of chromatin accessibility requires a critical dosage expression. **(A)** Schematic for NKX2-1 dosage modulation by constitutive overexpression. **(B)** Immunoblot analysis of NKX2-1, GFP, and V5 levels upon dosage modulation of NKX2-1. **(C)** ATAC-seq sites with increased (red) or decreased (blue) accessibility upon NKX2-1 dosage modulation. **(D)** Boxplot of log2(fold change) in chromatin accessibility at sites of increased accessibility upon NKX2-1(high) expression in PC9 or NCI-H1975 cells, as compared to GFP control. Whiskers = 1-99 percentile. **(E)** Plot of log2(fold change) in ATAC-seq accessibility induced by either NKX2-1(low) (x-axis) or NKX2-1(high) (y- axis) overexpression, as compared to GFP control, for all sites of increased accessibility by NKX2-1(high) in PC9 (n=10082) or NCI-H1975 (n=24573) cells. A boxplot of values for each condition is included alongside scatterplot, whiskers=1 to 99 percentile. **(F)** Boxplot of the proportion of maximum ATAC-seq accessibility induced by NKX2-1(low) (GFP control = 0, NKX2-1(high) = 1) at NKX2-1(high) upregulated sites in PC9 or NCI-H1975 cells. Whiskers = 1-99 percentile. **g.** Genome accessibility upon NKX2-1 modulation by *NKX2-1* suppression or overexpression in indicated cell lines. * indicates sites significantly regulated with NKX2-1 modulation by DESeq2 (FC ≥ 1.25, Bonferroni adj. p-value ≤ 1e-3). See also Figure S8.

Surprisingly, NKX2-1(low) expression has little effect on the accessible genome, with few sites of changed accessibility in PC9 (103 sites) and NCI-H1975 (194 sites) cells, respectively (**Fig. 5c**). The failure of NKX2-1(low) overexpression to induce genome-wide accessibility is notable and surprising, especially as PC9 cells lack endogenous NKX2-1 (**Fig. 5b**) that could mask the effects of overexpression. NKX2-1(low) induces modest changes in accessibility at NKX2-1(high) induced sites in PC9 (17% increase) and NCI-H1975 (12.8%) cells (**Fig. 5d-e**), however these increases are fractional when compared to accessibility changes induced by NKX2-1(high) in PC9 (14.9% of NKX2-1(high) accessibility) or NCI-H1975 (5.9% of NKX2-1(high) accessibility) (**Fig. 5f**). Few sites in PC9 (4.0%, 402/10082) or NCI-H1975 (2.6%, 636/24573), achieve at least 50% of maximal accessibility in NKX2- 1(low) conditions (**Fig. 5f**). We find significant overlap between sites controlled by NKX2-1 upon knockdown and overexpression (**Fig. S8d**), and can see this concordant regulation at genes such as *LMO3, HOPX, SFTPB,* and *SPTB* (**Fig. 5g, S8e**). Our data suggest this dosage requirement governs NKX2-1 regulation at endogenous target sites.

Prior studies of c-MYC^14^ or SOX9^17^ suggest either linear or buffered responses to dosage modulation. In the case of SOX9, most changes in chromatin accessibility occur in the increase from 0% to 25% of wild-type SOX9 dosage. Our data suggests a distinct mechanism of dosage regulation— modulation of NKX2-1 dosage from 0% to 20% of highest NKX2-1 expression has little effect genome-wide on chromatin accessibility, whereas increasing NKX2-1 dosage from 20% to 100% drives widespread rearrangement of the accessible genome. Our data suggest that a dosage threshold model governs NKX2-1 regulation, whereby a physiologically relevant minimum NKX2-1 dosage is required for NKX2-1 to remodel chromatin accessibility. Our data suggests that *NKX2-1* amplification in LUAD may be a mechanism to modulate NKX2-1 dosage to drive gain of function enhancer remodeling.

### NKX2-1 controls EGFR TKI persistence

Previous studies have found that alveolar lineage genes, and *NKX2-1* specifically, are upregulated by LUAD cells in a persistent state to EGFR tyrosine kinase inhibitors such as osimertinib^63^. *NKX2-1* expression and *EGFR* mutations are intrinsically linked in the terminal respiratory unit (TRU) subtype of LUAD^64^—NKX2-1(–) *EGFR* mutant LUAD tumors and cell lines are rare^65^ (**Fig. 6a, S9a-b).**

**Figure 6.**
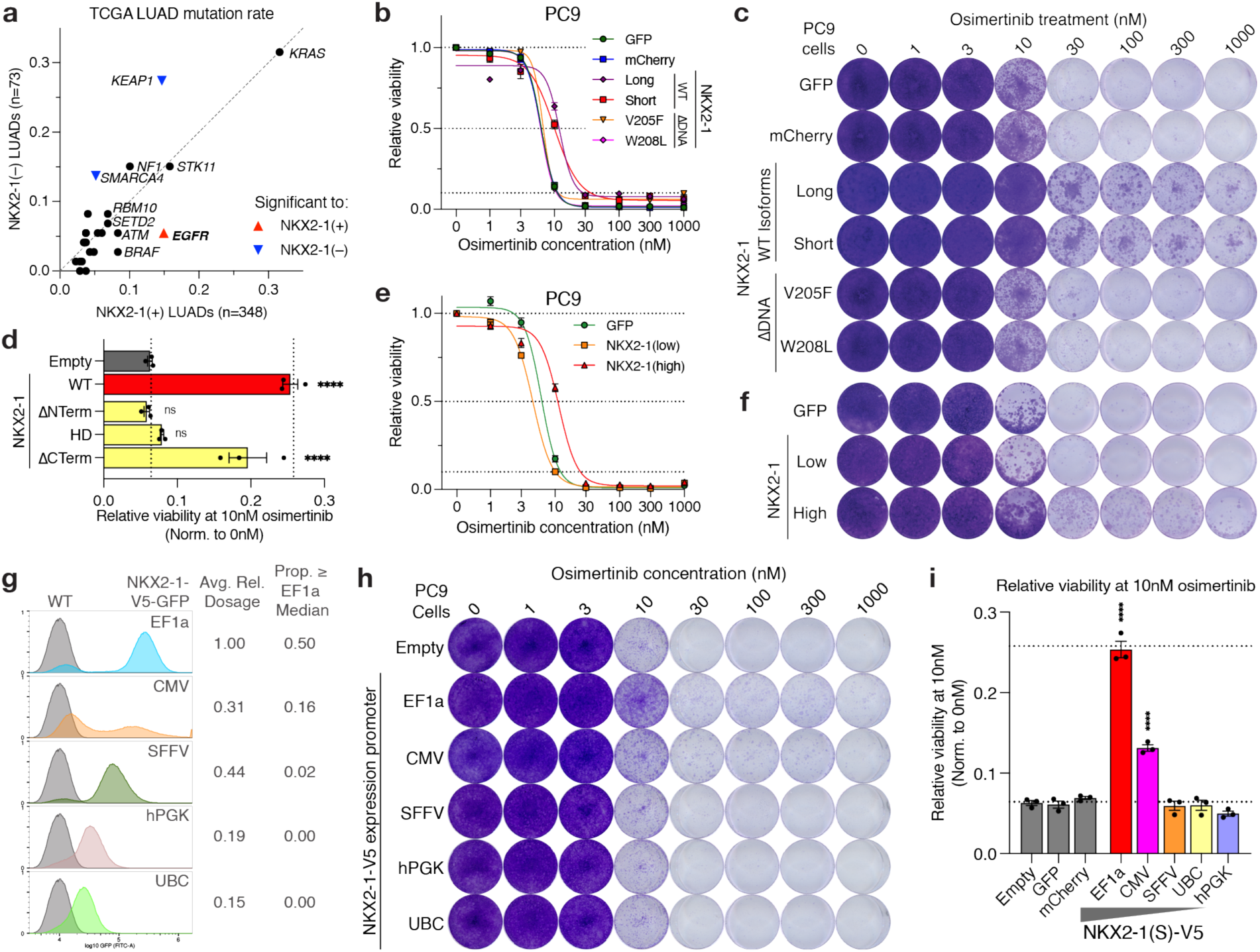
NKX2-1 drives EGFR TKI persistence in a dose-thresholded manner. **(A)** Mutation rate for top 25 significant driver alterations in TCGA LUAD tumors by NKX2-1(+) or NKX2-1(–) status. Significantly enriched mutations for NKX2-1(+) (red) or NKX2-1(–) (blue) are labeled. **(B)** Relative viability of PC9 cells in the indicated conditions upon osimertinib (EGFR tyrosine kinase inhibitor) treatment, normalized to 0nM control. n=3 biological replicates, error bars=mean±SEM. **(C)** Crystal violet staining of PC9 cells as in **(B)**, NKX2-1 increases the IC50 as well as cell persistence to osimertinib treatment. **(D)** Relative viability of PC9 cells in the indicated conditions at cytostatic 10nM osimertinib concentration. **(E)** Relative viability of PC9 cells, upon osimertinib treatment, n=4 biological replicates, error bars=mean±SEM. **(F)** Crystal violet staining of PC9 cells as in **(E)**, NKX2-1(high) expression is required for TKI persistence. **(G)** Flow cytometric analysis of PC9 cells expressing NKX2-1-V5-GFP from the indicated promoters, or naive control cells. Average expression relative to EF1a and proprotion of cells above median EF1a expression are indicated. **(H)** Crystal violet staining of PC9 cells expressing NKX2-1 on the indicated promoters, or empty vector control, upon osimertinib treatment. **(I)** Relative viability of PC9 cells in the indicated conditions at cytostatic 10nM osimertinib concentration, n=3 biological replicates, error bars = mean±SEM, individual points labeled, Two-tailed t-test vs controls, ****p<0.0001. See also Figure S9-10.

We find that NKX2-1 overexpression at high dosage in PC9 (*EGFR*-Exon19Δ, NKX2-1(–)) cells increases cell survival at both cytostatic (10nM) and cytotoxic (≥30nM) osimertinib doses, while expression of NKX2-1 harboring DNA-binding mutations V205F or W208L fails to increase survival (**Fig. 6b-c, S9c**). We find that the N-terminal domain of NKX2-1 is required for NKX2-1 to drive cell persistence (**Fig. 6d, S9d-f**). The closely related NKX2-4 protein also drives cell persistence to EGFR inhibition, while NKX2-5 is largely unable to do so (**Fig. S9d,g-h**). Our data show that NKX2-1 directly drives a persistent state to TKI inhibitors in PC9 cells, and map the precise domain requirements for NKX2-1 to drive TKI persistence.

Conversely, we find that strong depletion of NKX2-1 in HCC827 (*EGFR*-Exon19Δ, NKX2-1(+)) cells induces increased sensitivity to osimertinib at doses tolerated by control cells, and disrupts cell persistence (**Fig. S9i-j**). This synergistic cell killing is beyond the viability defects of NKX2-1 depletion, as evidenced by specific decreases in cell viability at ≤3nM (**Fig. S9k**). This effect is not seen with the weaker sh*NKX2-1*#1, suggesting only near-complete NKX2-1 depletion is able to mediate synergistic cell killing with osimertinib treatment in these cells. Together, our results demonstrate that NKX2-1 mediates EGFR TKI persistence in EGFR-mutant LUAD cells.

### Precise mapping of how NKX2-1 dosage controls TKI persistence

Notably, we find that NKX2-1 control of EGFR TKI survival is governed by the same dosage requirement we observed for genome accessibility—PC9 cells (as profiled by ATAC-seq, see **Fig. 5b**) expressing NKX2-1(low) fail to exhibit increased EGFR TKI survival mediated by NKX2-1(high) (**Fig. 6e-f, S10a-b**). Therefore, we sought to precisely map this dosage requirement for NKX2-1 to drive EGFR TKI survival. We overexpressed *NKX2-1-V5* as well as a GFP-fused *NKX2-1-V5-GFP* on five constitutive promoters (EF1a, CMV, SFFV, hPGK, UBC) to titrate NKX2-1 dosage in a precise and quantitative manner across single cells (**Fig. 6g, S10c-d**). 4 of 5 promoters exhibit consistent and stepwise dose-titrated expression as shown by flow cytometry, with highest expression induced by EF1a > CMV > SFFV > hPGK > UBC, except that CMV expression is variable across cells, with some cells expressing near-EF1a levels and some at near-background GFP levels (**Fig. 6g, S10e**). Immunoblot analysis of *NKX2-1-V5* expression affirms the relative average expression levels captured by flow cytometry (**Fig. S10f**).

We find that a dosage threshold governs NKX2-1 regulation of EGFR TKI persistence; SFFV-driven expression levels of NKX2-1 (44% of maximum EF1a-driven expression levels) fail to drive any significant persistent state that is driven by EF1a high expression (**Fig. 6h-i, S10g**). Notably, CMV-driven *NKX2-1* expression is able to mediate EGFR TKI survival, but at a lower fraction than cells with EF1a-driven *NKX2-1* expression, likely owing to a highly variable CMV-driven expression pattern (**Fig. 6h-i, S10e**). Using the EF1a median expression as our dose threshold, we find that the proportion of cells above this threshold directly correlates with the degree of TKI persistence observed (**Fig. S10h**).

Our data demonstrate that NKX2-1 mediates cell persistence to EGFR inhibition, and we map this precise dosage requirement for NKX2-1 to mediate survival to targeted therapy. We suggest that dosage modulation of NKX2-1 may target persistent LUAD disease upon EGFR inhibitor treatment, beyond the viability defects upon NKX2-1 dosage modulation we have identified herein.

## DISCUSSION

Oncogene amplification is a common and critical event in cancer^1,2^. Here, we identify amplification of a super-enhancer of *NKX2-1* as the most significant amplification in LUAD, and find that the *NKX2-1* SE dictates *NKX2-1* expression in LUAD cells. We identify *NKX2-1* as the master regulator of LUAD cell state—NKX2-1 remodels enhancer accessibility to drive an alveolar differentiated state, where NKX2-1 is required across LUAD cell models for cell viability, and NKX2-1 controls EGFR oncogenic signaling to mediate a TKI persistent state. Importantly, we find that NKX2-1 oncogenic regulation is subject to a critical dosage requirement we precisely map—NKX2-1 dosage at sub-oncogenic levels fails to mediate chromatin accessibility, TKI persistence, or mediate cancer cell survival. Our data suggests that oncogene amplification is a gain-of-function driver alteration in cancer, wherein dosage modulation enables neomorphic regulation that sub-oncogenic levels fail to mediate.

Our laboratory and others previously identified focal enhancer amplifications as driving events in human cancer^6-13^—however, few events have been documented in cancer. We identify focal amplification of an *NKX2-1* super-enhancer as a novel driving oncogenic event in LUAD, and find that the *NKX2-1* SE is a hallmark of *NKX2-1* amplification in LUAD. Hereby, we add *NKX2-1* to the small class of oncogenes (*MYC, KLF5, AR*) validated to be activated by focal enhancer amplification to drive oncogenic expression. In addition, our data suggests that focal enhancer amplification is a specifically critical oncogenic process in LUAD through targeting both the *NKX2-1* and *MYC*^6^ oncogenes, similar to the outsized role of regulatory translocations in hematopoietic malignancies^66^. The *NKX2-1* SE also harbors risk variants for thyroid and lung cancer and developmental disorders^48,49^, including focal deletion events^51,52^, suggesting a broader role for the *NKX2-1* SE in *NKX2-1* expressing tissues and tumors.

The role of *NKX2-1* in lung adenocarcinoma is complex—*Nkx2-1* is a tumor suppressor lost in LUAD mouse models^32^, however, *NKX2-1* is the most significantly amplified gene in human LUADs^39^ and is expressed in 85-90% of human LUADs^67^. Our data suggest that NKX2-1 is such a critical oncogenic driver in LUAD that multiple types of CNAs target *NKX2-1* and its SE to amplify *NKX2-1* expression to oncogenic levels, and its expression is required across most NKX2-1(+) LUAD cell lines, adding evidence to its oncogenic role in LUAD. Importantly, the role for *Nkx2-1* as a tumor suppressor in mouse LUADs is specific to *Kras*-mutant contexts^32,33^, *EGFR*-mutant mouse LUADs require *Nkx2-1* for tumorigenesis^33^. Our data suggest a fundamental distinction between human and mouse LUADs, wherein *NKX2-1* is a lineage oncogene in most human LUADs.

Importantly, we find that human LUADs are largely defined by lineage addiction driven by *NKX2-1*, suggesting a critical oncogenic role across most human LUADs. NKX2-1 drives a lineage addicted state in LUAD, remodeling lineage enhancers to regulate alveolar differentiation. We find that NKX2-1(+) LUAD models broadly require *NKX2-1* for oncogenic proliferation, substantiating a role for NKX2-1 in mediating cancer cell survival as an oncogene. Recently, alveolar differentiation was identified as an upregulated signature for persistence to targeted therapy^63^—here, we show that *NKX2- 1* controls this differentiation state to drive persistence to EGFR targeted therapies. Together, we identify lung adenocarcinoma as a cancer type defined by lineage addiction through the amplified *NKX2-1* lineage oncogene.

Oncogene amplification is a critical and defining driver alteration in many cancer types^3^, however little is known about how dosage modulation of a wild-type transcription factor drives oncogenic regulation in cancer. A previous study modulated SOX9 dosage in a developmental context, and identified largely buffered effects of SOX9 on chromatin accessibility—that is, most changes in chromatin accessibility occurred from 0-25% of wild-type SOX9 dosage^17^. Here, we present a distinct model of dosage modulation, wherein NKX2-1 oncogenic regulation is governed by a dosage threshold—modulation of NKX2-1 dosage from 0-25% of oncogenic expression levels has negligible effect on chromatin accessibility, whereas oncogenic dosage mediates widespread changes in chromatin accessibility. We find these results extend to EGFR inhibitor persistence and cancer cell dependency, wherein modulation of NKX2-1 to sub-amplified dosage is sufficient to confer a viability defect in most LUAD cell lines. Our data suggest that dosage control may differ between developmental and oncogenic contexts, such that dosage modulation by gene amplification can modulate dosage past a critical threshold required for gain-of-function regulation by an oncogene.

This study demonstrates how the analysis of non-coding cancer genome alterations coupled with epigenomic data can elucidate novel mechanisms of regulation that link cellular differentiation and oncogenesis, and a strong need for further genome-wide characterization of the cancer epigenome and SCNAs in cancer.

## Acknowledgments

We thank Andrew Cherniack, Douglas Wheeler, Brian Golbourn, Alfredo Valencia, Roodolph St. Pierre, Gabriel Sandoval, Xiaoyun Wu, Xiaoyang Zhang and members of the Meyerson Lab for helpful discussions and experimental protocols. We thank Leslie Gaffney for graphical design assistance. This work is supported by a grant from the National Cancer Institute to MM (R35CA197568) and an NCI Diversity Supplement to this grant, 3R35CA197568-09S1, that supports JLP. JLP was supported by an NDSEG Graduate Research Fellowship as well as a T32 training grant (5T32GM007226-43),

## Author Contributions

**J.L.P.** and **M.M.** conceived and designed the study, acquired funding, and wrote the manuscript. **J.L.P.** designed and conducted all experiments, and performed all data analyses and visualizations.

## Materials & Correspondence

Further information and requests for resources and reagents should be directed to and will be fulfilled by the lead contact, Matthew Meyerson

## STAR METHODS

### Cell Culture

A549, HCC2935, HCC4006, HCC827, NCI-H1437, NCI-H1648, NCI-H1781, NCI-H1838, NCI-H1975, NCI-H2009, NCI-H2087, NCI-H2228, NCI-H2291, NCI-H2347, NCI-H358, NCI-H441 and HEK293T cells were obtained from ATCC, HCC78 cells were obtained from DSMZ, HCC1833 and HCC2279 were obtained from KCLB, NCI-H3122 and NCI-H3255 were originally obtained from the Minna laboratory, PC9 cells were obtained from ICB, HCC364 cells were obtained from CCLE. All cell lines were identity validated by STR Profiling with PowerPlex16HS (Labcorp) at initial low-passage samples. All cell lines were confirmed mycoplasma negative using the PCR Mycoplasma Detection Kit (ABM). Cell lines were cultured in either DMEM + 10% Fetal Bovine Serum + 1X PenStrep (A549, HEK293T) or RPMI-1640 + 10% Fetal Bovine Serum + 1X PenStrep (all other cell lines).

### Tumor copy number data analyses

Segmented copy number data for Campbell et al. (2016)^39^ SNP6 data were obtained from cbioportal.org, copy number data for Weir et al. (2007) (https://portals.broadinstitute.org/tcga/home) were assembled from analysis in Beroukhim et al. (2010) and mapped to hg19 from hg18 using UCSC.

Copy number values for the NKX2-1 transcription start site (TSS) (hg19; chr14:36988761) and NKX2-1 SE (hg19; chr14:36707741) were obtained using segmented copy number values at the specific position. If no value was available at this exact position, the nearest value to the specified location was used. For SNP6 copy number data, a sample was classified as “*NKX2-1* amplified” if the relative SNP6 copy number was ≥2.5. Enhancer amplified samples were identified based on (*NKX2-1* SE copy number ≥ 0.1 + *NKX2-1* TSS copy number). Average CN profile for n=660 profiles was generated using R, averaging the CN value in each window for all samples that have a value for that region—regions not called for a tumor were ignored in these analyses. Heatmaps of segmented copy number were generated using IGV (v. 2.15.5).

For upstream/downstream co-amplification analyses, we identified regions equally upstream of the NKX2-1 SE (hg19: 36426721) or equally downstream of *NKX2-1* (hg19: 37269781) as the distance between the NKX2-1 SE and *NKX2-1*. Copy number analyses and co-amplification determination occurred as above.

Mutation data for Campbell et al. (2016) were acquired from cbioportal.org as above, segmented copy number profiles were separated based the presence of a *KRAS*, *EGFR*, or neither mutation. GISTIC2 (v. 2.0.23)^68^ was run on mutation-specific Campbell et al. copy number data set using genepattern.org.

Segmented copy number profiles of LUAD tumors from CPTAC (n=338)^40^ and APOLLO (n=83)^41^ were obtained from the GDC. GISTIC2 (v. 2.0.23) was run on the full CPTAC data set using genepattern.org, filtering out germline CNVs identified in hg38, with parameters (amplification threshold = 0.1, deletion threshold = 0.1, q-value = 0.25, remove X = no, cap value = 1.5, confidence peel = 0.90, focal length cutoff = 0.50, maximum sample segments = 2000, and arm peel = yes). Copy number profiles for CPTAC+APOLLO LUADs are previously ploidy assigned to the nearest whole copy number value, so we classified a sample as “*NKX2-1* amplified” if the copy number was ≥3. Copy number values for the *NKX2-1* TSS (hg38; chr14: 36519556) and NKX2-1 SE (hg38; chr14:36238535) were obtained using segmented copy number values at the specific position. If no value was available at this exact position, the nearest value to the specified location was used.

OncoSG^69^ GISTIC2 analyses of copy number by WES (n=305) were obtained from Supplementary Table 8 of that paper^69^. This applies to both overall analyses and to ancestry and smoking status specific analyses from the OncoSG study. LUAD never-smoker^70^ GISTIC2 analyses of copy number by WGS (n=232) were obtained from Supplementary Table 3.

Structural variant analyses for whole-genome sequencing of LUAD tumors was obtained from Gillette et al. (2020)^40^ or Carrot-Zhang et al. (2021)^71^. Breakpoints were obtained from this analysis of whole-genome sequencing, copy number values were used from publicly available WGS or SNP6 copy number profiles of these tumors.

### TCGA RNA-seq data analysis

RNA-seq expression for TCGA LUAD were acquired from the PanCanAtlas^72^—samples specific to Campbell et al. from the Imielinski et al. cohort did not have paired RNA-seq for downstream analyses. Samples were classified as no amplification (CN < 2.5), low amplification (2.5≤CN<4), and high amplification (CN≥4) based on segmented copy number at the *NKX2-1* TSS as above. Enhancer amplified samples were identified as above (NKX2-1 SE copy number ≥ 0.1 + *NKX2-1* TSS copy number), samples with gene amplification but also greater SE copy number were classified as enhancer amplified. TCGA LUAD tumors were classified as high expression at *NKX2-1* RSEM ≥ 4000, and low/absent expression at *NKX2-1* RSEM ≤ 1000. UMAP clustering analysis performed on the top 5% most variable genes using all LUAD tumor and normal lung RNA-seq data, using the R package umap.

An NKX2-1 signature was identified by correlating all genes with *NKX2-1* expression in TCGA LUAD tumors (n=518), and selecting genes with R ≥ 0.5 correlation with *NKX2-1*, yielding a n=262 gene set.

### TCGA ATAC-seq analysis

TCGA ATAC-seq tracks and normalized peak-specific ATAC-seq values were obtained from deposited Corces et al. (2018)^31^ analyses. For visualization, we used prepared normalized ATAC-seq bigwig tracks <u>from the GDC</u>. For simplicity, we used the T1 first technical replicate for each tumor for visualization. For ATAC-seq tracks by tumor type, we calculated the average ATAC-seq signal across all tumors of that tumor type as follows. We converted each bigwig to a bedgraph using bigWigToBedGraph (UCSC), calculated the average signal for each 100bp window using bedtools and awk: “bedtools unionbedg -I $file_list > | awk ‘OFS=”\t” {sum=0; for (col=4; col<=NF; col++) sum += $col; print $1, $2, $3, sum/(NF-4+1); } > comb_ATAC.bdg”, and then converted back to a bigwig using bedGraphToBigWig (UCSC).

To analyze the specificity of ATAC-seq signal in the NKX2-1 SE, we used published log2 normalized ATAC-seq signal^43^ for each peak called in the TCGA ATAC-seq cohort. We overlapped the Campbell et al. amplification region with the TCGA ATAC-seq peak calls, and identified 49 ATAC-seq peaks within the NKX2-1 SE. For each tumor, we then averaged the signal at all 49 of these peaks to identify the ATAC-seq signal within the NKX2-1 SE in each tumor. For tumor type scores, we then averaged the NKX2-1 SE ATAC-seq score for all tumors in each tumor type. ATAC-seq peaks corresponding to the E1-10 of the NKX2-1 SE were assigned using bedtools overlap.

### CCLE cell line analyses

CCLE cell line analyses relied on DepMap Public 22Q4 data files^42,59^. We relied on CCLE assignment of tumor (sub)type for identification and analysis of LUAD cell lines. Mutation, expression, and *NKX2-1* gene copy number values were obtained from current data releases for 22Q4. We denoted a cell line as NKX2-1 positive if it had log2(TPM+1) *NKX2-1* expression > 3, as this naturally bifurcated the LUAD cell lines. However, we manually annotated NCI-H1648 as NKX2-1(+) as western blot analysis confirmed NKX2-1 protein expression.

For analysis of the NKX2-1 SE copy number, the primary 22Q4 copy number profiles rely on a mix of WES and WGS profiles, with most LUAD cell line data derived from WES profiles which lack coverage of the NKX2-1 SE. Therefore, we used the legacy segmented CN values based on SNP6 profiles and identified the *NKX2-1* TSS and SE copy number as above using hg19 genome coordinates.

For Celligner analyses^59^, we used the Skyros DepMap portal to calculate distance values for all tumor models to lung adenocarcinoma primary tumors. We separated CCLE LUAD cell lines by NKX2-1 (+/–), and assigned rank values to cell lines based on distance values. We used a two-tailed t-test to compare the distance values to LUAD tumors for NKX2-1(+) and NKX2-1(–) LUAD cell lines.

An NKX2-1 signature was identified by correlating all genes with *NKX2-1* cell line expression in CCLE LUAD cell lines (n=74), and selecting genes with R ≥ 0.5 correlation with *NKX2-1*, yielding a n=259 gene set.

### Analysis of published LUAD cell line ChIP-seq data

We analyzed published ChIP-seq data for H3K27ac as well as input controls for 26 LUAD cell lines as well as small airway epithelial cells (SAEC) from Suzuki et al., 2014^44^, as well as for NCI-H3122 from Rusan et al., 2018^73^. We kept these initial analyses within this cohort due to the high quality of the data as well as ability to easily compare data from the same preparation. Data was downloaded from the Sequence Read Archive (SRA) using sratoolkit (v. 2.10.7), and aligned to the hg38 reference genome (GRCh38_noalt_as) using bowtie2 (v. 2.2.9) with parameters “-k 1” and converted to a final BAM file with samtools (v. 1.9). To generate genome-wide occupancy tracks, we used macs2 (v. 2.1.1) with parameters “-g hs -f BAM --nomodel -B --SPMR” to generate a per million normalized genome-wide track, and then converted this to a bigwig file using bedGraphToBigWig (UCSC). We called peaks on H3K27ac ChIP-seq using macs2 (v. 2.2.7.1) using a q-value cutoff of 0.001. Peak calls were filtered against ENCODE blacklist regions (ENCFF356LFX_blacklist.bed) using bedtools (v. 2.30.0). For super-enhancer analysis, we used the standard analysis pipeline ROSE (v 1.3.1), with H3K27ac ChIP-seq as well as input control at H3K27ac peak sites for that cell line.

For UMAP clustering analysis, we generated a union peak set across 27 published LUAD cell lines and 4 newly generated in this study. We mapped reads to these peaks using the R package featureCounts, normalized per million mapped reads, and quantile normalized this data using the R package limma. UMAP clustering analysis performed on the top 5% most variable sites using the R package umap.

### scATAC-seq data analyses

We analyzed published processed data for scATAC-seq of normal human tissues from Zhang et al. (2021)^47^. Bigwig tracks as well as cCRE bed files were obtained as listed in Key Resource Table of that paper^47^ and visualized using IGV (v. 2.15.5)^74^. We used the union set of cCRE elements identified in Zhang et al. and overlapped this with the Campbell et al. GISTIC peak to identify n=62 scATAC-seq peaks within the NKX2-1 SE.

We converted each bigwig file of scATAC-seq accessibility by cell type to a bedgraph file using bigWigToBedGraph, then overlapped the n=62 NKX2-1 SE CREs with this file using bedtools overlap. For the bedgraph ATAC-seq signal with our n=62 CREs, we multiplied the length of each bedgraph track value by the length of that value in base pairs, and divided by the total base pair length of the 62 CREs to get a normalized ATAC-seq signal for each tissue type. These values were ranked to determine the relative ATAC-seq signal by tissue type for the NKX2-1 SE.

### LUAD scRNA-seq analyses

We analyzed published normalized scRNA-seq of normal lung and LUAD tumors from (Kim et al., 2020)^75^. For normal/LUAD analyses by stage, we combined all cells from tumors of each annotated stage or lung cell type, and visualized the normalized *NKX2-1* expression using a truncated violin plot in Prism 9. We excluded tumor samples from individual tumor/site analyses if less than 50 tumor cells were sequenced in the final processed scRNA-seq data.

### TAD boundary analyses

TAD boundary called from Hi-C sequencing data were obtained from ENCODE^76^, specifically for IMR90 (ENCFF307RGV), A549 (ENCFF716CFF), NCIH460 (ENCFF822VBC), and Lung Lower Lobe (ENCFF525ISU). No LUAD cell lines expressing NKX2-1 were profiled by ENCODE, so we used primary lung with other LUAD cell lines to confirm TAD structure. Hi-C heatmap data from lung lower lobe (ENCFF896OFN)^76^ was visualized using Juicebox (v. 2.2.6)^77^. Accompanying NCI-H2087 CTCF ChIP-seq was generated in this study (above).

### *NKX2-1* and *MBIP* expression analyses

*NKX2-1* and *MBIP* normalized expression values were obtained from all TCGA PanCanAtlas (n=10332), CCLE Cell Lines (n=1565), and Gtex Human Tissues (n=54). Expression was visualized using a truncated violin plot using Prism 9.

### ChIP-seq sample preparation

Cells were processed according to prior protocols^78^. Specifically, cells were trypsinized, counted, and fixed in culture media at 1% formaldehyde at 37°C for 10min, quenched in 0.125M glycine for 5min at 37°C. Cells were pelleted, PBS washed, pelleted, resuspended and aliquoted in 10M increments in 1.5 mL Eppendorf tubes, pelleted again, aspirated, and stored at -80°C.

Fixed cells were thawed on ice and resuspended in Rinse Buffer 1 (50 mM HEPES pH 7.5, 140 mM NaCl. 1 mM EDTA, 10% Glycerol, 0.5% NP-40, 0.25% Triton-X + 1X Halt Protease/Phosphatease Inhibitor + 1 mM PMSF), 10min on ice, then pelleted 500g/5min. Buffer was aspirated, pellet was resuspended in Rinse Buffer 2 (10 mM Tris-HCl pH 7.5, 1 mM EDTA, 0.5 mM EGTA, 200 mM NaCl + 1X Halt Protease/Phosphatease Inhibitor + 1 mM PMSF), pelleted at 500g/5min. Buffer was aspirated, and cells were resuspended in 275µL of Covaris Shearing Buffer (10 mM Tris-HCl pH 8, 1 mM EDTA, 0.1% SDS + 1X Halt Protease/Phosphatease Inhibitor + 1 mM PMSF) per 10M fixed cells. Cells were aliquoted to an 8 tube microtube strip (Covaris) at 5M cells in 130 µL per tube.

Cells were sonicated on a Covaris LE220 at PIP 300/DF 15/CBP 200 for 20min total at 2min intervals. After sonication, cells were pooled by condition/line from individual microtubes, and pelleted at 21130g/10min. Supernatent was pipetted to a new microtube, and Covaris Shearing Buffer was added to comprise 45 0µL per IP. 10 µL of this was saved at -20°C for ChIP-seq input sample per line/condition. To 450µL 35assette35nt, added 300µL of 2.5X ChIP Buffer (High Triton) (110 mM Tris-HCl pH 7.5, 375 mM NaCl, 4.425% Triton-X, 0.33% SDS + 1X Halt Protease/Phosphatease Inhibitor + 1 mM PMSF). To this we added 2-3 µg of ChIP antibody to each sample as indicated, incubated rotating at 4°C overnight.

ChIP-seq antibodies were used as follows: NKX2-1 (Abcam ab137061; Lot# YJ050719CS; 3µg), H3K27ac (Abcam ab4729; Lot# GR3251519-2; 2.5µg), H3K4me1 (Abcam ab8895; Lot# GR3369517-1; 2.5µg), H3K4me3 (Millipore 05-745R; Lot# 3158071; 3µL), RNAPolII (Diagenode C15200004; Lot# 001-12; 2.5µg), EP300 (CST D2X6N; Lot# 1; 10µL), c-Jun (CST 60AB; Lot# 11; 10µL), FOXA1 (CST E7E8W; Lot# 1; 10µL), FOXA2 (Abcam ab108422; Lot# GR3289185-; 2.5µg), SOX2 (CST D9B8N; Lot# 1; 10µL), CTCF (CST D31H2; Lot# 4; 10µL). See **Supplementary Table 1** for antibody information.

We used 30 µL of G dynabead slurry per IP, washed once in 1X ChIP buffer, and resuspended overnight ChIP lysates in dynabeads. Rotated ChIP lysates in dynabeads for 2hr at 4°C. After rotation, immobilized beads on a magnet, and removed supernatant. Proceeded to wash ChIP samples on dynabeads a total of 12 times, 3 times each in the following 4 buffers in sequence: Wash Buffer 1 (/RIPA150) (10 mM Tris-HCl pH 7.5, 150 mM NaCl, 0.1% SDS. 0.1% Sodium Deoxycholate, 1.0% Triton-X, 1 mM EDTA), Wash Buffer 2 (/RIPA500) (10 mM Tris-HCl pH 7.5, 500 mM NaCl, 0.1% SDS, 0.1% Sodium Deoxycholate, 1.0% Triton-X, 1 mM EDTA), Wash Buffer 3 (/LiCl Buffer) (10 mM Tris-HCl pH 7.5, 250 mM LiCl, 0.5% Sodium Deoxycholate, 0.5% Triton-X) and Wash Buffer 4 (10 mM Tris-HCl pH 7.5). After final wash, any buffer was removed by pipette. Bead pellet was resuspended in Elution Buffer (TE Buffer pH 8.0, 150 mM NaCl, 0.1% SDS, 5 mM DTT) and heated at 65°C for 1hr at 1000rpm. Beads were again placed on magnet, and supernatant was removed to a new tube. 10 µL input sample was also resuspended in 100 µL elution buffer and processed in parallel with ChIP samples. To eluted samples, added 1 µL Rnase (Roche) and incubated at 37°C for 30min at 1000rpm. Next, added 3 µL of Proteinase K (Lifetech) and incubated at 65°C for 3hr at 1000rpm. Placed samples at RT to cool overnight.

To each ChIP, added 160µL of PEG/NaCl (20% PEG, 2.5 M NaCl), then added 100 µL of SPRISelect beads, vortexed, incubated at RT. Washed twice with 80% ethanol, then eluted DNA in 20 µL of 10 mM Tris-HCl pH 8. DNA concentration was quantified using Quant-IT dsDNA HS assay, and libraries were prepared with eluted ChIP DNA using the NEBNext Ultra II DNA Library Prep Kit. Final libraries were quantified using Qubit, pooled, and sequenced by Broad Walk-Up Sequencing on a Nextseq 500 using 2×40bp sequencing.

### ChIP-seq data analysis

We trimmed paired end ChIP-seq reads using Trimmomatic (v. 0.36)^79^ for properly paired reads and to remove any TruSeq adapter sequences. We aligned to the hg38 reference genome (GRCh38_noalt_as) using bowtie2 (v. 2.2.9)^80^ with parameters “-k 1” and converted to a final BAM file with samtools (v. 1.9)^81^. To generate genome-wide occupancy tracks, we used macs2 (v. 2.1.1)^82^ with parameters “-g hs -f BAM --nomodel -B --SPMR” to generate a per million normalized genome-wide track, and then converted to a bigwig file using bedGraphToBigWig.

Narrow ChIP-seq peaks were called using macs2 (v. 2.2.7.1) with parameters “g hs -q qe-3 -f BAMPE --nomodel” against an input control generated from sample cells. Peak calls were filtered against ENCODE hg38 blacklist regions using bedtools subtract with parameter “-A” to remove peaks with any overlap. For determination of the top 10,000 binding sites, ChIP-seq peaks were subsetted by narrowPeak signalValue (column 7). Venn diagrams of ChIP-seq peaks were generated using the R package ChIPpeakAnno, number labels were scaled relative to their numeric values. For distance to transcription start sites, we generated a TSS annotation file using the hg38 refFlat annotation, and calculated the distance of each peak to the nearest TSS using bedtools closest with parameters (-t first -d). Motif analyses were performed using the 500bp surrounding the ChIP-seq peak summit, using bedtools getfasta with the hg38 reference genome, and analyzed using MEME-Suite (v. 5.3.3)^83^. Motif discovery occurred using STREME (v. 5.3.3)^84^, and motif enrichment p-values and site recognition were obtained from analyses using Centrimo (v. 5.3.3)^85^.

For super-enhancer analysis, we used the standard analysis pipeline ROSE (v 1.3.1)^46^ with H3K27ac ChIP-seq peaks. Graphs of ranked enhancer/SE peaks were generated using ROSE merged peak regions using the H3K27ac signal – Input signal for ranking regions.

### Analysis of RNA-seq and TSS-seq from RERF-LC-Ad1

RERF-LC-Ad1 RNA-seq (DRR016714) and TSS-seq (DRR095980) were obtained from Suzuki et al., 2014^44^ and Sereewattanawoot et al. (2018)^86^, respectively. RNA-seq reads were mapped to the human reference genome (hg38) using STAR (v. 2.3.1)^87^ version 2.3.1 with default parameters. RNA-seq tracks were generated using bedtools genomecov –split –scale, normalized using a value of 10^6^ divided by the total mapped read count by samtools (samtools view -c -f 67 -F 256) to generate a bedgraph RNA-seq file normalized per million reads, then convereted to a bigwig using bedGraphToBigWig.

TSS-seq reads were processed as above, except for the following modifications. TSS-seq data were single end whereas RNA-seq were paired end, so reads were mapped as single-end using STAR. TSS-seq reads were stranded (RNA-seq was not)—as such, we split TSS-seq reads between forward and reverse strand using samtools view (-b -F 276 for forward strand reads, -b -f 16 -F 260 for reverse strand reads). Forward/reverse strand tracks were converted to normalized bigwig files as above for RNA-seq.

### Luciferase assay cloning

All luciferase assay cloning occurred into the pGL4.23 miniP-luc2 vector (Promega) to minimize background transcription and enable modulation of enhancer sequences. Genomic DNA was purified from NCI-H441 cells using the DNeasy Blood & Tissue Kit (Qiagen). NKX2-1 SE E1-10 and MYC-E3 enhancer sequences were PCR amplified from NCIH441 genomic DNA using Phusion HF Polymerase (NEB). Subsequent PCR added flanking restriction sites, as well as a BsmBI or BbsI cut sequence to enable specific cutting if an enhancer sequence contains an internal cut site. MYC E3 PCR primers were obtained from (Zhang et al., 2016)^6^. Primer sequences are available in **Supplementary Table 1**. hg38 enhancer coordinates are available in **Supplementary Table 2**. Primers were designed using Primer3.

Full length enhancer sequences were incorporated at XhoI/BglII sites in the pGL4.23 MCS. PCR product was purified, digested, and ligated into the pGL4.23 backbone using Quick Ligase (NEB). Downstream cloning occurred as upstreaming, but cloning into the BamHI/SalI downstream sites using the same XhoI/BglII overhangs, as BamHI/SalI contain matching overhangs. For E7 amplification assays, a second E7 sequence was cloned into KpnI/XhoI sites as above. Primer sequences are available in **Supplementary Table 2.** hg38 coordinates are available in **Supplementary Table 3**.

For 100bp walking fragments, individual E7 sequences were generated by Phusion PCR amplification of E7, and cloned into XhoI/HindIII sites as above. For region deletions and motif deletions, we used an E7(–) inverted sequence to avoid any proximity effects as seen in the walking fragments. Motifs for known transcription factors were identified using FIMO (v. 5.3.3)^88^ on the 692bp E7 sequence. Region and motif deletion constructs were designed using FIMO motif locations, deleting clusters or individual motif binding sites from this sequence. These sequences were synthesized and inserted into the pGL4.23 XhoI/BglII sites by Twist Bioscience. For the minimal E7(301-450) fragment, we synthesized this minimal fragment as a gBlocks Gene Fragment at IDT, digested this fragment using BsmBI, heat inactivated, and ligated into pGL4.23. Synthesized E7 sequences are available in **Supplementary Table 4**.

For promoter reporters, promoter sequences were cloned to replace the miniP sequence using the HindIII/NcoI sites, and include a Kozak sequence (GCCACC) directly upstream of the *luc2* transcription start site. For miniTK, oligos were synthesized, annealed, and ligated into pGL4.23. For the *NKX2-1* promoter, the sequence was PCR amplified from NCI-H441 genomic DNA using Q5 or Phusion polymerases, added Kozak and restriction sites on, digested and ligated into pGL4.23. For *NKX2-1*, a two-step PCR was undertaken so the final cloned promoter contains the same translation start site as the short *NKX2-1* isoform, so as to best model the endogenous locus. *SFTPC* primer sequences Primer sequences are available in **Supplementary Table 2**. hg38 promoter coordinates are available in **Supplementary Table 3**.

### Luciferase assay

Final luciferase assay plasmids were prepared via Plasmid Plus Midiprep (Qiagen), diluted to approximately 100 ng/µL in diH2O, and quantified by Qubit dsDNA BR Assay (Thermo Fisher) on a Perkin Elmer Envision Plate Reader. pGL4.23 is a low copy plasmid so nanodrop quantification was inaccurate in many cases. Plasmid dilutions were also confirmed by restriction enzyme digestion on a DNA gel to confirm similar plasmid presence.

For transfection, a ratio of 15 0ng of pGL4.23-Empty to 50 ng of pRL-CMV was standard, transfected using Lipofectamine 3000 (Thermo Fisher) at 3 µL/µg DNA. For plasmids containing reporter inserts, construct transfections were adjusted so that the same total number of molecules of each plasmid were transfected into cells as the empty vector control. To do this, the plasmid length in bp of each plasmid was divided by the empty vector size, and this scaler was multiplied to the plasmid for assembly of transfection mix.. This was done to avoid post hoc processing to adjust for plasmid size. For standardization, transfection reagent volumes were constant across conditions. We also used a pLX302-GFP transfection control to confirm cell transfection after 24h.

For luciferase assay transfection, cell lines were plated on day -1 at 100 µL/well in 96w assay plates (Corning) in PenStrep-free media at 10-20K cells/well. On day 0, transfection mixes were prepared as follows, and added to cells in biological triplicate. On day 1, cells were imaged for GFP fluorescence, and then luciferase activity was determined using the DualGlo Luciferase Assay System (Promega). Specifically, 40 µL/well of Dual-Glo Luciferase Buffer were added per well, plates were incubated at RT shaking for 15min, spun at 1000g/5min to remove any bubbles, and luminescence was quantified using a Perkin Elmer Envision Plate Reader using a 96w ultrasensitive luminescence aperture with a 2s exposure to enable full dynamic range of quantification. After quantification, we proceeded similarly with Dual-Glo Stop & Glo Buffer—40 µL per well, 15min shaking, 1000g/5min to collect, ultrasensitive luminescence quantification.

### Luciferase assay analysis

For data analysis, firefly reporter luminescence for each well was normalized to the mean *Renilla* luminescence for each condition to account for relative transfection efficiency. All paired conditions were then normalized to the mean levels of the empty vector control to generate plotted arbitrary units (AU) of luciferase activity. Due to the inherent variability of cell plating and transfection efficiency, all experimental comparisons plotted were from the same day on the same cells. However, results were very consistent across distinct experiments. Statistics are two-tailed t-tests to the indicated condition.

### Lentiviral production

Lentivirus was produced by transfection of HEK293T cells using Transporter 5 transfection reagent (Polysciences). For each µg of lentiviral plasmid, 1 µg of psPAX2 and 100 ng of pCMV-VSV-G were combined in Opti-MEM media with 5 µL of Transporter 5 transfection reagent per µg of plasmid transfection. After 15min of incubation, transfection mix was added dropwise to 15cm plate of HEK293T cells in DMEM+10% FBS (without Pen-Strep). The next day, media was changed, and virus production continued for 3 days. Supernatant virus media was collected, spun at 4000g/10min to remove any cells, filtered using a Steriflip 0.45 µm filter (Fisher Scientific), then concentrated using a Amicon 100kDa Centrifugal Filter (Sigma). Concentrate was resuspended in RPMI + 30% FBS + 8 µg/mL Polybrene (SCBT), aliquoted, and stored at -80°C for future use.

### Lentiviral infection

Cells were plated on day -1 in a 6 well plate, on day 0 media is changed to culture media + 8 µg/mL Polybrene. To this, prepared virus is added to well(s), cells were spun at 1178g for 30min-1hr. On day 1 post-infection, cell media is changed to hard selection (1 µg/mL puromycin or 10 µg/mL blasticidin). On day 4, cells were changed to low selection (0.5 µg/mL puromycin or 2.5 µg/mL blasticidin) contingent on cell death in a ‘kill’ (no virus and antibiotic selection) well. Virus was titred to ensure 30-50% cell infection rate (MO1 of 1) as compared to an unselected well of cells. Cells were replated/cultured in low selection media until assay is performed as indicated.

### RIPA lysis

RIPA buffer (50 mM Tris-HCl pH 7.5, 150 mM NaCl, 1% NP-40, 0.5% sodium deoxycholate, 0.1% SDS with 1X Halt Protease/Phosphatase Inhibitor and 1 mM PMSF) was added to cells on ice, vortexed/pipetted to disrupt, incubated on ice for 10min, then spun at 21130g for 10min. Supernatant was pipetted to a new tube, protein extraction was quantified by BCA assay using a SpectraMax M5 at 562 nm absorption. After BCA assay, DTT was added a final concentration of 1 mM, and protein is stored at -20°C for further use.

### Whole cell extraction

Whole cell extract buffer (10 mM Tris-HCl pH 7.5, 1 mM EDTA, 1% SDS with 1X Halt Protease/Phosphatase Inhibitor) were prepared and heated at 98°C. Cell pellets stored at -80°C were kept on dry ice, to which heated 98°C WCE buffer is added, and incubated at 98°C for 10min. Cells were spun to collect, then sonicated using a probe sonicator at 10-20% amplitude for 30s-1min to disrupt the pellet, spun again to collect, and placed back at 98°C for 10min. Sample is spun at 21330g/5min to collect, quantified by BCA assay using a SpectraMax M5 at 562 nm absorption, then added 10X 1 M DTT (final concentration 100 mM) and 10X bromophenol blue to assemble sample buffer for western blot analysis.

### Western blot

For RIPA lysates, samples were assembled using NuPAGE LDS Sample Buffer (4X) and NuPAGE Sample Reducing Agent (10X). For WCE lysates, samples were assembled complete in sample buffer. Samples were heated at 95°C for 10-15min to fully denature protein, then cooled and mixed. 10-20 µg of protein lysates (RIPA or WCE) were standard loading for western blot analysis—PC9 CRISPRa was loaded at 30 µg. Dual Precision Plus (Bio-Rad) protein ladder was loaded at 0.5 µL/well in sample buffer. Samples were loaded onto a Novex WedgeWell Tris-Glycine Mini Protein Gels (Thermo Fisher), 14% gels were generally used with 8-16% and 10% occasionally substituted.

We transferred our western blots to PVDF membrane (Millipore) using a wet transfer approach, transfer occurred for 2hr at 45V in transfer buffer (12.5% Ethanol + 10 mM CAPS + NaOH to adjust pH) at RT. Membranes were incubated for 1hr in TBS-T + 5% w/v milk at RT. Primary antibody incubation occurred in TBS-T + 5% w/v milk, secondary antibody incubation occurred in TBS-T + 5% w/v milk + 0.0025% SDS), either for 1hr at RT or overnight at 4°C rocking.

Membranes were washed with TBS-T between incubations, and were rinsed with diH2O and stored in PBS for imaging. Western blots were imaged on a LICOR Odyssey CLx imaging system, and analyzed/processed in ImageStudio.

Primary antibodies were used as follows: NKX2-1 (Primary/C-Term) (Abcam ab133737 rabbit mAb; Lot# GR154535-2; 1:1000), Beta Actin (CST 8H10D10; Lot# 18; 1:1000), V5 (Invitrogen 46-0705; Lot# 2190378; 1:1000), EIF4A3 (Proteintech 17504-1-AP; Lot# NA; 1:1000), Alpha Tubulin (CST DM1A; Lot# 15; 1:1000), Cyclophilin B (PPIB) (Abcam ab16045; Lot# GR3263788-2; 0.5µg/mL), H3K27ac (CST D5E4; Lot# 8; 1:1000), Histone H3 (CST 96C10; Lot# 10; 1:1000), NKX2-1 (N-Term) (Abcam ab137061; Lot# YJ050719CS; 1:2000). Secondary antibodies were used as follows: Anti-rabbit IgG AF800 (green) (CST 5151S; Lot# 15; 1:30000), Anti-mouse IgG AF647 (red) (CST 5470S; Lot# 14; 1:15000). NKX2-1 (ab137061) is designed to a region within 1- 150AA (N-term) of NKX2-1, NKX2-1 (ab133737) is designed to a region within 300-371AA (C-term) of NKX2-1. See **Supplementary Table 1** for antibody information.

### Cas9 stable line generation

For all stable Cas9 line generation, cells were infected with lentivirus, selected with 10 µg/mL blasticidin for long enough to fully kill an uninfected well, as blasticidin selection can require longer than puromycin. For CRISPR- Cas9, we utilized pLX311-Cas9, for CRISPRi we utilized a pLenti-dCas9-KRAB-MeCP2-Blast as previously designed^89^, for CRISPRa we utilized either pXPR109 (dCas9-VP64-Blast) or pXPR123 (dCas9-p300(core)-Blast). For all but pXPR123 we generated lentivirus according to methods listed above, for pXPR123 we purchased lentivirus from Broad GPP.

### CRISPRi guide cloning

Individual guides targeting E4, E6 and E7 were designed using the Broad GPP guide design tool, screened by NCBI Blast for off-target sites, and selected based on location within the enhancer sequence. A positive control guide targeting *CD81* was also cloned for activity assays. Guides were cloned into lentiGuide-Puro by according to Zhang Lab protocols (https://media.addgene.org/cms/files/Zhang_lab_LentiCRISPR_library_protocol.pdf). Guide oligos were ordered with overhangs and 5’ G (if absent) for transcriptional efficiency from IDT, annealed with T4 PNK (NEB) in T4 Ligation Buffer (NEB), ligated into BsmBI-linearized and gel purified lentiGuide-Puro. Guide sequences are available in **Supplementary Table 5**.

For a triple guide expression cassette, we designed a cassette as follows:

hU6_**E7-sg2_tracrRNA(VCR1)_H1_E6-sg3_tracrRNA(CR3)_7SK_E4-sg2**_tracrRNA(spCas9) [bold sequences were cloned into the lentiGuide-Puro cassette] Full cassette sequence is available in **Supplementary Table 5**.

We designed this using alternative tracrRNA sequences as previously characterized^90^, as well as distinct promoters to minimize recombination post-lentiviral infection, as the guides would need to be expressed continually for CRISPRi. The above cassette was designed into two fragments with BsmBI sites to enable scarless cloning into lentiGuide-Puro, and synthesized by Twist Bioscience. We assembled this using Golden Gate cloning, with Twist fragments, BsmBI-linearized lentiGuide-Puro, 1X Tango Buffer, 1 mM DTT, 1 mM ATP, 1 µL Esp3I/BsmBI, and 1 µL T7 DNA ligase. This was assembled as follows: 99 cycles of 37°C 5min and 20°C 5min, with 37°C 30min, and 65°C 10min to complete.

### CRISPRa guide cloning

Guides were cloned as above—we used the same enhancer guides used for CRISPRa and CRISPRi. Guides were cloned into the pXPR502 (PP7-NLS-HSF1-p65) SAM activation system as for previous guide cloning. A positive control guide targeting *CD4* was also cloned for activity assays. Guide sequences are available in **Supplementary Table 5**.

### CRISPR activity assays

We assayed the activity of each of our stable CRISPR lines after selection as follows.

For CRISPR-Cas9 lines using pLX311-Cas9, we used the PX458 (*<u>GFP</u>*/sg*GFP*) GFP cleavage vector to assay Cas9 cutting activity. We cloned a PX458-sgNT vector to use alongside PX458 and lentiGuide-sgNT in Cas9 expressing lines. We infected cells with lentiGuide-sgNT, PX458, or PX458-sgNT, selected for 10 days, and assayed the percent GFP positive cells by flow cytometry analysis on FITC channels.

For CRISPRi activity assays, we performed activity assays according to Broad GPP standard protocols. Specifically, dCas9-KRAB-MeCP2 lines were infected with a control guide (eg. sgNT) as well a guide targeting the sg*CD81* cell surface marker. Cells were infected, selected, and propagated. On day 7-10, cells were trypsinized, pelleted, resuspended in staining buffer (PBS+2% FBS+5 µM EDTA), pelleted, then resuspended in 99 µL staining buffer + 1 µL of APC-CD81 antibody (Biolegend) alongside unstained controls, incubated on ice for 30min, pelleted and washed twice, then analyzed by flow cytometry.

For CRISPRa activity assays, cells were prepared similarly to CRISPRi via Broad GPP standard protocols. Specifically, cells were infected with pXPR502 with a control guide (eg. sgNT) or a guide targeting *CD4*, and incubated with an APC-CD4 antibody (Biolegend) after selection, and analyzed by flow cytometry.

### Flow cytometry analysis

Cells from experiments as indicated were suspended in staining buffer (PBS+2% FBS+5 µM EDTA), and analyzed by flow cytometry on a CytoFlex S or LX machine. Cells were gated by FSC-A/SSC-A for total cell population and by SSC-A/SSC-H for single cells. Flow analysis and plotting occurred using FlowJo (v. 10.8.2).

### CRISPRa/I characterization of enhancer guide activity

For assaying the ability of designed CRISPRi/a guides to find and target their intended locus, we used the ability of a catalytically-active Cas9 to cut the target locus as a metric of guide activity.

For CRISPRi, we generated a stable Cas9-expressing NCI-H2087 line using pLX311-Cas9 as above. We similarly made a stable Cas9-expressing NCI-H1975 alongside CRISPRa lines. We infected Cas9 lines with lentiGuide-Puro (CRISPRi) or pXPR502 (CRISPRa) guide expression vectors, infected, and selected cells as done with CRISPRa/I experiments. We purified genomic DNA from these cells using the Dneasy Blood & Tissue Kit (Qiagen). We then performed PCR from genomic DNA for E4, E6 and E7 in enhancer and control guide conditions, and performed Sanger sequencing on the enhancer PCR products. We determined the cut percentage for each guide using TIDE (v. 3.3.0)^91^, using the sgNT as the control. If possible, we used both primers for the enhancer PCR product to determine the cutting percentage, however some enhancer guides (ie. E6 sg1) were too close to one end to determine cut percentage from that side. Primer sequences are available in **Supplementary Table 2.**

### shRNA cloning or plasmid acquisition

shRNAs were obtained from the Broad TRC, selecting the 8 best shRNAs targeting *NKX2-1* with the highest intrinsic score, in either pLKO.1 or pLKO_TRC005 as available. shRNA targeting *GFP* was likewise acquired from Broad TRC. shRNA targeting *Luciferase* was selected as a previously used shRNA with low toxicity^78^, cloned into the pLKO_TRC005 backbone using AgeI/EcoRI sites as standard. shRNA sequences are available in **Supplementary Table 6**.

### shRNA infection for RNA-seq and ATAC-seq

H2087 and H441 cells were spinfected with shLuc, sh*NKX2-1*#1 or sh*NKX2-1*#8 in biological duplicate on day 0, changed over to culture media containing hard antibiotic selection (1 µg/mL puromycin) on day 1, then reduced to culture media containing low selection (0.5 µg/mL puromycin) on day 4. On day 7 post-infection, cells were harvested for cell pellets and ATAC-seq. For pellets, cells were pelleted from each replicate, or as a pool between replicates for western blot analysis, at 500g/5min, PBS washed, pelleted again 1000g/5min, aspirated PBS, stored at -80°C for protein and RNA extraction. For ATAC-seq, cells were counted using a ViCell XR, diluted to 100K/mL, counted again, and then 50K cells per biological replicate were used for ATAC-seq (below). Remaining cells from this experiment were propagated, and used for cell cycle analysis (below).

### ATAC-seq sample preparation

ATAC-seq samples were prepared according to the Omni-ATAC protocol^92^. Cells were tagmented using Illumina Tn5 enzyme, purified using a Zymo DNA Clean and Concentrator-5 Kit, PCR amplified for 5 cycles using NEBNext Ultra II Master Mix. Additional amplification was determined using qPCR on a Qstudio 6 FLX real-Time PCR System, with the only modification being inclusion of ROX passive dye for qPCR. Samples were amplified for 2-3 additional cycles, column purified again using a post-ATAC-seq Zymo DNA Clean and Concentrator-5 Kit, quantified by qPCR or Qubit and pooled. ATAC-seq libraries were sequenced by Broad Walk-Up Sequencing on a Nextseq 500 using 2×40bp sequencing.

### ATAC-seq data analysis

We analyzed our ATAC-seq data using a pipeline modeled off PEPATAC^93^. We used skewer (v. 0.2.2) to pair reads and remove any Nextera adapters from fastq files. We performed two rounds of read removal, mapping paired-end reads to the rCRSd (mtDNA) and human_rDNA reference genomes from refgenie, using bowtie2 (v. 2.2.9) and mapping parameters (-k 1 -D 20 -R 3 -N 1 -L 20 -I S,1,0.50 --un-conc $dir), and retaining unmapped reads to either genome. We then mapped reads to the hg38 (GRCh38_noalt_as) bowtie2 reference genome using parameters (--very-sensitive -X 2000). We filtered reads using samtools view (-f 2 -q 10 -@ 8 -b) for quality and then using picard (v. 2.27.4) (MarkDuplicates VALIDATION_STRINGENCY=LENIENT REMOVE_DUPLICATES=true). We also quantified insert metrics using picard (CollectInsertSizeMetrics, M=0.5).

We converted each ATAC-seq BAM file to a bed file using bedtools bamtobed. For each read, we corrected the Tn5 insertion bias for (-4/+5 for –/+ strand) in the bed tile using awk (awk ‘BEGIN {OFS=”\t”} {if ($6 ==”+”) print $1,$2-4,$2-4; else print $1,$3+5,$3+5 }’). We then sorted these Tn5 sites, and removed any sites that corrected to outside the hg38 reference genome. We used bedtools makewindows (-w 100) (v. 2.26.0) to make 100bp windows of the chr1-22,X,Y for hg38. We used bedtools map (-c 2 -o count) to map the individual Tn5 cut sites to the 100bp genome-wide windows. We normalized this bedgraph file such that each 100bp window is the number of Tn5 cut sites per 10M Tn5 cut sites mapped to hg38. We converted this to a bigwig for visualization using bedGraphToBigWig. For tracks representing multiple replicates, we merged the aligned BAM files using samtools merge, and calculated the normalized tracks as above.

For ATAC-seq differential analyses, we merged all ATAC-seq BAM files for a cell line assayed (NCI-H2087, NCI- H441, NCI-H1975, PC9), and called peaks on this merged BAM file using macs2 (v. 2.1.1) with parameters (-q 1e-3 -f BAMPE --nomodel). We used the summits generated by this peak calling, extended by 200bp on each side to generate 400bp windows, and filtered for peaks outside hg38. We mapped Tn5 corrected cut sites to these windows using bedtools map. These counts were used for DESeq2 differential analysis—differential peaks were called based on a Bonferroni-corrected p-value ≤ 1e-3 and a fold-change ≥ 1.25. MA plot of ATAC-seq changes for NCI-H1975 and PC9 occurred using DESeq2 mean counts and fold change values.

Analyses of distance to TSS, venn diagrams, and motif enrichment occurred for these 400bp windows as for ChIP-seq data analyses above. For log2FC values for violin plot analyses, we normalized the ATAC-seq signal in each peak per million cuts in all peak regions, and calculated the FC as above for TPM RNA-seq, with a pseudocount of 1. For principal component analysis of ATAC-seq, we mapped ATAC-seq to a union peak set for cell lines to be analyzed, normalized per million as above. We performed principal component analysis on the top 5% most variable sites, quantile normalized, using the R package pcas.

### RNA-seq sample preparation

RNA was extracted from cell pellets stored at -80°C using an RNeasy Mini Kit (Qiagen) with on-column Dnase digestion using Rnase-Free Dnase Set (Qiagen). RNA was eluted in Rnase-free diH2O, and quantified by nanodrop. RNA integrity was determined using the Agilent RNA 6000 Nano Kit on an Agilent 2100 Electrophoresis Bioanalyzer, all RNA samples used for RNA-seq had a RNA integrity number (RIN) > 9.0. 1000- 2000 µg of RNA was polyA-selected using the NEBNext Poly(A) mRNA Magnetic Isolation Module (NEB) and libraries were prepared using the NEBNext Ultra II Directional RNA Library Prep Kit (NEB). Final libraries were quantified using Qubit, pooled, and sequenced by Broad Walk-Up Sequencing on a Nextseq 500 using 2×40bp sequencing.

### RNA-seq data analysis

RNA-seq reads were mapped to the human reference genome (hg38) using STAR (Dobin et al., 2013) version 2.3.1 with default parameters. Raw read counts were generated with Rsubread featureCounts against the hg38 refFlat annotation from UCSC—reads were allowed to map to multiple annotations (allowMultiOverlap=T) but we only counted the primary alignment of each read (primaryOnly=T). Significance was assessed using the R package DESeq2 (v. 1.38.3)^94^. Significantly changing genes were assessed with a Bonferri-corrected p value≤1e- 3 and a fold change≥1.5 to determine set of significantly changing genes. DESeq2 files were filtered for genes that were expressed (mean count < 1), as well as to remove small RNA genes (MIR and SNO). Transcripts per million (TPM) normalized expression values were calculated from featureCounts raw counts normalized by all mapped reads per million. Principal component analysis of cell line RNA-seq occurred on the top 5% most variable genes by log2(TPM+1) expression, quantile normalized, using the R package pcas.

We calculated the log2(fold change) from TPM normalized RNA-seq values using a pseudocount of 1, i.e., log2((TPMExp. + 1)/(TPMCt + 1)). For ranked analyses, we calculated the average TPM across all replicates in both conditions, and calculated as above between mean values. For heatmap and replicate-specific fold change calculations, we calculated the fold change of each replicate above against the average of all control RNA-seq replicates. We excluded any gene as “not expressed” if no replicate had a TPM expression ≥1. These final pseudo-count normalized fold changes were used for GSEA preranked analysis (v. 4.3.2)^95^.

RNA-seq tracks were generated using bedtools genomecov –split –scale, normalized using a value of 10^6^ divided by the total mapped read count by samtools (samtools view -c -f 67 -F 256) to generate a bedgraph RNA-seq file normalized per million reads, then convereted to a bigwig using bedGraphToBigWig. For tracks representing multiple replicates, we merged the aligned BAM files using samtools merge, and calculated the normalized tracks as above.

### Overexpression cloning

pOTB7-NKX2-1 containing the full *NKX2-1* coding sequence was obtained from the Mammalian Gene Collection (MGC) from Dharmacon. The *NKX2-1* coding sequence for both long and short isoforms, as well as with or without a stop codon (to allow C-terminal gateway tagging) were PCR amplified from pOTB7-*NKX2-1*. Primer sequences are available in **Supplementary Table 2.** PCR overhangs included a GGCACC Kozak sequence as well as ATTB1/2 sites for BP cloning into pDONR223. *NKX2-1* sequences were cloned into pDONR223 using BP clonase, transformed into Stbl3 bacteria, plated, miniprepped, and validated by Sanger Sequencing. *NKX2-1* overexpression constructs were generated by introduction of the short *NKX2-1* isoform with an open C-terminal reading frame into pLX302, pLX306, and pLX307 constitutive expression vectors using LR clonase. A C-terminally open *GFP* ORF in pDONR223 was cloned into pLX307 as an expression control. UBC pLX expression vector was generated by cloning from Luc.Cre vector (Addgene #20905) and SFFV pLX expression vector was generated by cloning from SFFV-DTAG-NTERM-GFP (Addgene #185760). For GFP fusion vectors a V5-linker-GFP fusion was cloned from SFFV-DTAG-NTERM-GFP (Addgene #185760) to replace the C-terminal V5 tag in pLX vectors.

### CRISPR-Cas9 guide cloning

Guides targeting *NKX2-1* were designed using the GPP sgRNA picker (now CRISpick)^96^, and 5 top scoring guides were selected, as well as non-targeting (NT), *GFP* targeting, and intergenic Chr2.2 controls, and cloned into the lentiCRISPRv2 backbone using oligonucleotide annealing/restriction digest/ligation as standard. Guides were also cloned into lentiGuide-Puro for use in cells with constitutive pLX311-Cas9 expression in individual cases.

### CRISPR-Cas9 infection for RNA-seq

NCI-H358, NCI-H2087, and NCI-H441 were infected similarly to above. lentiCRISPRv2 with sgNT or sg*NKX2-1*#2 were infected in biological triplicate on day 0, changed over to hard selection (1 µg/mL puromycin) on day 1, then reduced to low selection (0.5 µg/mL puromycin) on day 4. On day 7 post-infection, cells were harvested for cell pellets from each replicate as well as pooled for each condition. Cells were used for RIPA lysis as well as RNA- seq.

### shRNA infection for clonogenic assay

Cells were spinfected with shRNAs on day 0, changed to media containing hard antibiotic selection (1 µg/mL puromycin) for 3 days, then low selection until day 7. Cells were trypsinized/replated 1-2 times to remove any dead cells. On day 7, cells were trypsinized, counted using a ViCell XR, diluted to 100K/mL, and counted again. With these final counts, cells were diluted to 10K/mL in culture media, and 4 wells per shRNA are plated in technical quadruplicate from the same lentiviral infection. Pellets of cells on day 7 post-transduction were harvested and stored at -20°C/-80°C for western blot analysis. Clonogenic assay cells are cultured until control shLuc condition was near-confluent, then stained by crystal violet as below.

### shRNA infection for proliferation assay

Cells were infected with shRNAs on day 0, spinfected, hard selection (1 µg/mL puromycin) for 3 days, then low selection until day 7 as above. On day 7, cells were counted, and plated at 50K/well in a 6 well plate in selection-free media. On day 11, day 16, and day 20 (post-infection), two wells of each shRNA infection (technical duplicates deriving from the same lentiviral infection and plating) were trypsinized and quenched in a total volume of 1 mL, and counted on a ViCell XR.

### Crystal violet staining and quantification

Crystal violet staining solution (0.5% crystal violet in 25% methanol) was prepared in 1L, and filter sterilized after full dissolution of the crystal violet powder. For crystal violet staining, culture media was aspirated off 12 well plates, and 0.5-1 mL of staining solution was added. Plates were incubated for at least 1h at RT on a rocker to fully stain cells. Plated were washed using a water bath, and incubated shaking with diH2O to fully wash for 1-2 rounds. After washing, plates were incubated at RT inverted to dry. Crystal violet plates were imaged using an Epson Perfection V600 Photo scanner, images were cropped at well boundaries, representative well shown.

For crystal violet quantification, 1 mL of 10% acetic acid was added to each crystal violet stained well, and incubated shaking at RT for at least 30 min to resuspend crystal violet. From this, 100 µL was pipetted into a 96- well plate in technical triplicate for each well by multichannel pipette. Absorption was quantified by Spectramax M5 absorption at 590 nm. For the technical triplicate quantification of each well, we used the median value of each well for further analyses. All conditions for a cell line/experiment were normalized to the mean absorption value of the control condition (siCt, shLuc, sgNT).

### siRNA transfection

Dharmacon SmartPool siRNAs targeting *NKX2-1* as well as a nontargeting control (siCt), a positive gene-targeting control (si*PPIB*), and a positive cell-killing control targeting a pan-essential gene (si*EIF4A3*) were obtained from Horizon Discovery. Acquired siRNAs were resuspended in 1x siRNA buffer 60 mM KCl, 6 mM HEPES-pH 7.5, and 0.2 mM MgCl2, quantified by nanodrop, diluted to approximately 10 µM in 1x siRNA buffer, and quantified again by nanodrop to calculate the final siRNA concentration. siRNA sequences and information are available in **Supplementary Table 7.**

For siRNA transfection, cells were transfected by reverse transfection with media changed after 48h of incubation. siRNA and RNAiMAX transfection reagent (at 0.3 µL RNAiMAX per pmol of siRNA) were combined in Opti-MEM serum free media for a final siRNA concentration of 10 nM upon addition to culture media, in 1/10 the volume of culture media (100 µL transfection mix per 1 mL culture media). Cells were plated in PenStrep-free media to avoid additional toxicity upon transfection.

For siRNA validation by western blot, 500K-1M cells were plated in a 6 well in 2 mL of media with 200 µL of siRNA transfection mix. Cells were harvested after 48h, stored at -80°C, lysed by whole cell extraction, and analyzed by western blot.

For clonogenic assay, 10k cells were plated in a 12-well plate in 1 mL of media with 100 µL of siRNA transfection mix. 3 wells of each siRNA (Ct, PPIB, NKX2-1, EIF4A3) are performed as biological triplicate. Plates were tapped/mixed to ensure even plating, and after 48h media were changed to 2 mL of PenStrep-free media per well. Cells were maintained with media changes as needed until the siCt well approaches confluence, anywhere from 7-18 days, though most cells reached confluency in 10-14 days. Cells were stained by crystal violet as above, washed, dried at RT, then resolubilized with 10% acetic acid and and quantified as outlined above.

### Cell cycle analysis

Cell cycle analysis on NCI-H2087 and NCI-H441 cells infected with shLuc, sh*NKX2-1*#1, and sh*NKX2-1*#8 in biological duplicate (see RNA/ATAC-seq methods). Cells were plated on day -1 at 3M cells in a 10cm plate, treated on day 0 with 10 µM BrdU for 2hr in culture at 37°C. ∼1 million cells of each condition were analyzed using the FITC BrdU Flow Kit (BD Biociences), assayed on a CytoFlex S flow cytometer.

### EGFR inhibitor treatment

Cells were plated as above for clonogenic assays in 12 well plates at 10K cells per well after shRNA or overexpression infection and selection, and then Osimertinib was added 1 day after plating. Drug media was refreshed 1-2 times over 10 days of treatment, then crystal violet stained and quantified as above. Relative viability was normalized to 0nM control for each condition.

## QUANTIFICATION AND STATISTICAL ANALYSIS

All bar graphs and dot plots show mean + SEM as indicated. For box and whisker plots, the whiskers are either mean to max or 1-99 percentile as labeled. A two-tailed t-test was used for calculating significance, p-values and significance levels are as indicated. A fisher exact test was used for analysis of ATAC-seq and ChIP-seq binding skew to sites of decreased accessibility, p-values are as indicated. R (v. 4.2.2) or Prism 9 were used for statistical analysis.

### Data and Code Availability

ChIP-seq, RNA-seq and ATAC-seq data generated in this study have been deposited at GEO under accession number **GSEXXXXXX**. This paper does not report original code.

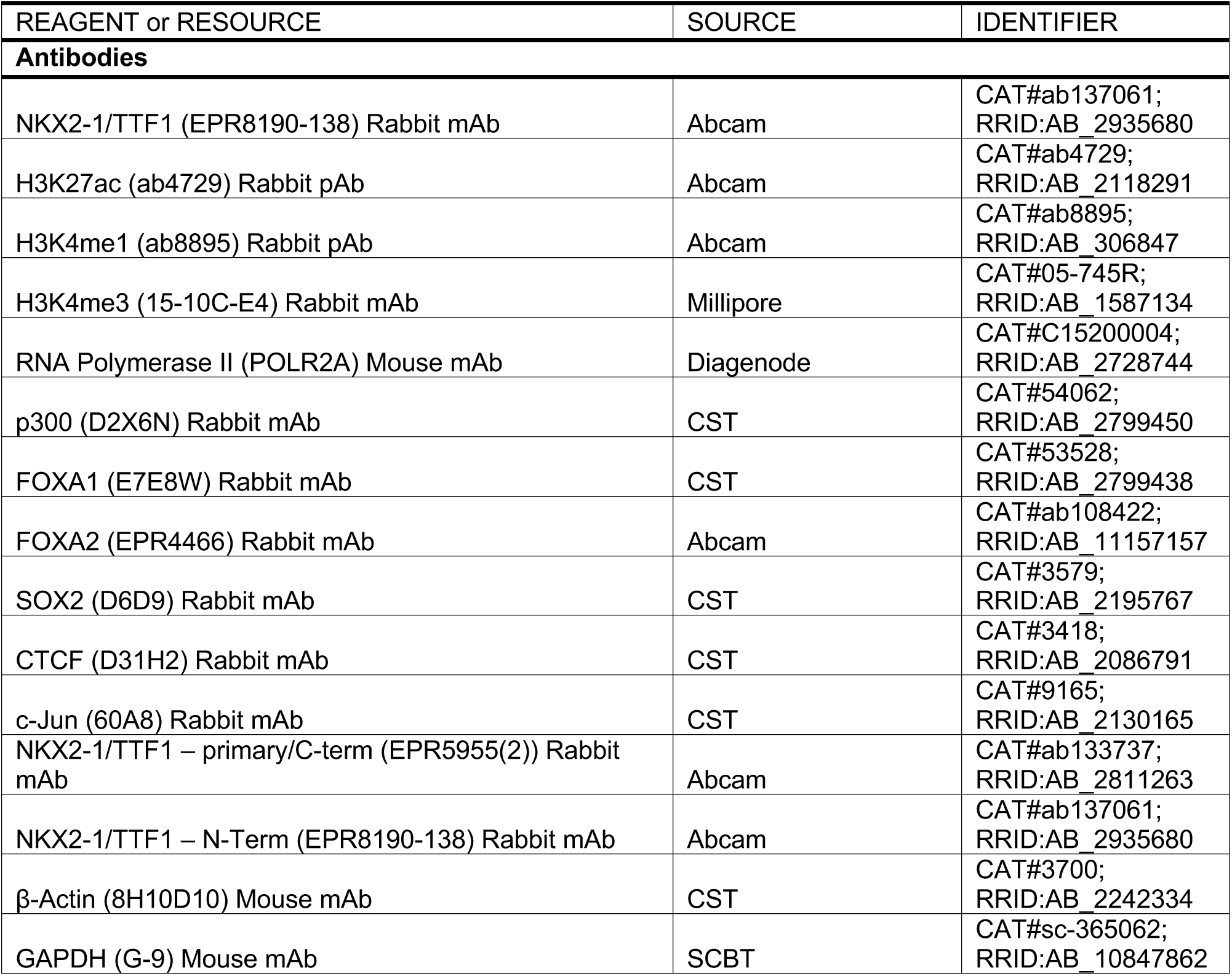
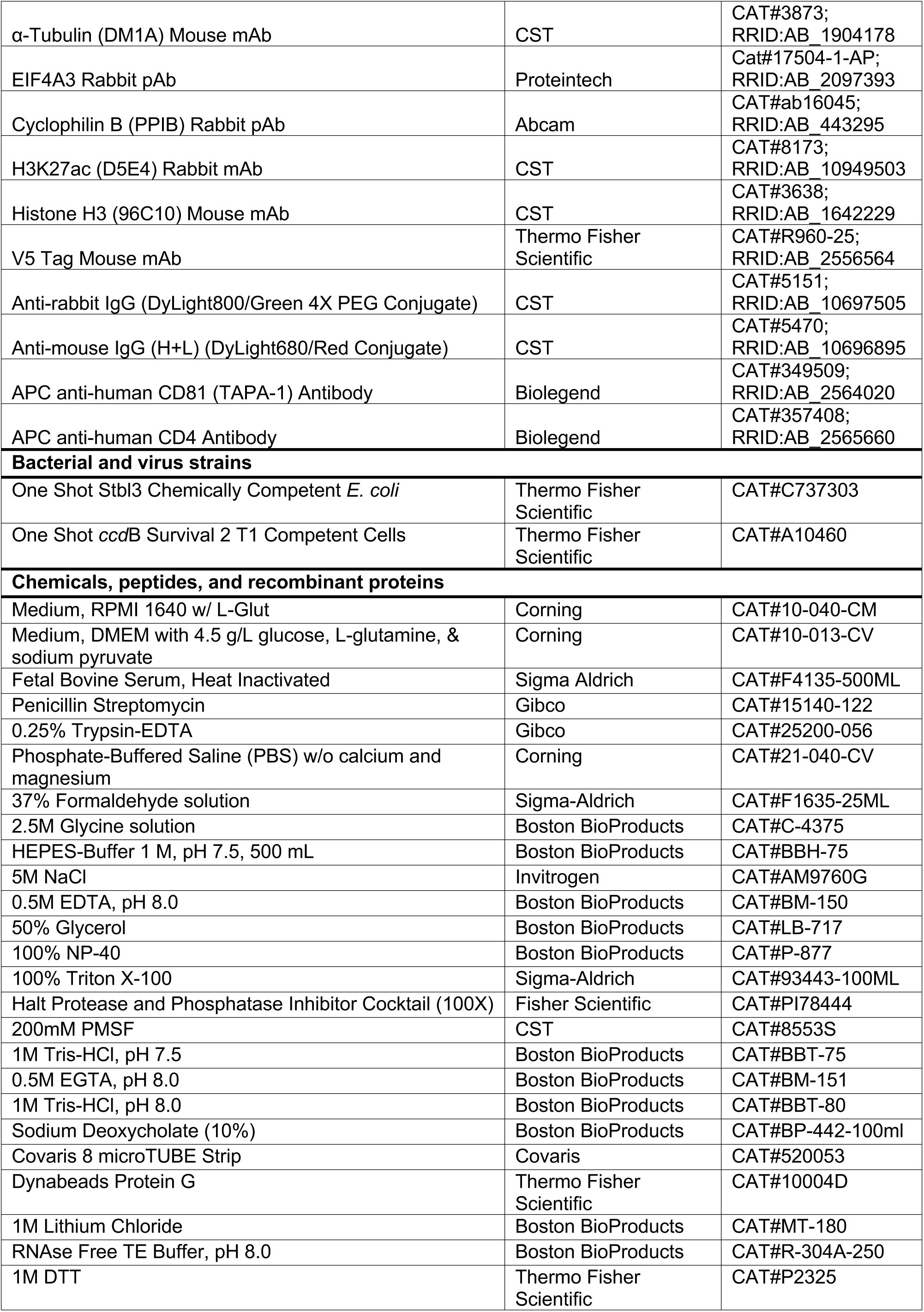
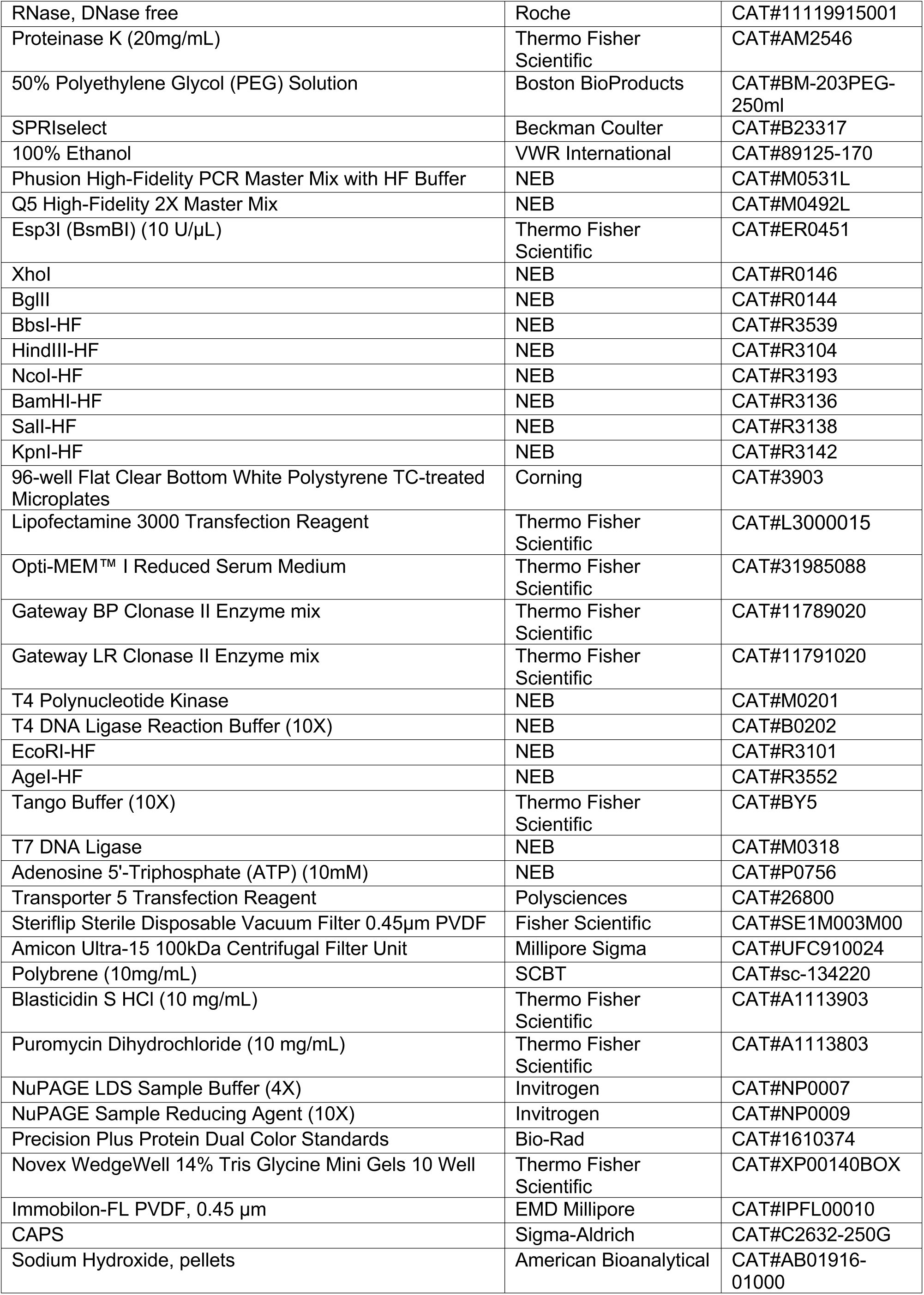
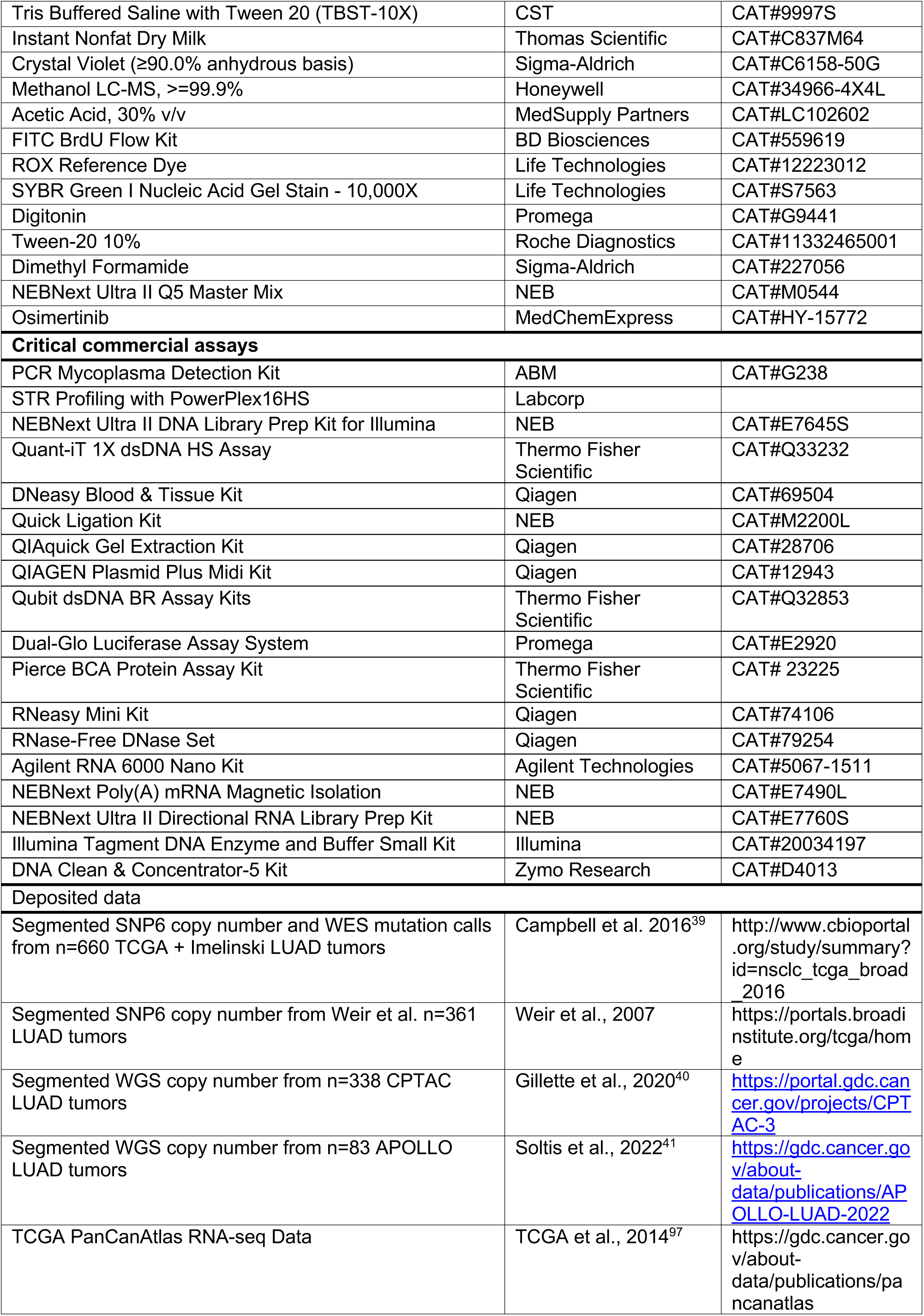
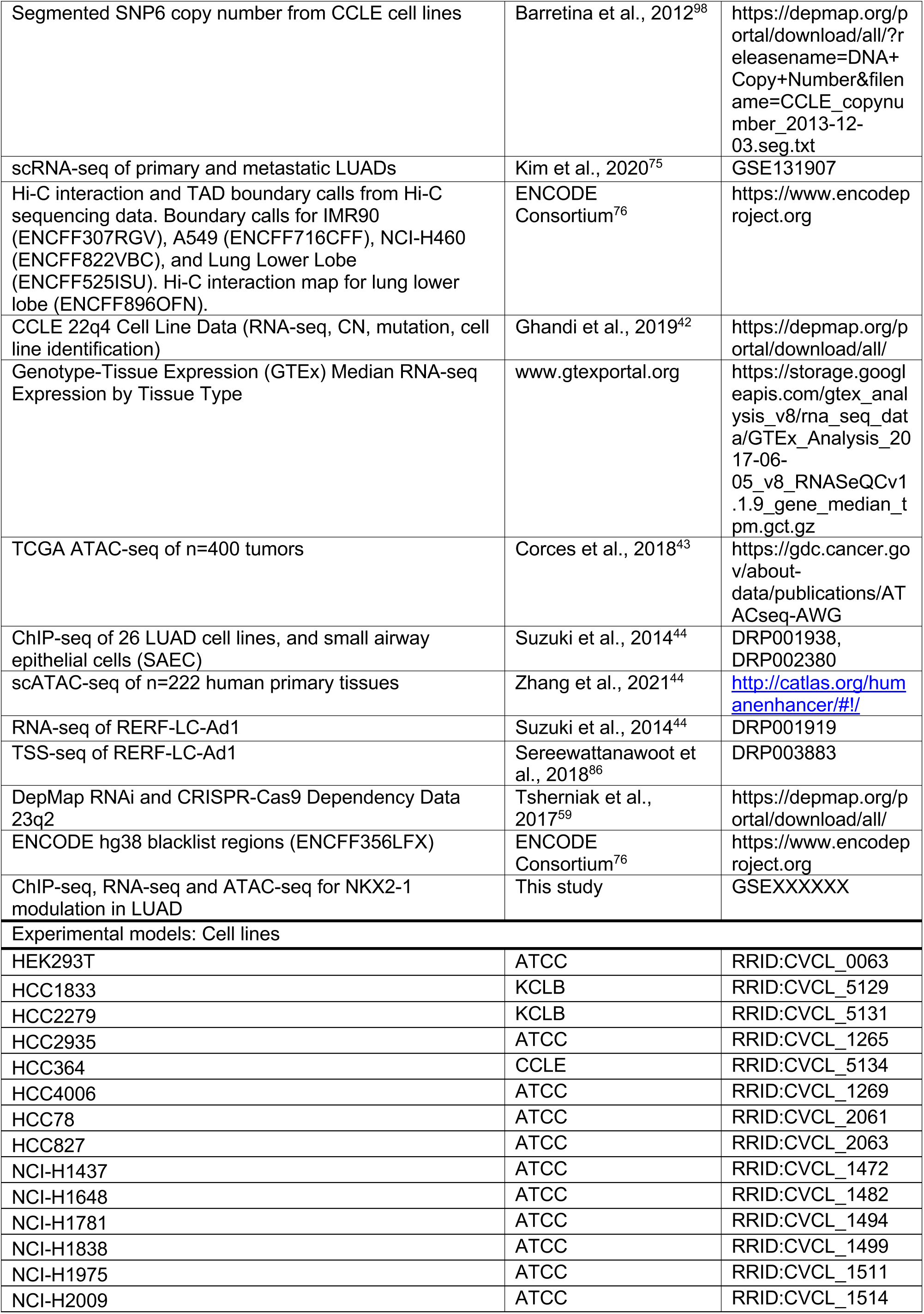
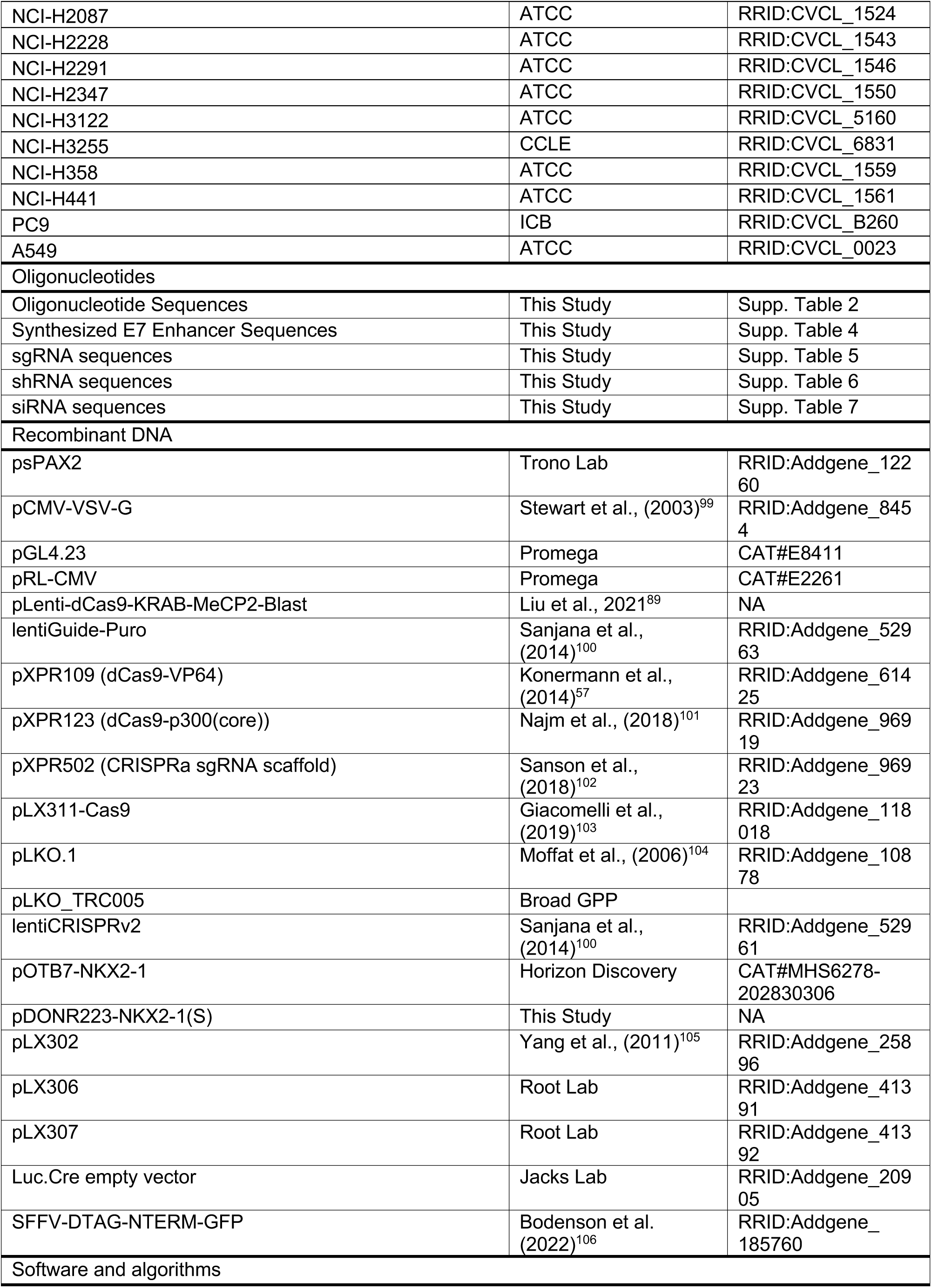
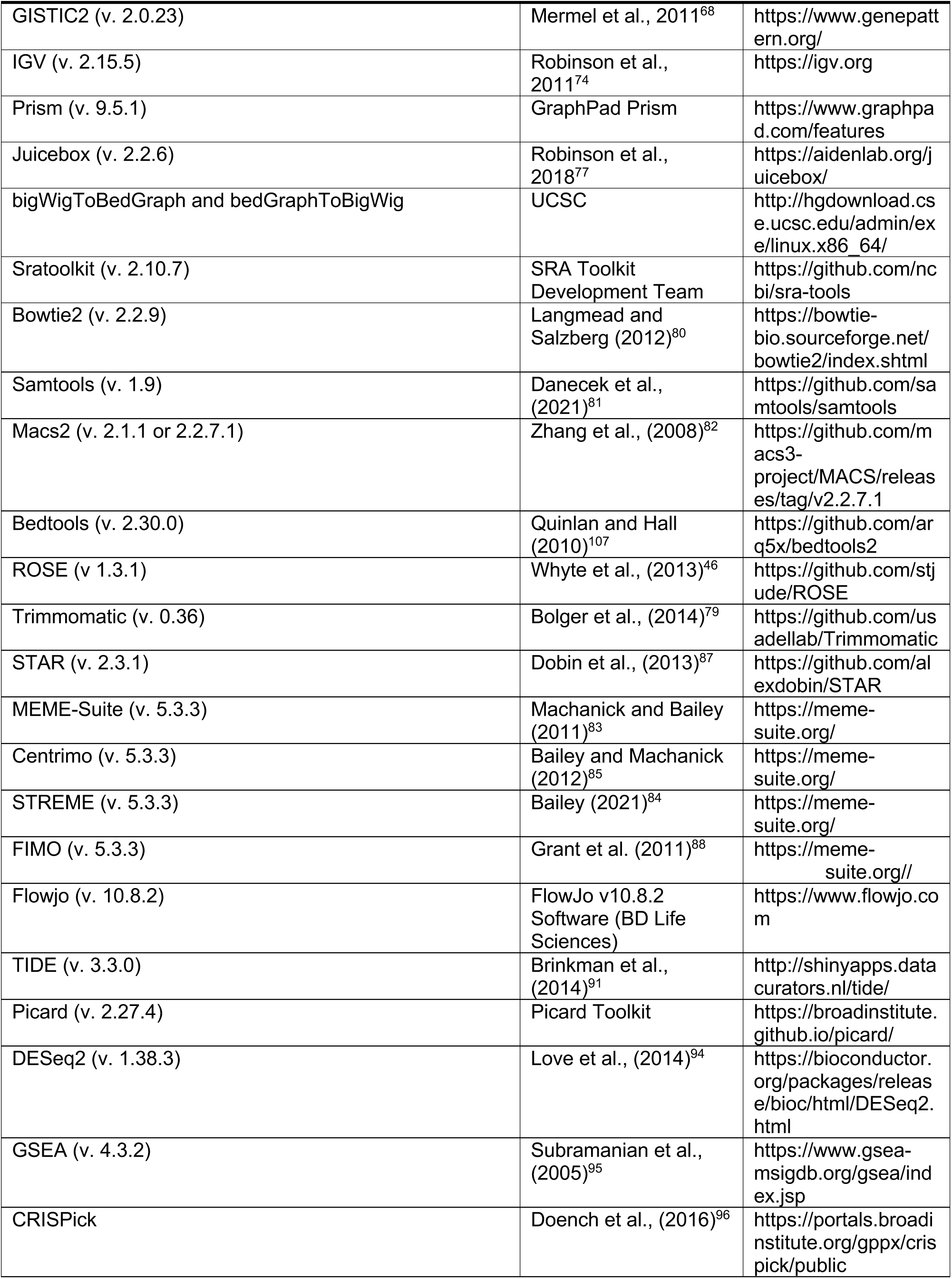
KEY RESOURCE TABLE.

## SUPPLEMENTARY FIGURES AND LEGENDS

**Supplementary Figure 1.**
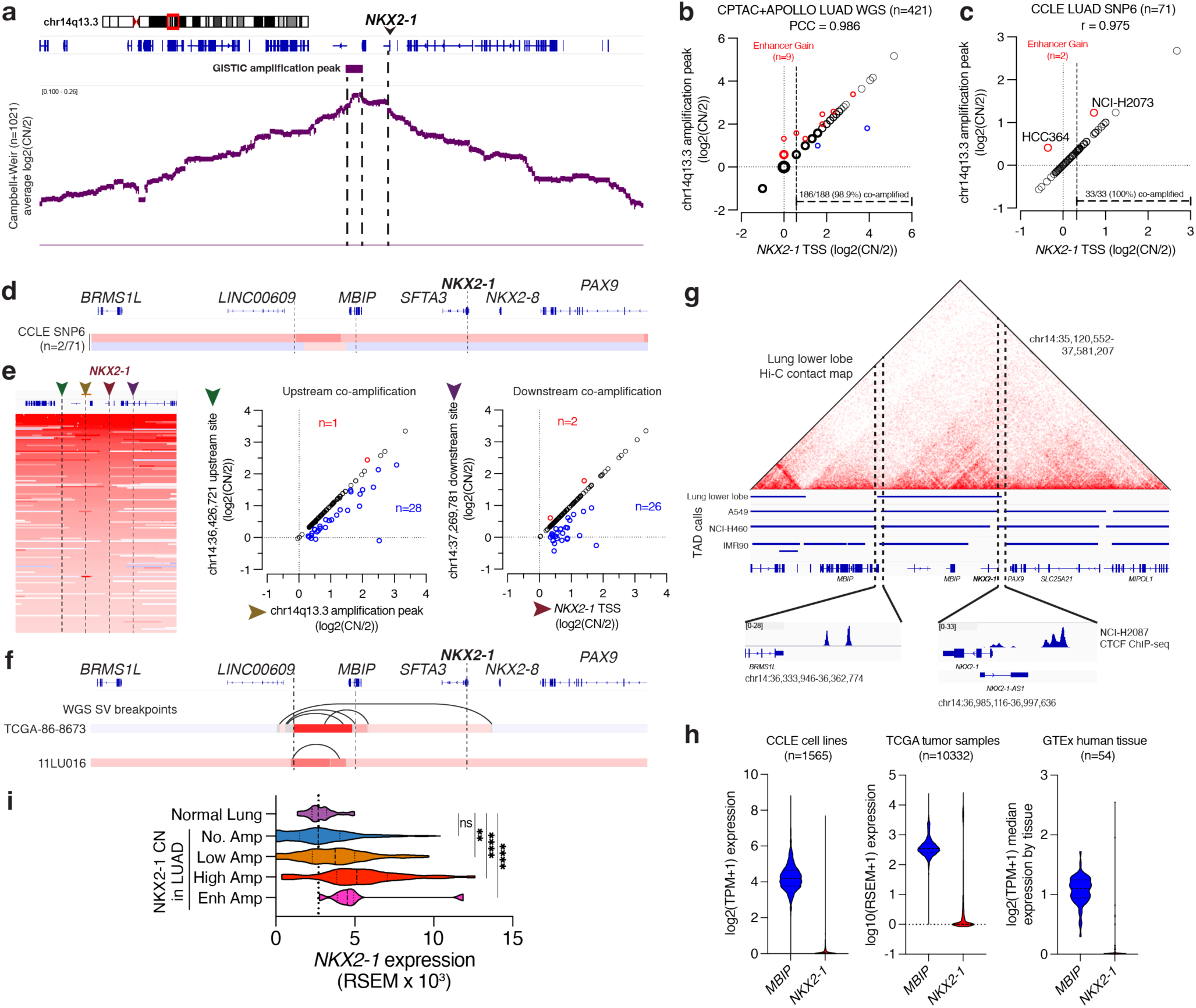
The chr14q13.3 amplification peak targets a non-coding region near *NKX2-1*, related to. **Figure 1** **(A)** (top) The chr14q13.3 focal amplification peak called by GISTIC 2.0 and (bottom) Average log2 copy number profile in SNP6 copy number profiles of Campbell+Weir^20,39^ LUAD tumors (n=660). The chr14q13.3 amplification peak comprises a 116kb non-coding region that is 206 kb centromeric to *NKX2-1*. **(B)** Plot of copy number at the *NKX2-1* transcription start site (TSS) vs. the chr14q13.3 amplification peak in whole genome sequencing (WGS) copy number profiles of CPTAC and APOLLO^39,41^ LUAD tumors (n=421)^40^. Segmented copy number profiles are called at integer values, point densities are indicated by circle size. Focally amplified tumors (n=9) are labeled in red. **(C)** Plot of copy number at the *NKX2-1* TSS vs. the chr14q13.3 amplification peak in SNP6 copy number profiles of CCLE LUAD cell lines^98^. Focally amplified cell lines (n=2) are labeled in red. **(D)** Copy number profiles of samples with focal amplification of the chr14q13.3 amplification peak near *NKX2-1* in CCLE^98^ LUAD cell line SNP6 data (n=71). **(E)** (left) Copy number profiles for *NKX2-1* amplified LUADs from Campbell et al.^39^, with *NKX2-1* transcription start site (TSS) (red) and the chr14q13.3 amplification peak (orange) marked. A region equidistant upstream (green) and downstream (purple) were selected for comparison of amplification rate. (right) Plots of copy number for chr14q13.3 amplification peak vs. upstream site, and for the *NKX2-1* TSS vs. downstream site. Amplification is lost upstream in 28/150 (18.7%) tumors with copy number gain at the chr14q13.3 amplification peak, and is lost downstream in 26/149 (17.4%) with copy number gain of the *NKX2-1* TSS. Co-amplification decays rapidly away from the *NKX2-1* locus. **(F)** Structural breakpoints by WGS (black lines) and copy number profiles by WGS from TCGA and CPTAC for 2 tumors harboring focal amplification of the chr14q13.3 amplification peak. Focal amplification of the chr14q13.3 amplification peak occurs in tandem. **(G)** (top) Hi-C contact map of the *NKX2-1* locus in primary human lower lung lobe. (middle) TAD contact boundary calls in LUAD cell lines and primary lung tissue. (bottom) CTCF ChIP-seq occupancy in NCI-H2087 cells at the putative TAD boundary elements for the *NKX2-1* TAD. **(H)** Truncated viiolin plots of RNA-seq expression of *MBIP* (blue) and *NKX2-1* (red) in (left) all CCLE cell lines, (middle) all TCGA tumor samples, and (right) all GTEx normal human tissues. *MBIP* is ubiquitously expressed whereas *NKX2-1* is tightly lineage restricted. **(I)** *NKX2-1* RNA-seq normalized expression (RSEM) from TCGA normal lung samples (n=59), or TCGA LUAD tumors with low *NKX2-1* amplification (n=87), high *NKX2-1* amplification (n=30), or chr14q13.3 focal amplification (n=7). Two-tailed t-test between normal lung and LUAD samples, ** p<0.01, **** p<0.0001.

**Supplementary Figure 2.**
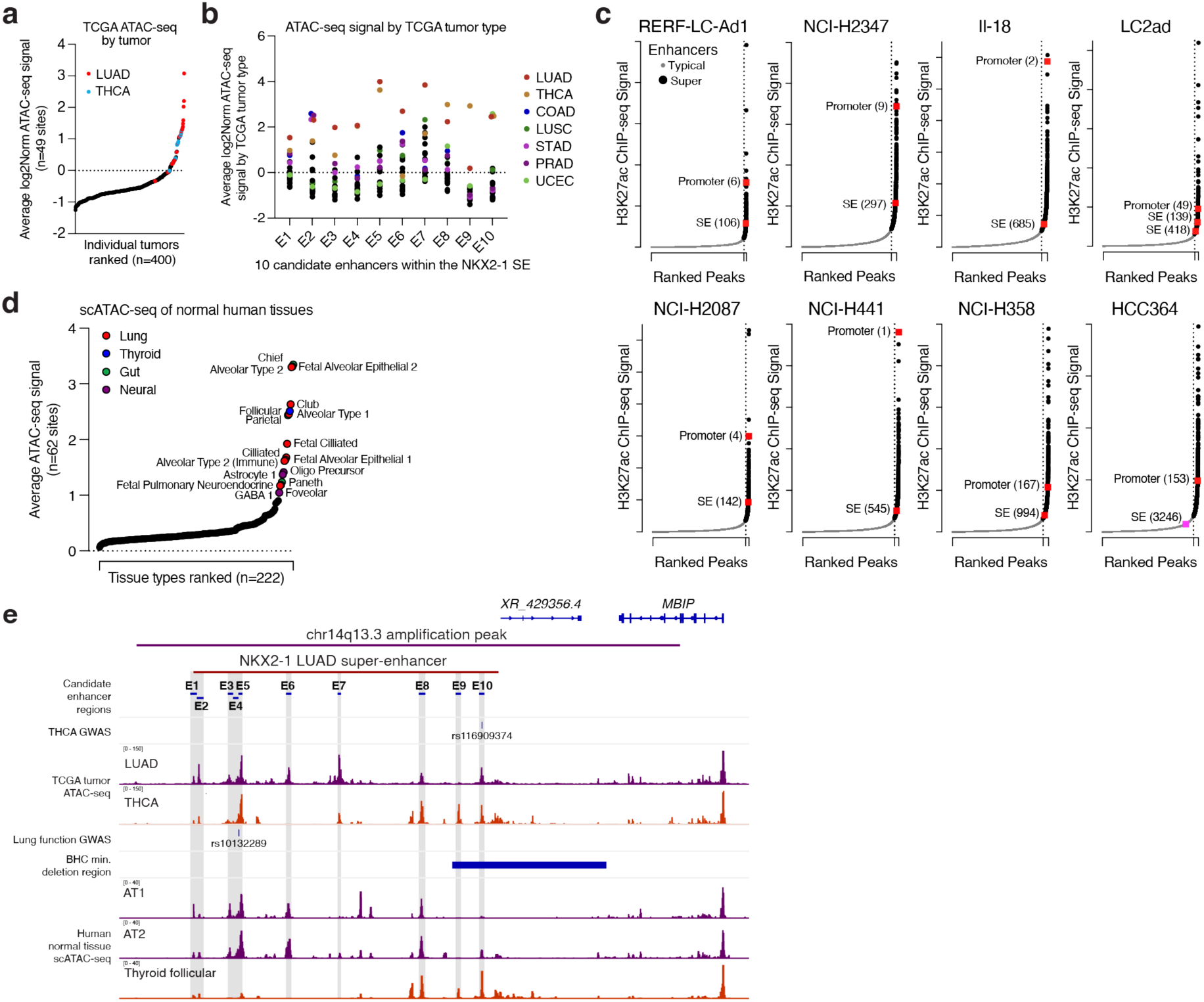
Focal amplification targets a lineage super-enhancer near *NKX2-1*, related to. **Figure 1** **(A)** Average log2 normalized ATAC-seq signal^43^ across called TCGA ATAC-seq peaks (n=49) within the chr14q13.3 amplification peak across n=400 TCGA primary tumors, ranked by signal. LUAD (red) and THCA (orange) tumors labeled. **(B)** Average log2 normalized ATAC-seq signal^43^ at the 10 enhancers of the *NKX2-1 SE* from TCGA ATAC-seq data for n=22 TCGA primary tumor types. **(C)** ROSE SEranking of H3K27ac signal from published (n=4) and newly-generated (n=4) ChIP-seq^44^ in 8 total NKX2-1(+) LUAD cell lines. *NKX2-1* promoter and SEregions are labeled. **(D)** Average normalized scATAC-seq signal^44^ across called ATAC-seq peaks (n=62) within the chr14q13.3 amplification peak across n=222 annotated cell types, ranked by signal. Lung (red), gut (green), thyroid (blue), and neural (purple) cell types with outlier accessibility at the *NKX2-1* SE are labeled. **(E)** The *NKX2-1* SE harbors alleles associated with thyroid cancer and lung developmental disorders (top) The chr14q13.3 amplification peak, LUAD *NKX2-1* SE region, and individual enhancer regions identified herein. Thyroid carcinoma (THCA) risk associated allele rs116909374 identified by GWAS (Gudmundsson et al., 2012)^48^ in the E10 enhancer. Average TCGA ATAC-seq accessibility for LUAD and THCA tumors. (bottom) Minimal deletion region identified in a family for benign hereditary chorea (BHC) in (Invernizzi et al., 2018)^51^—multiple families with *NKX2-1* proximal deletions not targeting the *NKX2-1* gene have been documented^51^. (middle) A lung function (FEV1/FVC) associated allele rs10132289 identified (Shrine et al., 2023)^49^. Merged scATAC-seq accessibility profiles^47^ for alveolar type 1 (AT1) and alveolar type 2 (AT2) distal lung cells as well as for thyroid follicular cells.

**Supplementary Figure 3.**
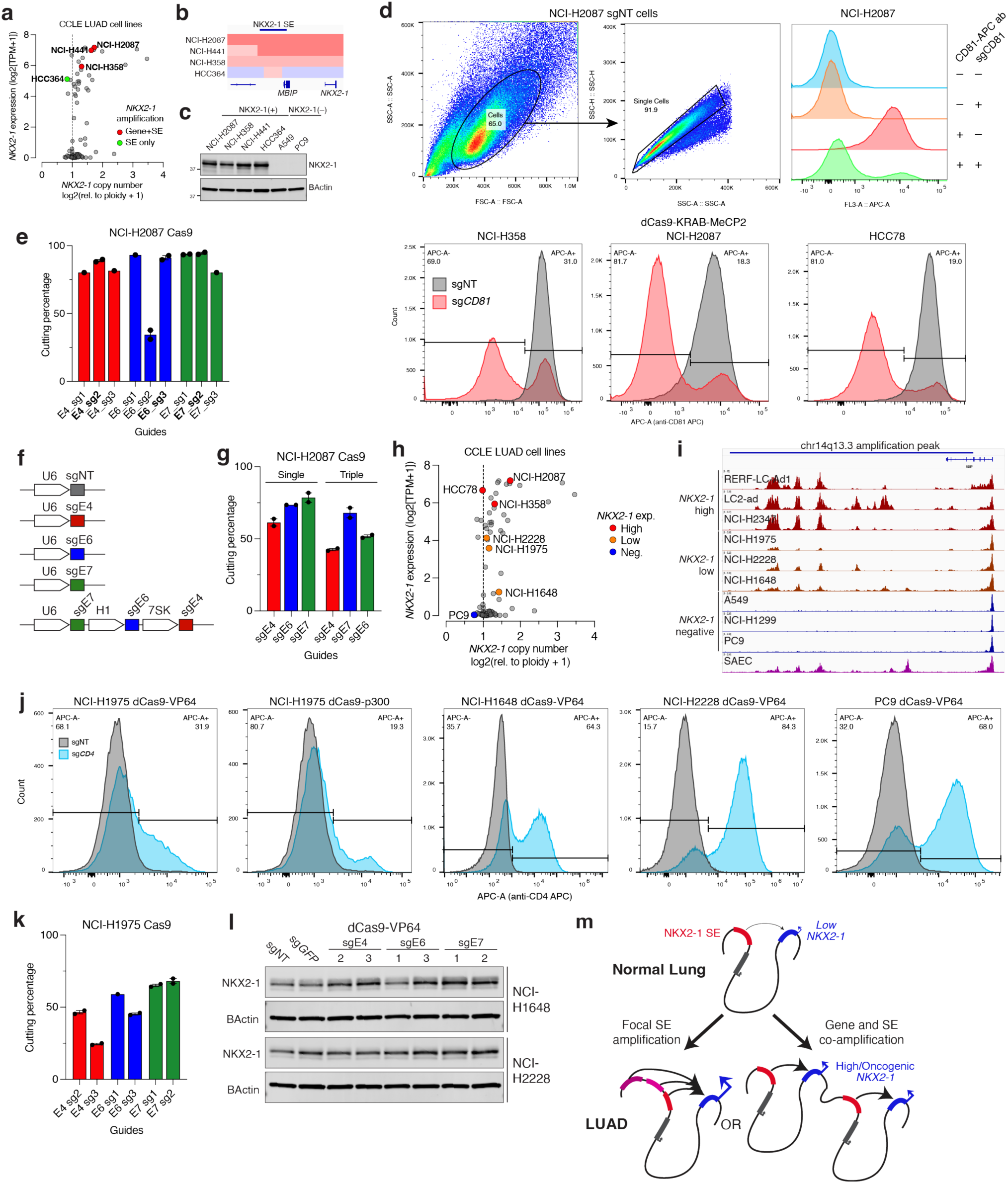
The NKX2-1 SE controls NKX2-1 expression in LUAD, related to. **Figure 2** **(A)** Plot of CCLE LUAD cell lines *NKX2-1* copy number vs. RNA-seq expression. Cells selected for enhancer validation harbor either focal amplification of the *NKX2-1* SE (green) or co-amplification the *NKX2-1* gene and SE. **(B)** Copy number profiles for 4 NKX2-1(+) LUAD cell lines at chr14q13.3, with *NKX2-1* and *MBIP* indicated. **(C)** Immunoblot analysis of NKX2-1 levels in 4 NKX2-1(+) and 2 NKX2-1(–) LUAD cell lines. **(D)** (top) Example gating schematic for NCI-H2087 dCas9-KRAB-MeCP2 cells for activity assay (bottom) Activity assay for CRISPRi Cas9 activity by flow cytometry in 3 NKX2-1(+) LUAD cell lines. Cells expressing dCas9-KRAB-MeCP2 were transduced with either sgNT (grey) or sgCD81 gene-repression guide control (red), and cells were assayed for repression of CD81 expression using an APC-CD81 antibody. **(E)** Cutting percentages for individual sgRNAs targeting E4, E6, or E7 in NCI-H2087 Cas9 cells. Individual guides were stably transduced into NCI-H2087 cells expressing catalytically-active Cas9. Indel percentage by sanger sequencing was used to assay guide activity in cells, and quantified using TIDE^91^. **(F)** Schematic for expression of single guides targeting E4, E6, or E7 within the *NKX2-1* SE, or a three guide expression cassette for coordinate repression of the *NKX2-1* SE by CRISPRi. **(G)** Cutting percentages for individual sgRNAs targeting E4, E6, or E7, or a triple guide expression cassette, in NCI-H2087 Cas9 cells. E4 and E7 cutting %s are decreased in part due to recombination between E4 and E7 resulting in failure to PCR amplify cut alleles. **(H)** Plot of *NKX2-1* copy number vs. RNA-seq expression across CCLE LUAD cell lines. Cells harboring high (red), low (orange) or negative (blue) *NKX2-1* expression for CRISPRi/a modulation of the *NKX2-1* SE are labeled. **(I)** H3K27ac ChIP-seq occupancy tracks for individual LUAD cell lines that are NKX2-1 high (red), low (orange) or negative (blue), as well as for SAEC, at the *NKX2-1* SE. **(J)** Activity assay for CRISPRa Cas9 activity by flow cytometry in 4 NKX2-1 low or negative LUAD cell lines. Cells expressing dCas9-VP64 or dCas9-p300(core) were transduced with either sgNT (grey) or sgCD4 gene-activation guide control (blue), and cells were assayed for activation of CD4 expression using an APC-CD4 antibody. **(K)** Cutting percentages for individual sgRNAs targeting E4, E6, or E7 in NCI-H1975 Cas9 cells. Individual guides were stably transduced into NCI-H1975 cells expressing catalytically-active Cas9. Indel percentage by sanger sequencing was used to assay guide activity in cells. Notably, the E6 sg1 guide fails to activate *NKX2-1*, however the guide is active and targets E6, suggesting a positional specificity for NKX2-1 activation in the *NKX2-1* SE. **(L)** Immunoblot analysis of NKX2-1 levels in NCI-H1648 or NCI-H2228 cells (NKX2-1 low) upon CRISPRa-mediated activation of the E4, E6, and E7 enhancers within the *NKX2-1* SE. Cells expressing dCas9-VP64 for CRISPRa were transduced with guides targeting E4/E6/E7, or control guides. **(M)** Model for *NKX2-1* amplification in LUAD. *NKX2-1* is targeted by either focal amplification of the *NKX2-1* SE, or co-amplification of the *NKX2-1* gene and SE, to activate *NKX2-1* expression in LUAD tumors. **(E,G,K)** Error bars = mean±SEM, 1-2 technical replicates, individual points labeled. If possible, the cutting percentage was assayed using multiple sanger sequencing primers from opposite directions. For n=1 samples, cutting percentage could only be reliably assayed using one primer.

**Supplementary Figure 4.**
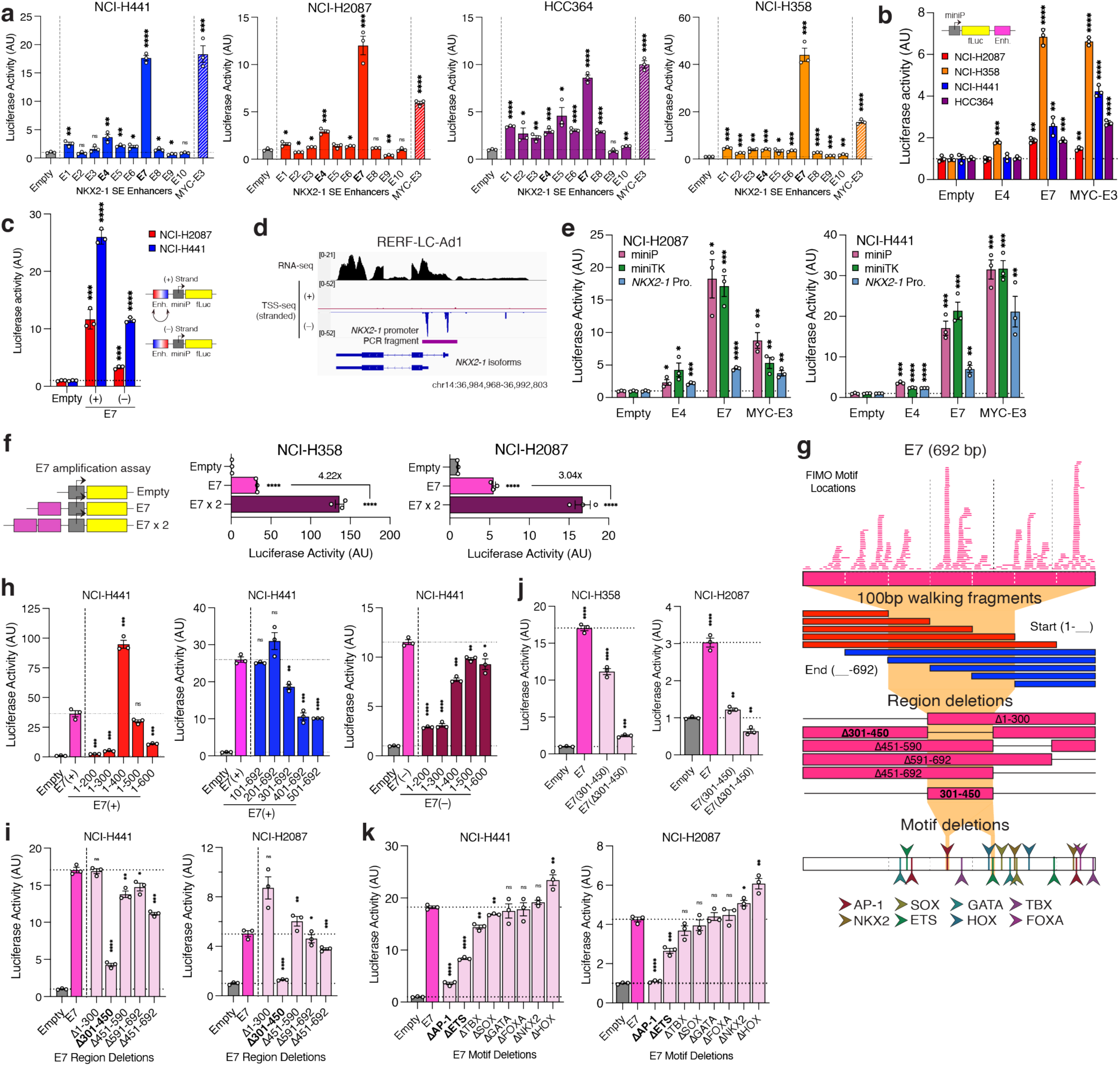
The NKX2-1 SE contains two enhancer elements that activate NKX2-1 promoter transcription, related to. **Figure 2** **(A)** Luciferase enhancer activity of 10 constituent enhancers of the *NKX2-1* SE, or the MYC-E3 LUAD enhancer^6^, upstream of a miniP luciferase reporter driving expression of firefly luciferase (miniP-fLuc), in 4 NKX2-1(+) LUAD cell lines. Consistently active enhancers (E4, E7) are indicated. E7 shows similar activity to the robust MYC-E3 LUAD enhancer we previously characterized^6^. **(B)** Enhancer activity of E4, E7, and MYC-E3 enhancers cloned downstream of miniP-fLuc in 4 NKX2-1(+) LUAD cell lines. E7 and MYC-E3 activate transcriptional at a distance in all 4 LUAD cell lines, E4 only shows activity in 1 cell line. **(C)** Luciferase enhancer activity of the E7 enhancer cloned upstream of the miniP-fLuc reporter, in (+) strand (by genome coordinates) and (–) strand/inverted orientations. The E7 enhancer shows directionally-biased activity, likely due to proximity of an active region at the 3’ end of the enhancer to miniP (30bp from MCS insertion site to start of miniP). However, both orientations show strong transcriptional activity by E7. **(D)** (top) Published RNA-seq^44^ and strand-specific TSS-seq^86^ at the *NKX2-1* gene locus in RERF-LC-Ad1 NKX2- 1(high) LUAD cells, showing usage of two TSS sites for *NKX2-1*. (bottom) Genome coordinates for the 1197bp *NKX2-1* promoter fragment cloned into pGL4.23 luciferase reporter construct. This fragment includes both isoform TSS sites, up to the translation start site of the short *NKX2-1* isoform, with a Kozak sequence for translation initiation of fLuc. **(E)** Luciferase enhancer activity of E4, E7, or MYC-E3 enhancers cloned upstream of miniP, miniTK, or the *NKX2-1* promoters in NCI-H441 or NCI-H2087 cells. E4 and E7 can activate all three promoters, but activation of the *NKX2-1* promoter is weaker than miniP or miniTK. However, the reduced activation is also seen with MYC-E3, suggesting this is due to either the larger enhancer-TSS distance for the *NKX2-1* promoter, or due to a weaker intrinsic activity of the promoter dampening its activation. **(F)** Luciferase enhancer activity of a duplicated E7 enhancer upstream of miniP promoter in NCI-H358 and NCI- H2087 cells shows >2x amplified activity as compared to a single E7 enhancer element. **(G)** (top) FIMO^88^ motif locations as identified in the E7 (692bp) sequence. (bottom) Schematic for E7 enhancer activity mapping, using walking 100bp fragments (red/blue), deletion fragments, and motif deletion assays of E7 transcriptional activity. **(H)** Luciferase enhancer activity of 100bp walking fragments of E7 normalized to the total activity of the full-length E7 sequence. Fragments were incorporated as (left) walking from the start (1-200/300/400/500/600/692bp, red) or (middle) walking from the end (101/201/301/401/501-692bp, blue) fragments. Due to hyperactivity of the 1-400bp forward fragment, forward fragments were incorporated in an inverted orientation ((–) strand, right) to account for possible transcriptional read-through from a transcription start site. Inverted forward fragments eliminate hyperactivity, pointing to a core 200-500bp region critical for activity. **(I)** Luciferase enhancer activity of E7 with deletion of 5 individual regions of transcription factor motif clustering, in NCI-H441 or NCI-H2087 cells. **(J)** Luciferase enhancer activity of E7(full length), E7(301-450), and E7(Δ301-450) in NCI-H2087 and NCI-H358 cells. Note that E7(301-450) shows stronger activity than E7(Δ301-450). **(K)** Luciferase enhancer activity of E7 with deletion of the individual binding motifs for 8 transcription factor families in NCI-H441 and NCI-H2087 cells. AP-1 and ETS motif deletions are required for E7 activity. **(A-C,E-F,H-K)** Two-tailed t-test, *p<0.05, **p<0.01, ***p<0.001, ****p<0.0001, for each condition compared to **(A-C,E-F,J)** empty vector control or **(H-I,K)** E7 wild-type control. Error bars = Mean±SEM, n=3 biological replicates, individual points labeled.

**Supplementary Figure 5.**
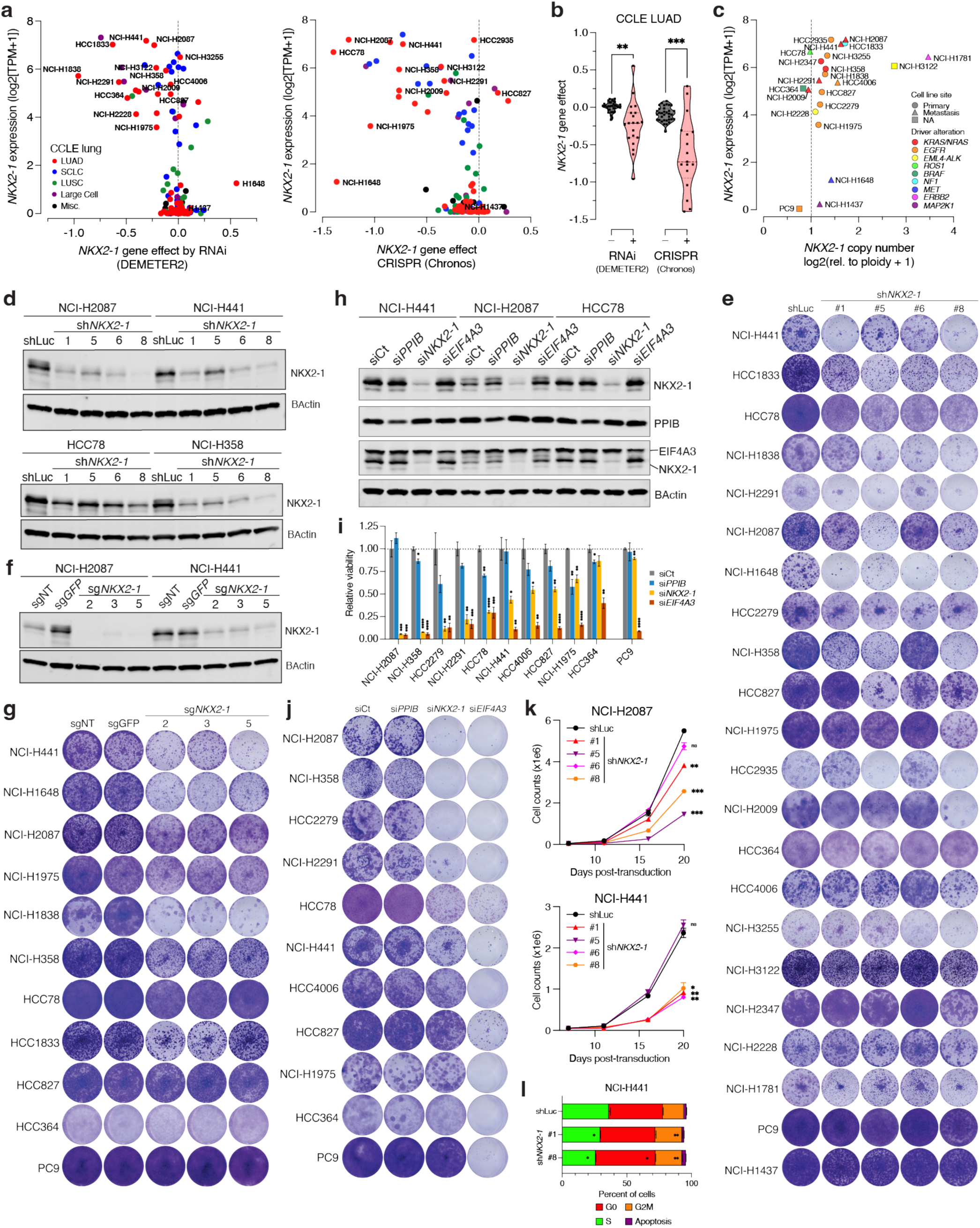
NKX2-1 is a dosage dependency across NKX2-1(+) LUAD cell lines, related to. **Figure 3** **(A)** Plot of *NKX2-1* dependency vs. expression in CCLE lung cell lines in genome-wide (left) RNAi and (right) CRISPR-Cas9 screens. Cell lines are colored by lineage subtype (LUAD: lung adenocarcinoma, SCLC: small cell lung cancer, LUSC: lung squamous cell carcinoma, Large Cell: large cell carcinoma, Misc.: NSCLC miscellaneous). Individual cell lines assayed labeled. **(B)** Truncated violin plot of *NKX2-1* dependency in NKX2-1(+) and NKX2-1(–) CCLE LUAD cell lines in RNAi and CRISPR-Cas9 genome-wide screens. Two-tailed t-test, **p<0.01, ***p<0.001. **(C)** Plot of *NKX2-1* copy number vs. RNA-seq expression in CCLE LUAD cell lines selected for *NKX2-1* dependency analysis (n=22). Cell lines are colored by putative oncogenic driver alterations (left) or primary or metastatic site-derived (right). **(D)** Immunoblot analysis of NKX2-1 levels in 4 NKX2-1(+) LUAD cell lines transduced with shRNAs targeting *NKX2-1* (#1,#5,#6,#8) or a luciferase-targeting control. **(E)** Representative images of clonogenic assays for *NKX2-1* dependency by shRNA in n=22 LUAD cell lines. Cells transduced with shRNAs targeting *NKX2-1* (#1,#5,#6,#8) or a luciferase-targeting control were plated for clonogenic assay, stained with crystal violet solution after expansion, and imaged. Images are representative of n=4 technical replicates per shRNA. **(F)** Immunoblot analysis of NKX2-1 levels in 2 NKX2-1(+) LUAD cell lines transduced with sgRNAs targeting *NKX2-1* (#2,#3,#5) or controls (sgNT, sg*GFP*). **(G)** Representative images of clonogenic assays for *NKX2-1* dependency by CRISPR-Cas9 in n=11 LUAD cell lines. Cells transduced with sgRNAs targeting *NKX2-1* (#2,#3,#5) or control sgRNAs (sgNT, sg*GFP*) were plated for clonogenic assay, stained with crystal violet solution after expansion, and imaged. Images are representative of n=4 technical replicates per sgRNA. **(H)** Immunoblot analysis of NKX2-1 levels in 3 NKX2-1(+) LUAD cell lines transfected with siRNAs targeting *NKX2-1*, *PPIB* (gene-targeting control), *EIF4A3* (pan-essential killing control), or a nontargeting control (Ct). **(I)** Relative viability by clonogenic assay in NKX2-1(+) (n=10) and NKX2-1(–) (n=1) LUAD cell lines upon siRNA-mediated knockdown of *NKX2-1*. n=4 technical replicates. Cells were plated for clonogenic assay and reverse transfected with siRNAs targeting *NKX2-1*, *PPIB*, *EIF4A3* or a nontargeting siCt at 10 nM for 48h, stained with crystal violet solution after expansion, and quantified. n=3 biological replicates, Two-tailed t-test against control, *p<0.05, **p<0.01, ***p<0.001, ****p<0.0001. Error bars = Mean±SEM **(J)** Representative images of clonogenic assays for *NKX2-1* dependency by siRNA as in (I). Images are representative of n=3 biological replicates per siRNA. **(K)** Proliferation curves of NCI-H2087 or NCI-H441 cells transduced with shRNAs targeting *NKX2-1*, or shLuc control. Days post-lentiviral infection are indicated, cells were plated at 50K/well at day 7 and counted in technical duplicate at days 11/16/20 post-transduction. n=2 technical replicates per shRNA, error bars = Mean±SEM. Two-tailed t-test against shLuc, *p<0.05, **p<0.01, ***p<0.001, ****p<0.0001. **(L)** Cell cycle analysis of NCI-H441 cells transduced with shRNAs targeting *NKX2-1* (#1,#8) or shLuc control by flow cytometry. Two-tailed t-test against shLuc control, *p<0.05, **p<0.01, ***p<0.001, ****p<0.0001. n=2 biological replicates.

**Supplementary Figure 6.**
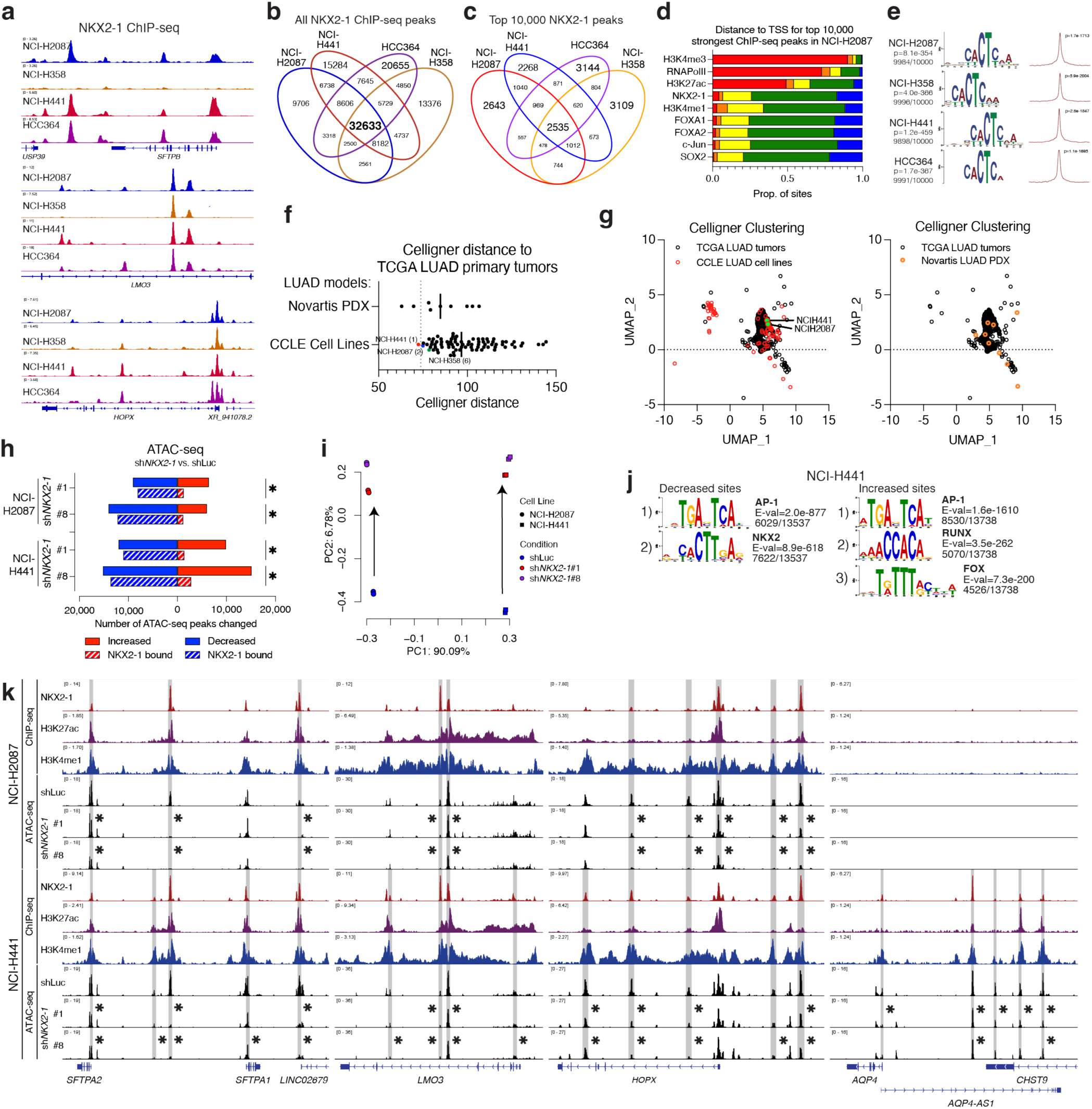
NKX2-1 binds to and remodels lineage enhancer accessibility in LUAD, related to. **Figure 4** **(A)** ChIP-seq occupancy tracks for NKX2-1 binding at the *SFTPB, HOPX*, and *LMO3* loci in 4 NKX2-1(+) LUAD cell lines. **(B-C)** Venn diagram of **(B)** all NKX2-1 ChIP-seq binding sites or **(C)** the top 10,000 strongest NKX2-1 ChIP-seq binding sites by signal in 4 NKX2-1(+) LUAD cell lines. **(D)** Distribution from TSS in the genome of the 10,000 strongest ChIP-seq binding sites for varied histone marks, regulators, and transcription factors in NCI-H2087 cells. NKX2-1 occupies a distal binding profile similar to other TFs profiled. **(E)** Motif discovery at the top 10,000 NKX2-1 ChIP-seq peaks in 4 NKX2-1(+) LUAD cell lines discovers and enriches the NKX2-1 binding motif. (left) p-value and number of matching sites for (center) the top motif discovered by STREME^84^, and (right) Centrimo^85^ central enrichment plot and p-value for the identified NKX2-1 motif. **(F)** Celligner distance to primary TCGA LUAD tumors for Novartis LUAD PDX and CCLE LUAD cell line models. NCI-H441 and NCI-H2087 are labeled. **(G)** Celligner UMAP plot of primary TCGA LUAD tumors with (left) CCLE LUAD cell lines and (right) Novartis LUAD PDX models. **(H)** Total number of ATAC-seq peaks with increased (red) or decreased (blue) accessibility upon *NKX2-1* knockdown by either sh*NKX2-1*#1 or *shNKX2-1*#8 in NCI-H2087 or NCI-H441 cells, as compared to shLuc control. Sites overlapping with NKX2-1 ChIP-seq peaks (hashed) are indicated, Fisher exact test *p<2.2e-16. **(I)** Principal component analysis of normalized ATAC-seq chromatin accessibility in NCI-H2087 and NCI-H441 upon knockdown of *NKX2-1*. shNKX2-1#1 and shNKX2-1(8) induce concordant changes across cell lines. **(J)** Motifs discovered by STREME^84^ at sites of (top) decreased or (bottom) increased ATAC-seq accessibility sites upon NKX2-1 knockdown in NCI-H441 cells. E-value and number of sites with motif identified are indicated. **(K)** NKX2-1, H3K27ac, and H3K4me1 ChIP-seq and ATAC-seq in shLuc, sh*NKX2-1*#1 and sh*NKX2-1*#8 conditions in (top) NCI-H2087 and (bottom) NCI-H441 cells. * indicates significantly downregulated ATAC-seq peak by DESeq2.

**Supplementary Figure 7.**
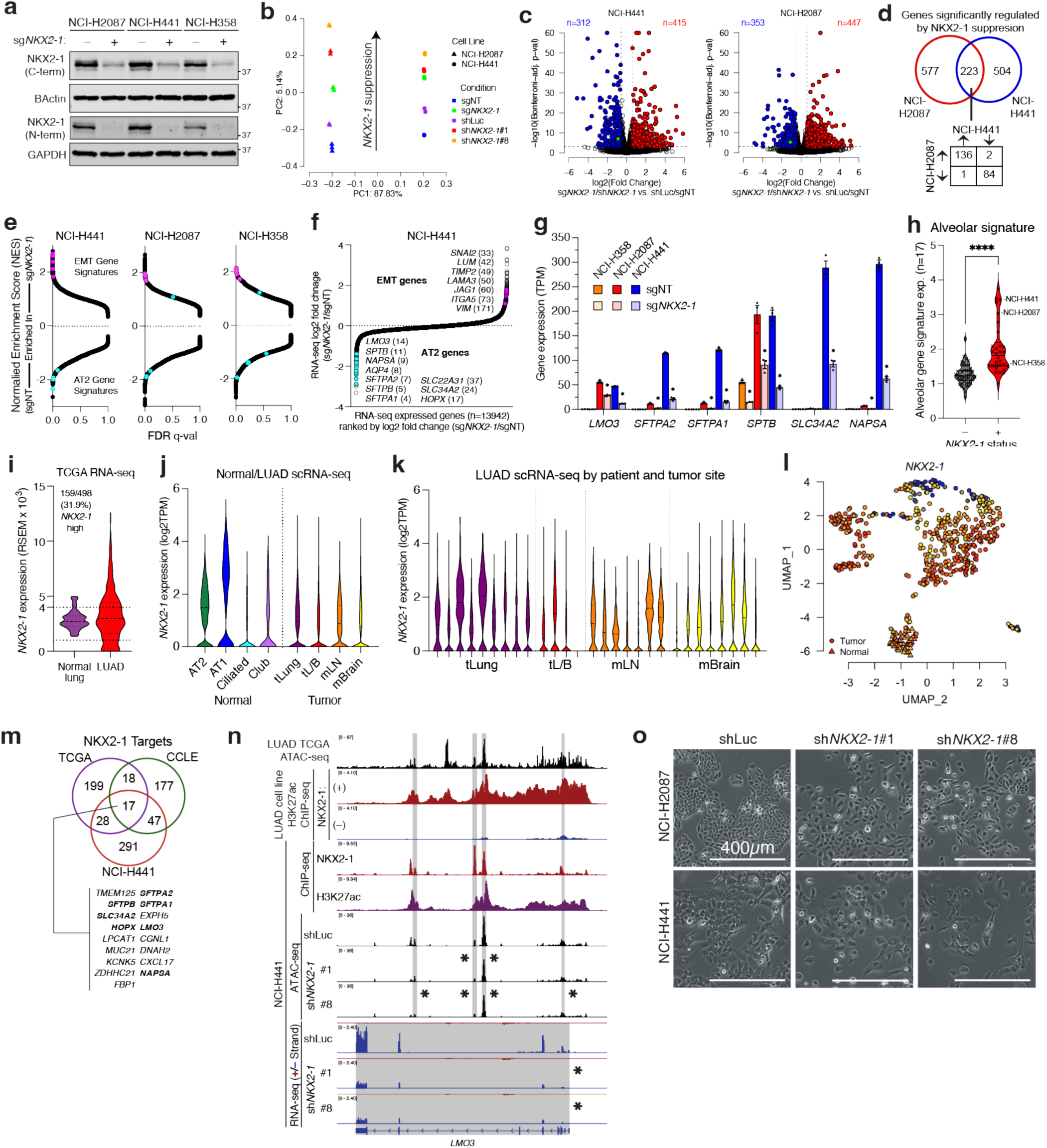
NKX2-1 defines a lineage differentiation state and regulates epithelial-mesenchymal transition in LUAD, related to. **Figure 4** (A) Immunoblot analysis of NKX2-1 levels in NCI-H2087, NCI-H441, and NCI-H358 cells transduced with sgRNAs targeting *NKX2-1* or a non-targeting control. (B) Principal component analysis of RNA-seq in NCI-H2087 and NCI-H441 upon knockdown of *NKX2-1* by sgRNA or shRNA with shLuc/sgNT controls. *NKX2-1* knockdown induces concordant changes across cell lines. (C) Plot of log2FC vs. Bonferroni adjusted p-value in RNA-seq expression upon *NKX2-1* knockdown (shNKX2-1 and sgNKX2-1) vs. controls (shLuc/sgNT) by DESeq2. Genes significantly upregulated (red) or downregulated (blue) are indicated, number of genes is labeled. *NKX2-1* is labeled in green. (D) Venn diagram of significantly changed genes identified in (J) upon *NKX2-1* knockdown by RNA-seq in NCI- H441 and NCI-H2087 LUAD cell lines. 223 genes are significantly changed in both lines—of these, 220/223 (99%) are regulated concordantly by NKX2-1. (E) Plot of FDR q-value vs. normalized enrichment score (NES) for ranked gene set enrichment analysis (GSEA) across 5175 MSigDB gene sets, upon NKX2-1 knockdown (sg*NKX2-1*/sgNT) in 3 NKX2-1(+) cell lines. Interferon (orange) and MYC gene sets (purple) are highlighted. (F) Ranked log2 fold-change in RNA-seq expression change upon NKX2-1 knockdown (vs. sgNT control) for all expressed genes (N=13942) in NCI-H441 cells. EMT (pink) and AT2 (cyan) genes are labeled and their rank stated in parenthesis. (G) Bar graph of normalized RNA-seq expression for AT2 genes regulated by NKX2-1, in 3 NKX2-1(+) LUAD cell lines, in transcripts per million (TPM). Error bars = mean±SEM, n=3 biological replicates, individual values labeled. * significant by DESeq2 (FC ≥ 1.5, Bonferroni adj. p-value ≤ 1e-3). (H) Expression of an alveolar signature (n=17) by RNA-seq is significantly higher in NKX2-1(+) LUAD cell lines than NKX2-1(–) cell lines. Two-tailed t-test, ****p<0.0001. (I) Violin plot of *NKX2-1* expression by RNA-seq in TCGA normal lung and LUAD tumors. *NKX2-1* is expressed higher than normal lung (RSEM ≥ 4000) in 159/498 (31.9%) of LUAD tumors. (J) Violin plot of expression of *NKX2-1* by scRNA-seq^75^ in malignant LUAD cells from early stage primary tumors (tLung), advanced primary tumors (tL/B), or metastatic tumors from lymph node (mLN) or brain (mBrain) sites, as well as normal lung cell types (AT1, AT2, ciliated, club). (K) Violin plot of expression of *NKX2-1* by scRNA-seq^75^ in malignant LUAD cells from individual tumor samples as in (C). *NKX2-1* expression is observed in the majority of cells from all tumor sites and stages. (L) UMAP clustering of RNA-seq gene expression from n=576 TCGA normal lung (triangle) and primary LUAD tumors (circle). Samples are colored by z-score normalized *NKX2-1* expression. Most primary LUAD tumors are NKX2-1(+), with a distinct population of tumors harboring low *NKX2-1*. (M) Venn diagram of genes in *NKX2-1* gene signatures identified in CCLE and TCGA by *NKX2-1* expression correlation, and in NCI-H441 cells by downregulation upon *NKX2-1* depletion. 17 shared genes are listed; alveolar markers are bolded. (N) (top) TCGA LUAD ATAC-seq and H3K27ac ChIP-seq from NKX2-1(+) and NKX2-1(–) LUAD cell lines at the *SFTPA1/2* locus (middle) NKX2-1 and H3K27ac ChIP-seq from NCI-H441 cells (bottom) ATAC-seq and strand-specific RNA-seq from NCI-H441 cells in shLuc, sh*NKX2-1*#1 and sh*NKX2-1*#8 conditions. * indicates significantly downregulated ATAC-seq peak or RNA-seq gene by DESeq2. (O) Light microscopy images of NCI-H2087 and NCI-H441 cells transduced with shRNAs targeting *NKX2-1*, or shLuc control. LUAD cell lines lose cell-cell adherence and exhibit distinct morphology upon suppression of *NKX2-1*. Scale bar = 400 µm.

**Supplementary Figure 8.**
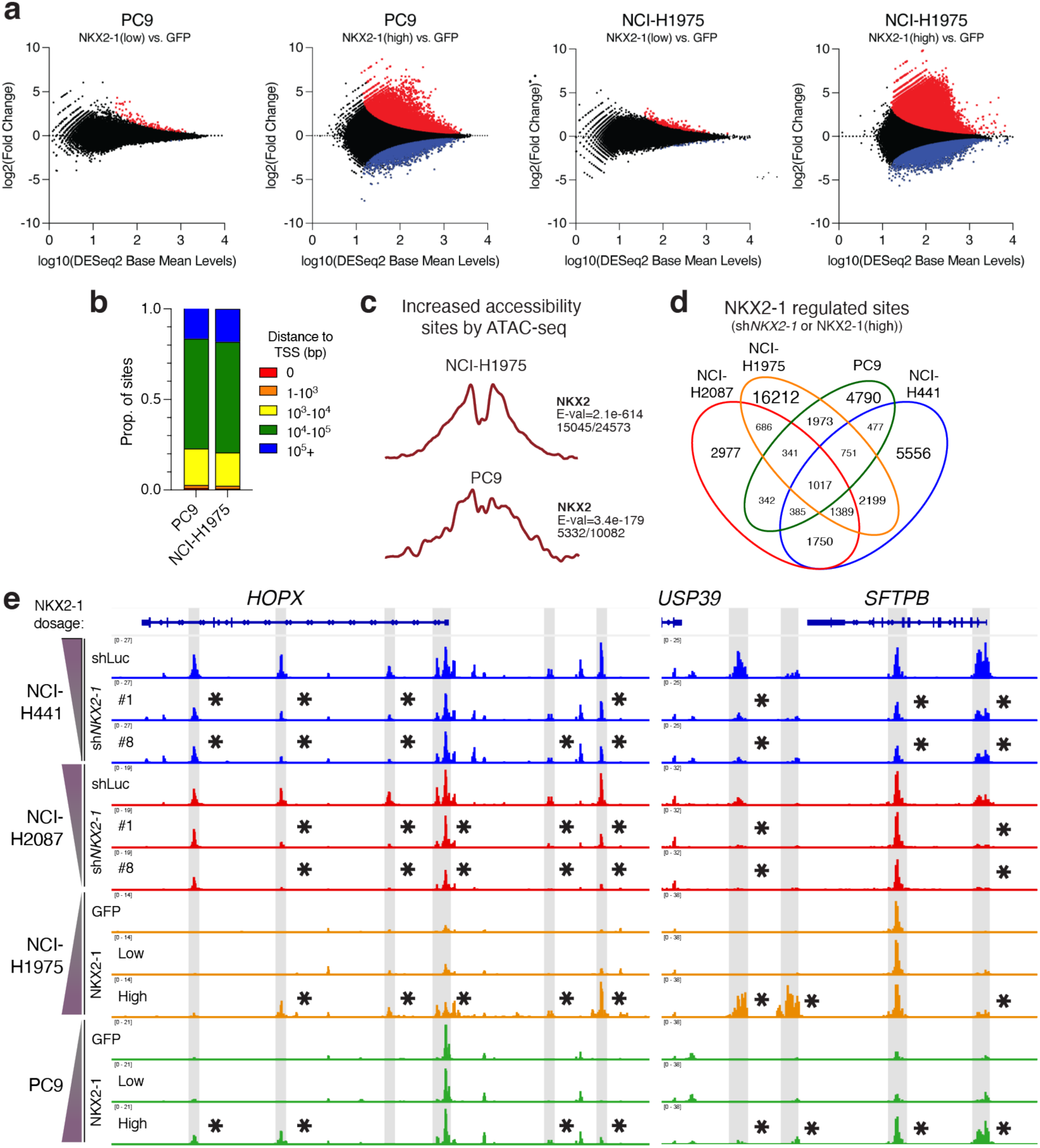
NKX2-1 remodeling of enhancer accessibility is dose-thresholded, related to. **Figure 5** **(A)** MA plot of base mean ATAC-seq signal vs. log2(fold change) for overexpression of NKX2-1 to low (hPGK) or high (EF1α) exogenous levels, as compared to GFP control, in PC9 and NCI-H1975 cells. Significantly increased (red) or decreased (blue) accessibility sites are indicated. **(B)** Distance to transcription start site (TSS) for peaks with increased ATAC-seq accessibility upon NKX2-1(high) overexpression in PC9 or NCI-H1975 cells. **(C)** CENTRIMO^85^ central motif enrichment of the NKX2-1 motif at sites of increased ATAC-seq accessibility upon NKX2-1(high) overexpression in PC9 or NCI-H1975 cells. Central enrichment plot, E-value, and number of sites with matching motif are indicated. **(D)** Venn diagram of ATAC-seq peaks with decreased accessibility upon *NKX2-1* knockdown in NCI-H2087 (red) or NCI-H441 (blue) cells, as well as sites of increased accessibility upon NKX2-1(high) overexpression in NCI- H1975 (NKX2-1 low, orange) and PC9 (NKX2-1 negative, green). **(E)** Normalized ATAC-seq genome accessibility tracks at the *HOPX, SFTPB*, and *SPTB* loci upon *NKX2-1* modulation. * indicates sites significantly decreased (NCI-H2087/NCI-H441) or increased (NCI-H1975/PC9) with *NKX2-1* modulation by DESeq2 (FC ≥ 1.25, Bonferroni adj. p-value ≤ 1e-3).

**Supplementary Figure 9.**
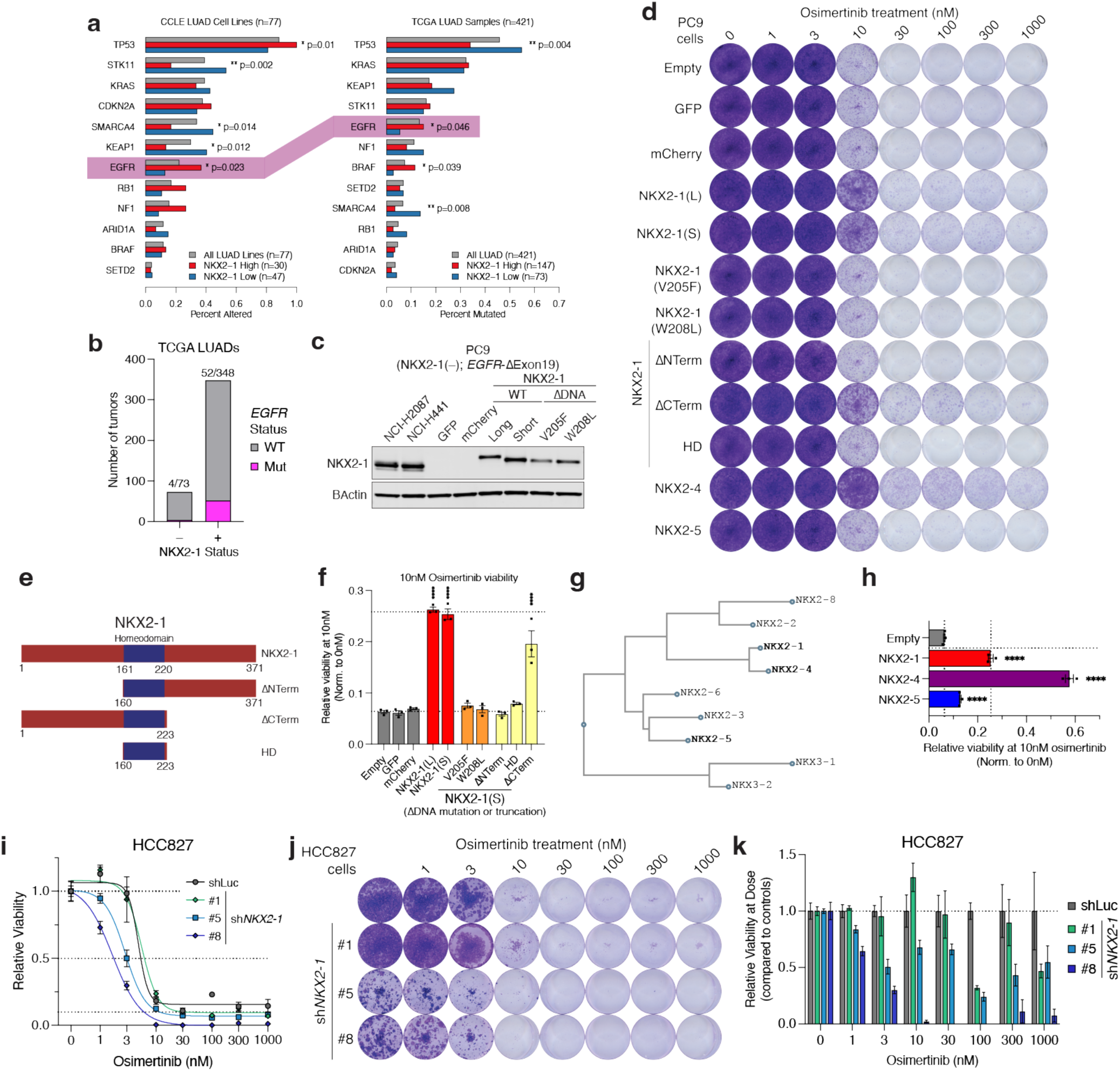
NKX2-1 drives an EGFR TKI persistent state in EGFR-mutant LUAD cells, related to Figure 6. **(A)** Mutation rates for top driver alterations in (left) CCLE LUAD cell lines (n=77) or (right) TCGA LUAD primary tumors (n=421) in NKX2-1(low) or NKX2-1(high) samples or overall mutation rate. *EGFR* mutations are highlighted in yellow. *EGFR* mutations are significantly depleted in NKX2-1(low) cell lines and tumors as compared to NKX2-1 competent cells. Significance by Fisher exact test. **(B)** Number of *EGFR* mutations in TCGA LUAD tumors by *NKX2-1* expression. EGFR mutations are largely exclusive of *NKX2-1* expression loss in LUAD. **(C)** Immunoblot analysis of NKX2-1 levels in PC9 cells in the indicated conditions, with NCI-H2087 and NCI-H441 NKX2-1(high) endogenous controls. **(D)** Crystal violet staining of clonogenic assays as indicated. NKX2-1 dosage dictates increased survival at both cytostatic (10nM) and cytotoxic (≥30nM) osimertinib dosage. DNA binding mutations fully suppress NKX2-1 regulation of TKI persistence, as does deletion of the N-terminus of NKX2-1. **(E)** Schematic of domain truncation variants of *NKX2-1*. **(F)** Relative viability of PC9 cells in the indicated conditions at 10nM osimertinib treatment, Two-tailed t-test vs controls, ****p<0.0001. **(G)** Phylogeny of NKX2/3 family transcription factors from human amino acid sequence, NKX2-1, NKX2-4, and NKX2-5 are indicated. **(H)** Relative viability of PC9 cells in the indicated conditions at 10nM osimertinib treatment, Two-tailed t-test vs controls, ****p<0.0001. **(I)** Dose response curve of HCC827 (*EGFR*-Exon19Δ) upon osimertinib treatment for 10 days, in shLuc and sh*NKX2-1* (#1, #5, #8) conditions. n=3 biological replicates, error bars = mean±SEM. **(J)** Crystal violet staining of clonogenic assays as in **(I)** for HCC827 cells. NKX2-1 strong knockdown by #5 and #8 drives increased sensitivity to osimertinib and loss of persistent cells at higher doses. **(K)** Relative viability of HCC827 cells at each Osimertinib dosage as in **(I)**, normalized to shLuc viability, for *NKX2-1* knockdown. sh*NKX2-1* cells exhibit increased viability defects relative to control cells.

**Supplementary Figure 10.**
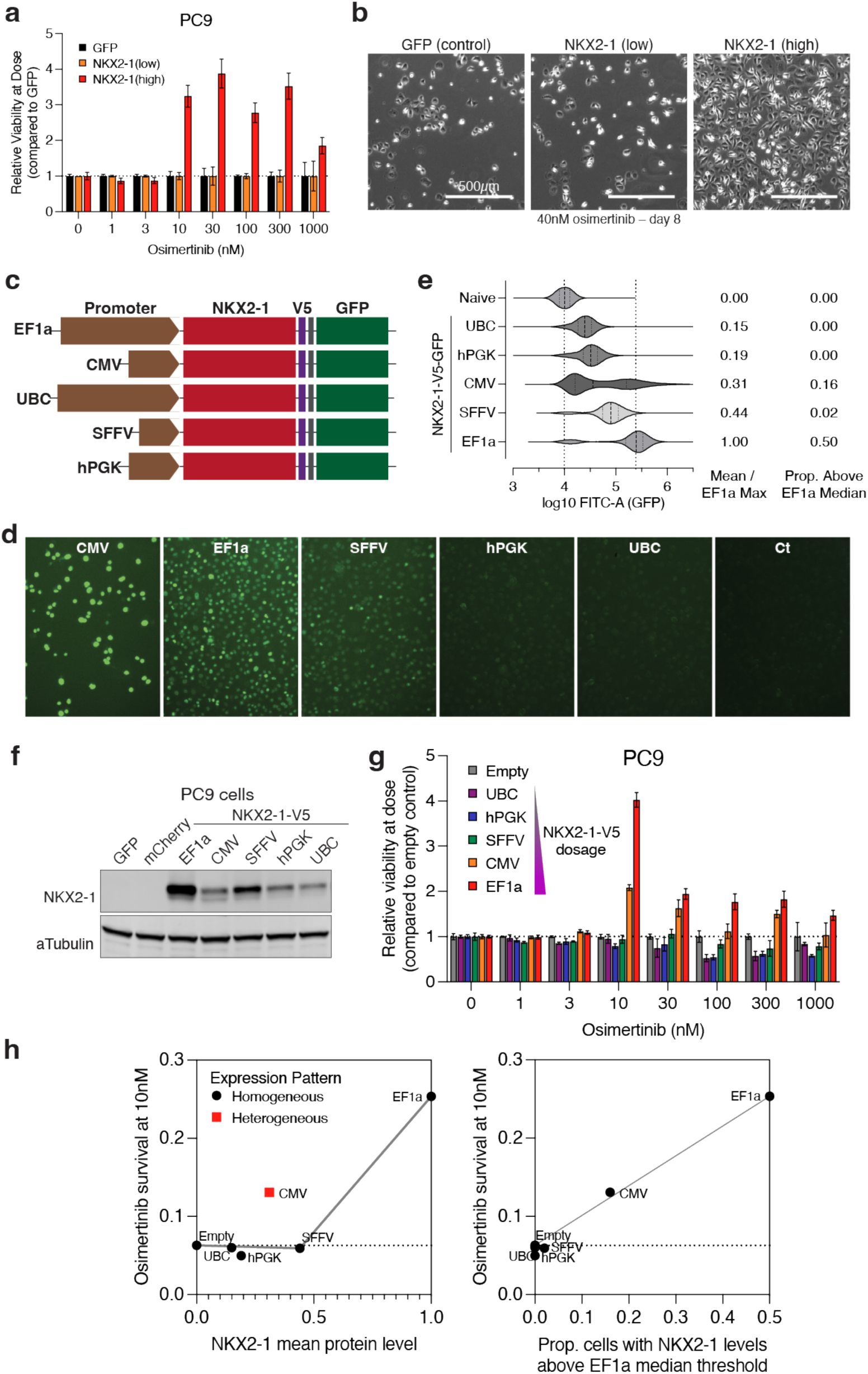
NKX2-1 control of EGFR TKI persistence is dose-thresholded, related to Figure 6. (A) Relative viability of PC9 cells at each osimertinib dose, normalized to GFP viability, for *NKX2-1* overexpression. NKX2-1(high) cells exhibit increased persistence relative to control cells. n=6 biological replicates, error bars = mean±SEM. (B) Light microscopy of PC9 cells at 40nM osimertinib for 8 days. NKX2-1(high) cells are able to proliferate and expand at cytotoxic osimertinib treatment. (C) Schematic for expression of *NKX2-1-V5-GFP* under the indicated promoters. (D) Light microscopy of PC9 cells expressing lentivirally-transduced NKX2-1-V5-GFP in the indicated conditions, or naïve control cells. (E) Violin plots of GFP (FITC-A) signal by flow cytometry, as in Fig. 6g. (F) Immunoblot of PC9 cells expressing NKX2-1-V5 on the indicated promoters, as in Fig. 6h. (G) Relative viability of PC9 cells at each osimertinib dose, normalized to GFP viability, for *NKX2-1* overexpression in the indicated promoters. n=3 biological replicates, error bars = mean±SEM. (H) (left) NKX2-1 mean expression level by flow cytometry vs. relative osimertinib survival at 10nM in clonogenic assays. (right) Proportion of cells with NKX2-1 expression levels above a dose threshold of the EF1a median expression vs. relative osimertinib survival at 10nM in clonogenic assays. NKX2-1 expression above this dose threshold precisely correlates with osimertinib survival observed.

## SUPPLEMENTARY TABLES

Supplementary Table 1 – Antibodies Used for ChIP-seq and Western Blotting

Supplementary Table 2 – Oligonucleotide Sequences

Supplementary Table 3 – Genome Coordinates

Supplementary Table 4 – Synthesized Enhancer Sequences for Luciferase Assay

Supplementary Table 5 – sgRNA Sequences

Supplementary Table 6 – shRNA Sequences and Vectors

Supplementary Table 7 – siRNA Sequences

Supplementary Table 8 – ChIP-seq Data Statistics

Supplementary Table 9 – ATAC-seq Data Statistics

Supplementary Table 10 – RNA-seq Data Statistics

